## Supplementary Material for "Real-time transcriptomic profiling in distinct experimental conditions"

**Supplementary Material 1. Step-by-step guideline to NanopoReaTA.** *(For additional information, please refer to Wierczeiko et al. 2023 or visit [https://github.com/AnWiercze/NanopoReaTA](https://github.com/AnWiercze/NanopoReaTA)).*

1) ***Run docker***

```
docker run -it -p 8080:8080 -v /:/NanopoReaTA_linux_docker
stegiopast/nanoporeata:references
```

2) ***Open in new window***

[http://0.0.0.0:8080](http://0.0.0.0:8080)

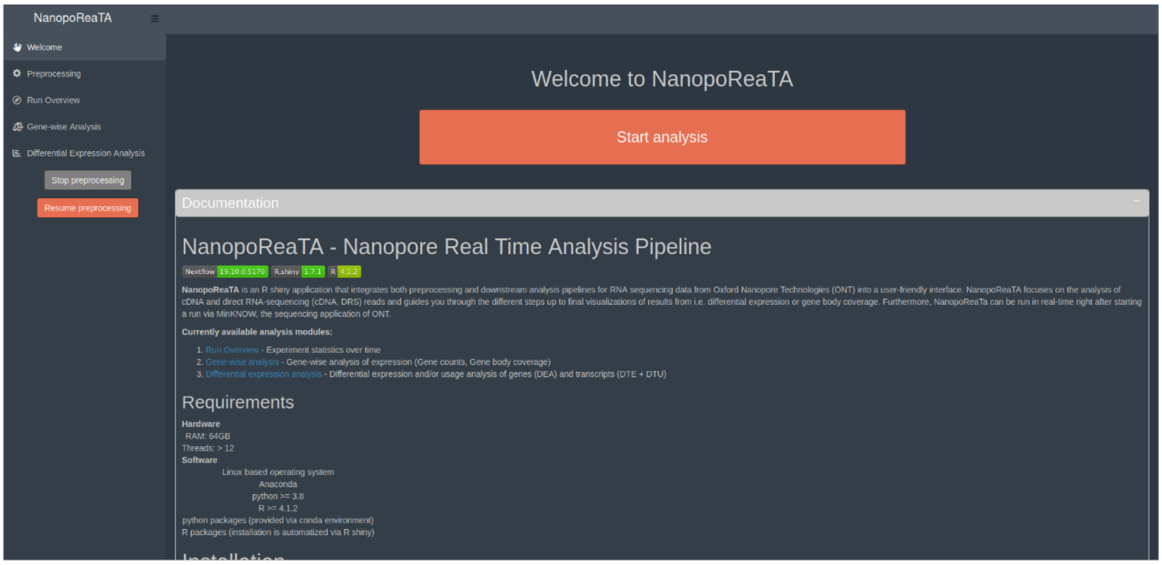


3) ***Metadata creator***

- insert in Samples the barcodes used with nanopore (e.g. barcode1, barcode2, etc..)
- insert in Condition the name of tested condition (e.g. Condition A, Condition B)
(Not recommended to insert "_" in the condition names)
- insert in Replicate the replicates names (e.g. R1)
- Download the metadata

- Users can prepare in advance metadata template from (*example_metadata.txt*):
[https://github.com/AnWiercze/NanopoReaTA/tree/master/example_conf_files](https://github.com/AnWiercze/NanopoReaTA/tree/master/example_conf_files)

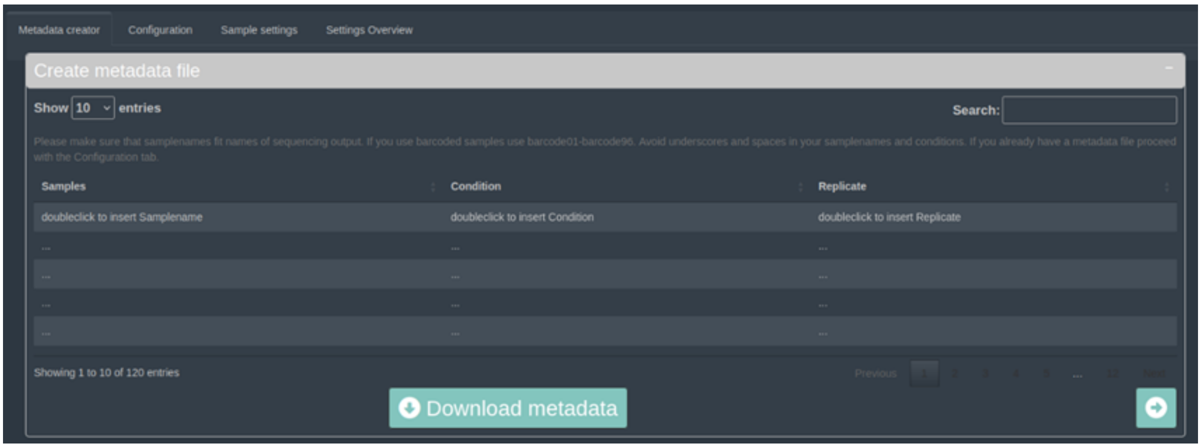

4) ***Creation of a configuration file***

- Download the configuration template from (*example_config.txt*):
https://github.com/AnWiercze/NanopoReaTA/tree/master/example_conf_files
- insert in Condition the name of tested condition (e.g. Condition A, Condition B) (No recommended _ in the condition names)
- insert in Replicate the replicates names (e.g. R1)
- Download the metadata

```
threads: 24 → - Number of threads to use
barcoded: 1 → - Barcoded samples. 0 – no; 1 - Yes
DRS: 0 → - Direct RNA sequencing. 0 – no; 1 - Yes
metadata: /NanopoReaTA_linux_docker/home/XXX/XXX/metadata_RiboM_RiboP.tsv → - Path to the metadata file (.tsv)
general_folder: /NanopoReaTA_linux_docker/home/XXX/XXX/XXX/ → - Path to the sequencing folder (MinKnow Output)
genome_fasta: /Reference_data/Human_reference_data/GRCh38.primary_assembly.genome.fa → - *Path to refence genome (.fasta)
transcriptome_fasta: /Reference_data/Human_reference_data/gencode.v43.transcripts.fa → - *Path to reference trascriptome (.fasta)
genome_gtf: /Reference_data/Human_reference_data/gencode.v43.primary_assembly.annotation.gtf - *Path to Gtf file (.gtf)
bed_file: /Reference_data/Human_reference_data/hg38_GENCODE_V42_Comprehensive.bed - *Path to Bed file (.bed)
run_dir: /NanopoReaTA_linux_docker/home/XXX/XXX/ - Path to NanopoReaTA output folder
```

*With the docker image tag "references", all human and mouse reference files needed for NanopoReaTA will be automatically downloaded from GENCODE (~36 GB) and saved in root.

5) ***Loading the configuration file***

- open the configuration tab and press on "Select file" to load the configuration file.

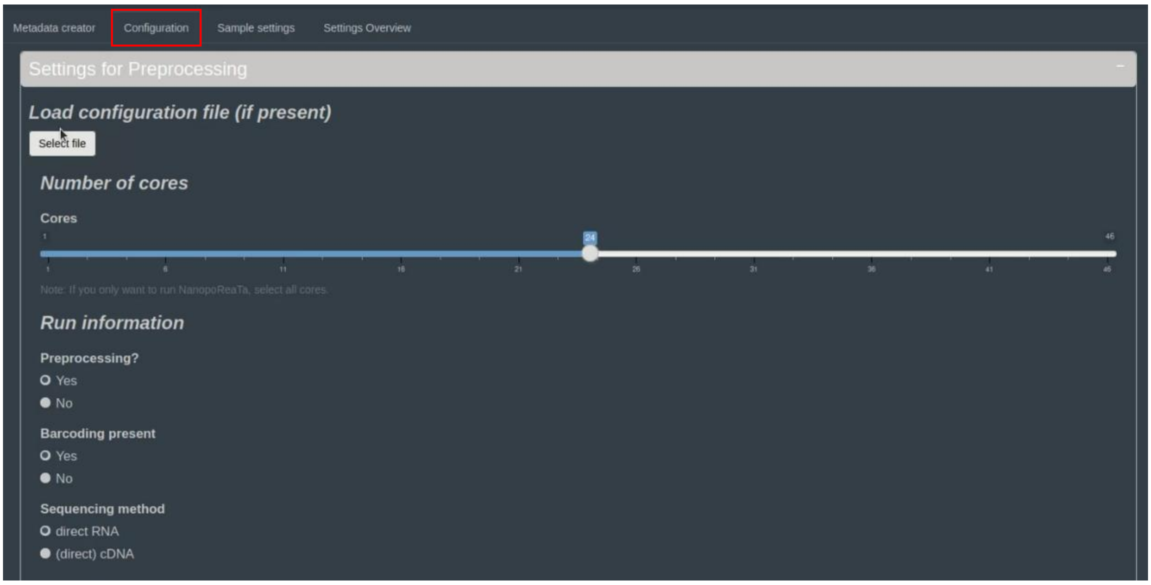


- Select the configuration file (.yaml)

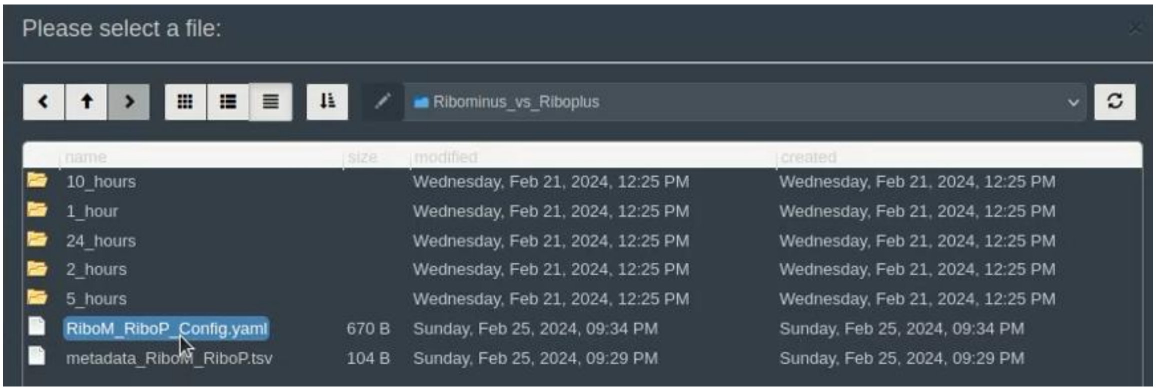

- Selected files and folder output should upload to the appropriate section
- Note: if not all of the files are uploaded, you can insert the missing file manually.

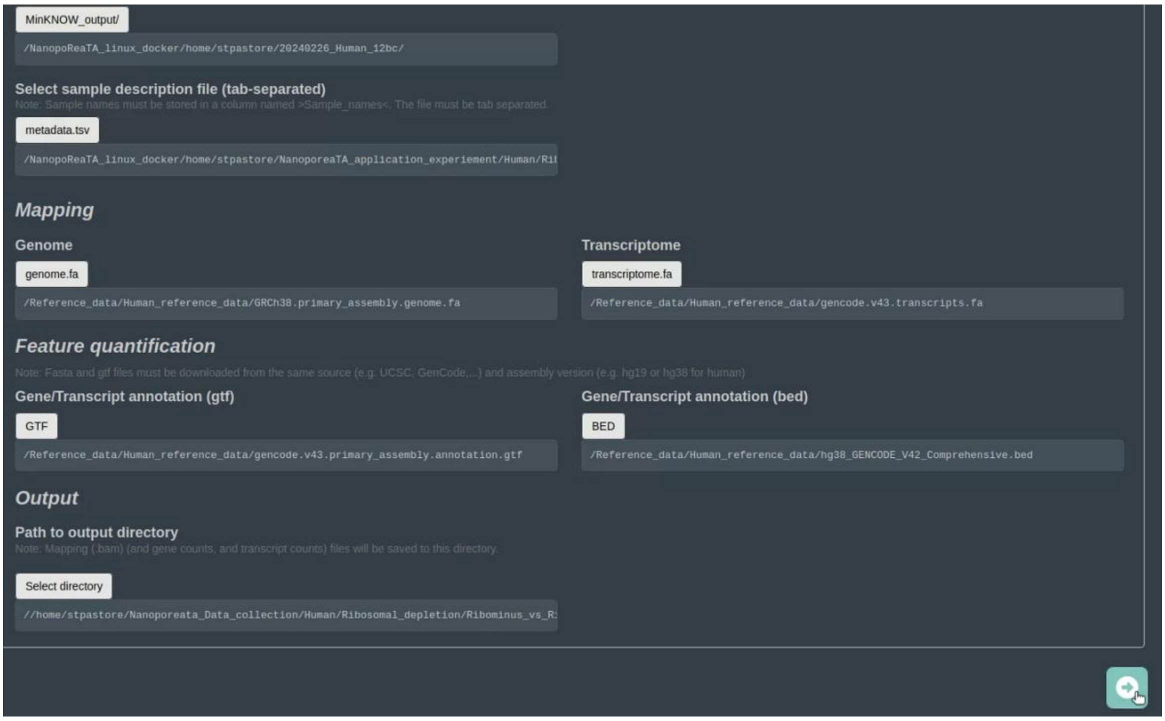


- Following configuration upload, check in “Sample settings” that all barcodes, conditions and replicates are inserted properly
- Users can select colors for the specific condition which will be shown in the visualized data analysis

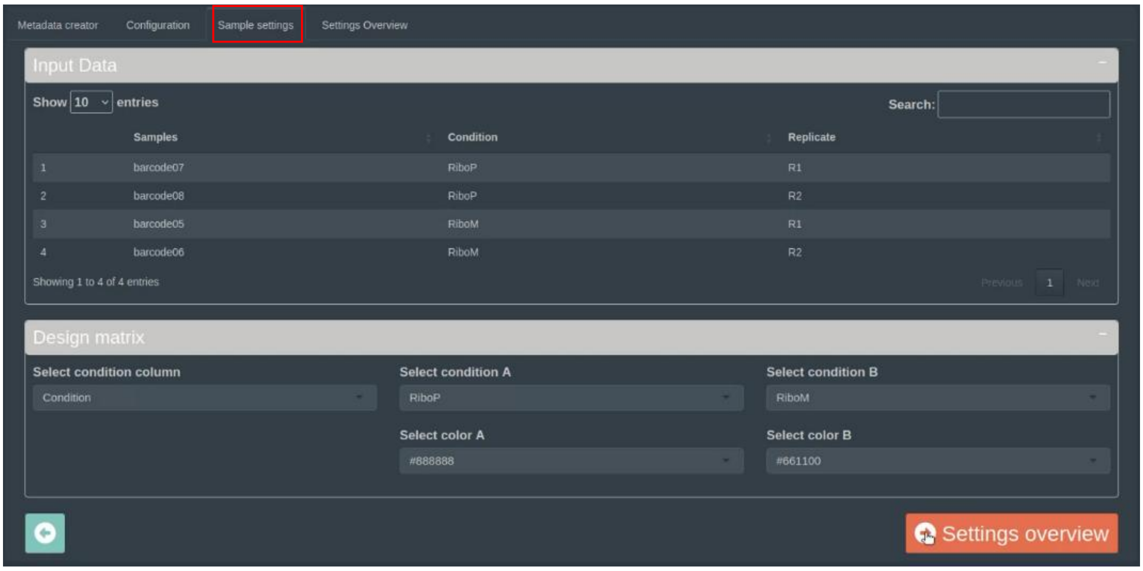

- In “Settings Overview”, check that all the details are inserted properly and press “Start” real-time transcriptomic analysis

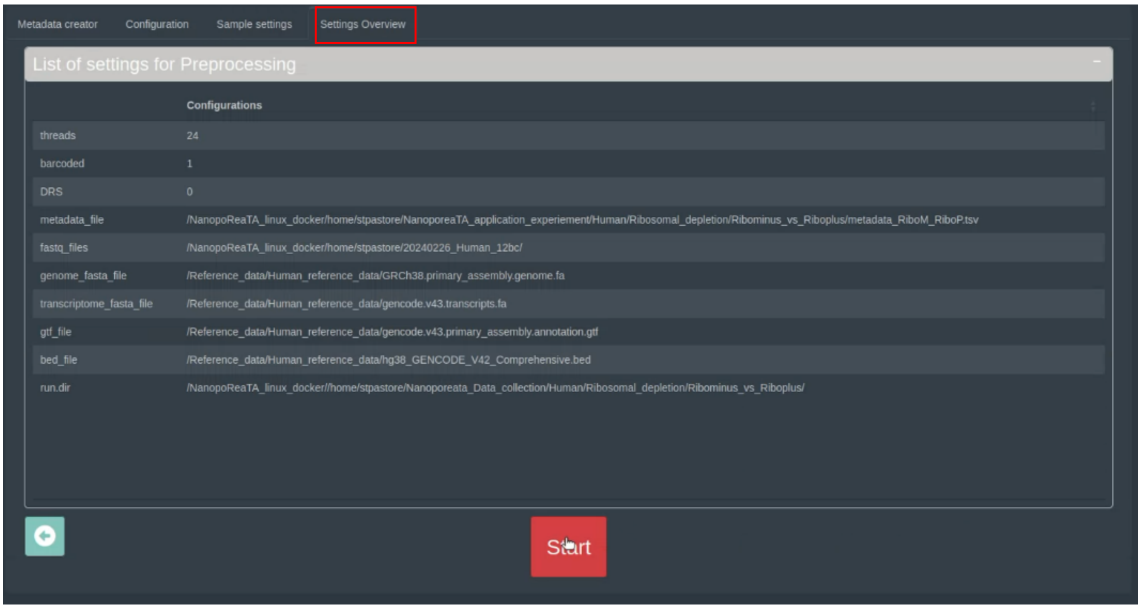


**6)** ***Run Overview***

- In “Run Overview”, users can check general information about the analyzed samples/conditions.
- At the top of the page, users can see a table with barcoded samples, mapped reads, gene counts and transcripts. The table can be copied or exported as *csv* or *txt* files.

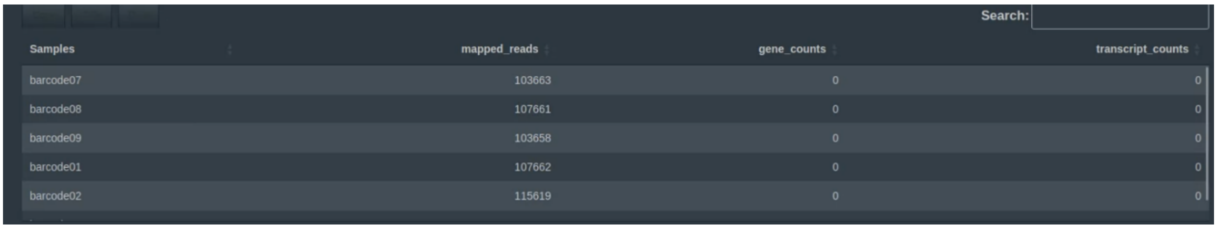

| Samples | mapped_reads | gene_counts | transcript_counts |
|---|---|---|---|
| barcode07 | 103663 | 0 | 0 |
| barcode08 | 107661 | 0 | 0 |
| barcode09 | 103658 | 0 | 0 |
| barcode01 | 107662 | 0 | 0 |
| barcode02 | 115619 | 0 | 0 |

- In the “Read length distribution” tab, users can have an overview of the read length overview per barcoded samples and conditions.
- Figures can be exported as png figure by clicking the “download” button.

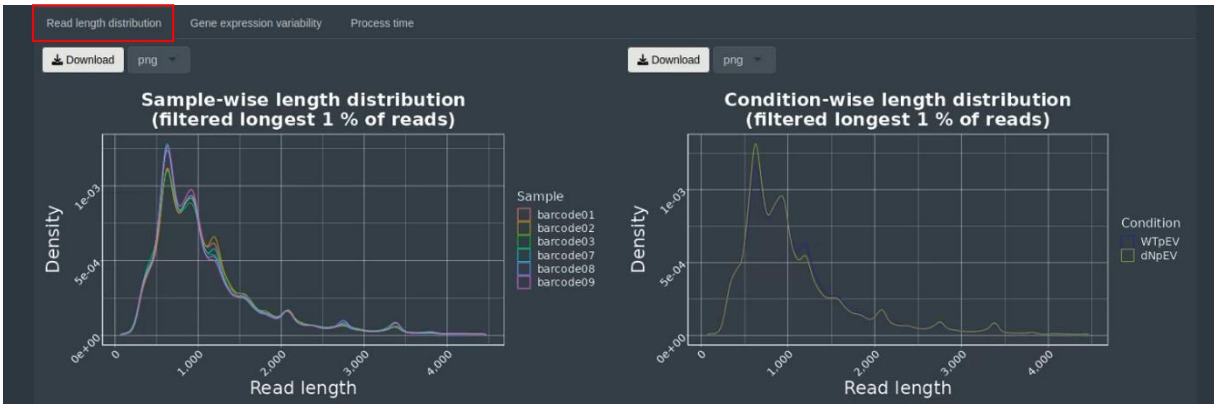

- In the “Gene expression variability” tab, users can have an overview number of genes detected and changes in gene composition plot, per sample and per condition
- Figures can be exported as png figure by clicking the “download” button.

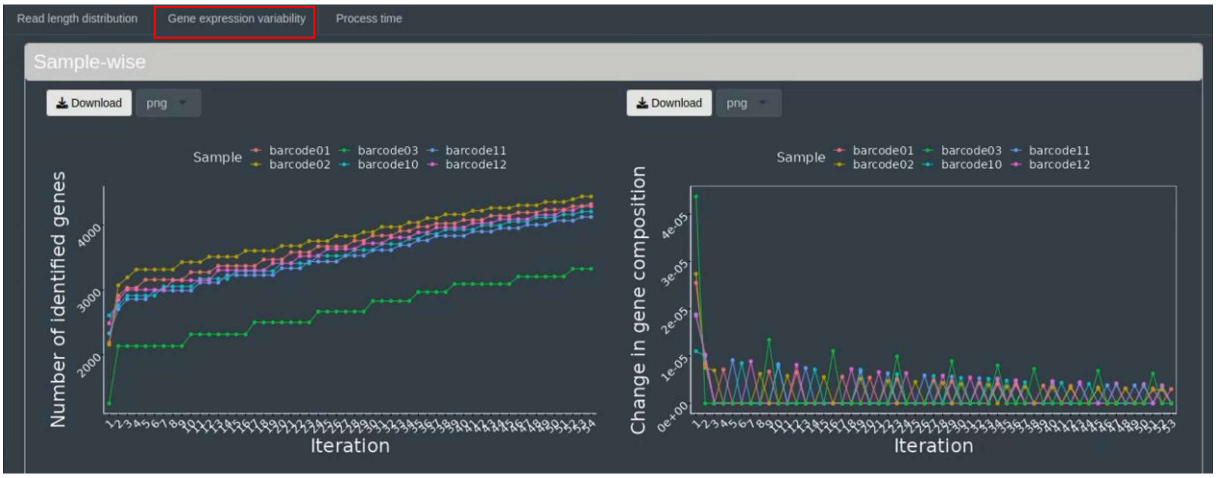


- In the “process time” tab, users can have an overview of each tool’s processing times per iteration.
- Figure can be exported as png figure by clicking the “download” button.

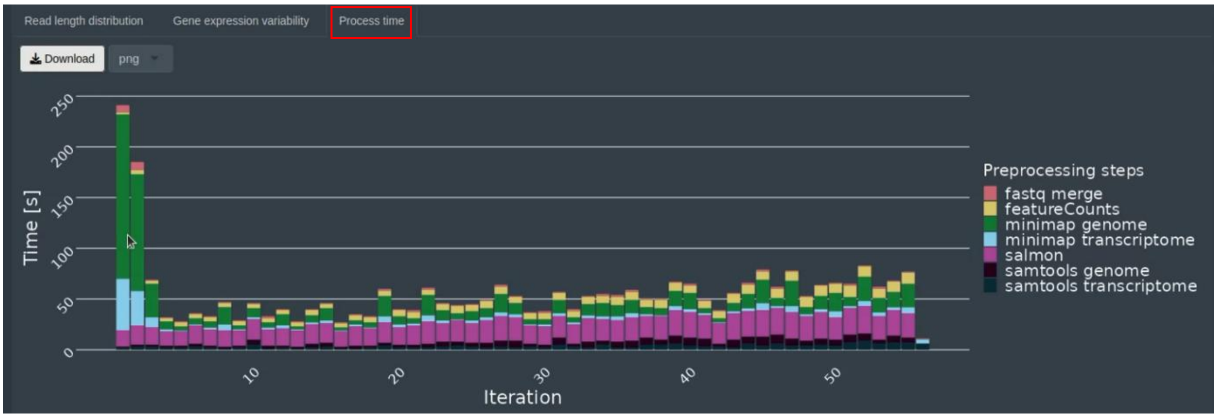


### 7) ***Differential expression analysis***

- In the "differential expression analysis" tab, users can choose the type of analysis they wish to perform, including "Gene expression analysis," "Transcript expression analysis," and "Transcript usage analysis".

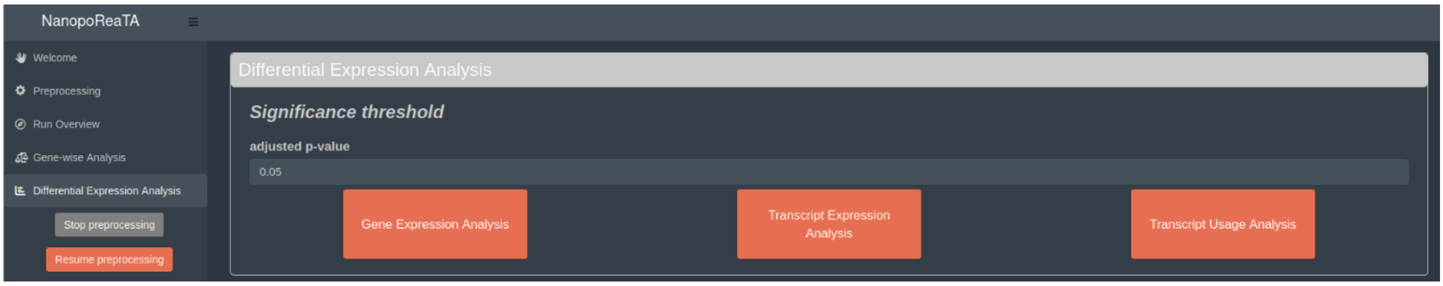

- Before activating any of the tools, it is important to follow these steps:

***Step 1*:** Press the "Stop preprocessing" button.
***Step 2*:** The button will turn grey, meaning the preprocessing of the sequenced files has stopped.
***Step 3*:** You can observe the last analysis steps being performed in the "process overview" tab.
***Step 4*:** Once the last analysis tool finalizes its task, it will be displayed in the "process overview" tab, and the "Resume preprocessing" button will become available.
***At this point, the activation of the "Differential Expression Analysis" tools is possible.***
**Step 5:** To resume preprocessing additional files, press the "Resume preprocessing" button.

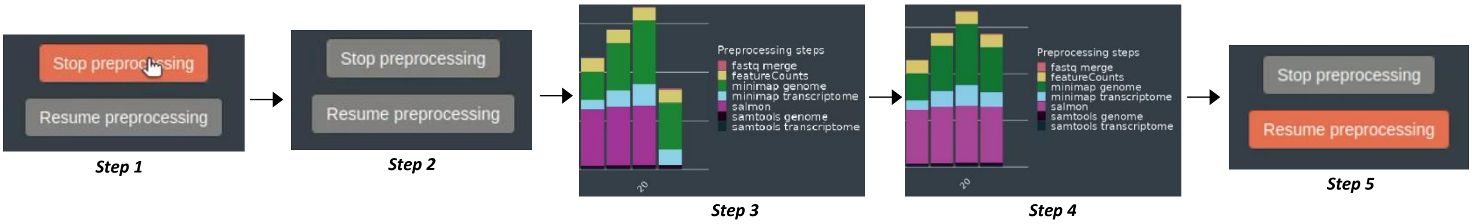


- Once one of the analysis tools is activated, users can check several analysis outputs provided by the tool shown in the “Gene expression”, “Transcript expression” and “Transcript usage” tabs.
- All tools will provide a table with differentially expressed genes/transcript and differentially used transcripts identified between the two conditions. The table can be copied or exported as *csv* or *txt* files.

Gene expression | Transcript expression | Transcript usage

Copy | CSV | Print — Search:

| gene name | transcript name | gene ID | feature ID | exon base mean | p-adjusted | dispersion |
|---|---|---|---|---|---|---|
| PPIA | PPIA-204 | ENSG00000196262.15 | ENST00000468812.6 | 33.0089407864887 | 0.000717803736078183 | 0.00438674551569673 |
| ACTB | ACTB-217 | ENSG00000075624.17 | ENST00000646664.1 | 25.4749219126232 | 0.00163475964243032 | 0.00692619644341414 |
| GAPDH | GAPDH-201 | ENSG00000111640.15 | ENST00000229239.10 | 55.2605867571875 | 0.00164186305365354 | 0.00182380574720786 |
| ENO1 | ENO1-201 | ENSG00000074800.16 | ENST00000234590.10 | 25.8515868294699 | 0.00217993466370087 | 0.00787709904694279 |
| GAPDH | GAPDH-211 | ENSG00000111640.15 | ENST00000619601.1 | 18.0615460838024 | 0.00217993466370087 | 0.00557825559729408 |

- Gene expression and transcript expression analyses will provide tabs to visualize: PCA analysis, volcano plot (DGE), Sample-2-Sample plot and heatmap of the top 20 differentially expressed genes based on p-adjusted.
- All figures can be exported as png figure by clicking the “download” button.

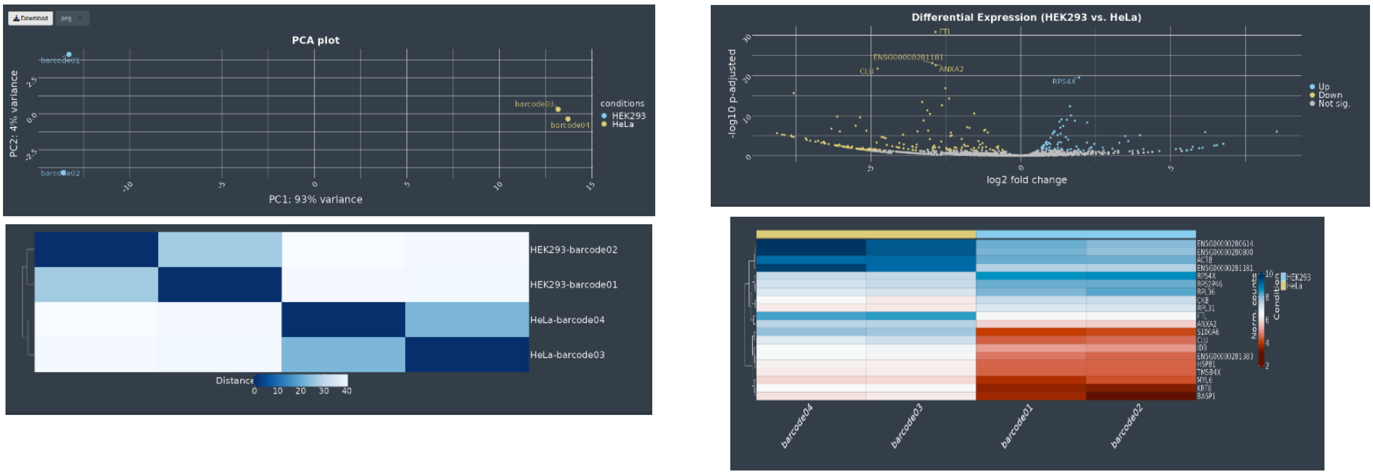

- For differential transcript usage, users can select “general” or “gene specific” tabs.
- In “General” – users can visualize differentially used transcript in a volcano plot, which can be exported as png figure by clicking the “download” button.

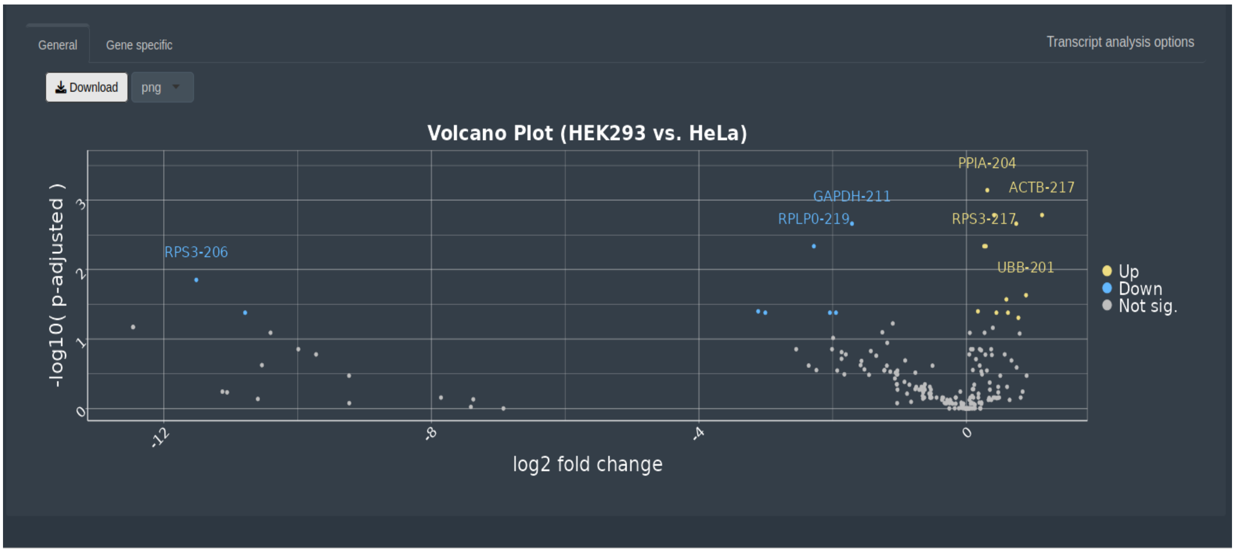


- In “Gene specific” – users can select a specific gene of interest and press “Submit gene selection”.

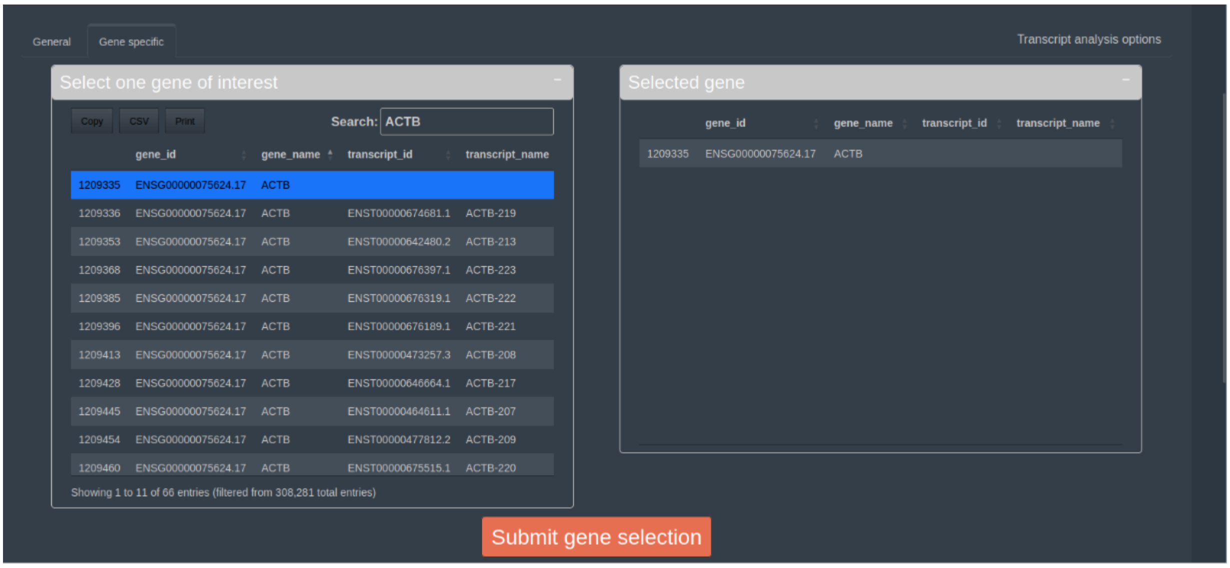

- After submission, a boxplot will appear, displaying the abundance of transcripts within a gene of interest for each condition, based on DRIMSeq's output. The figure can be exported as png figure by clicking the “download” button.

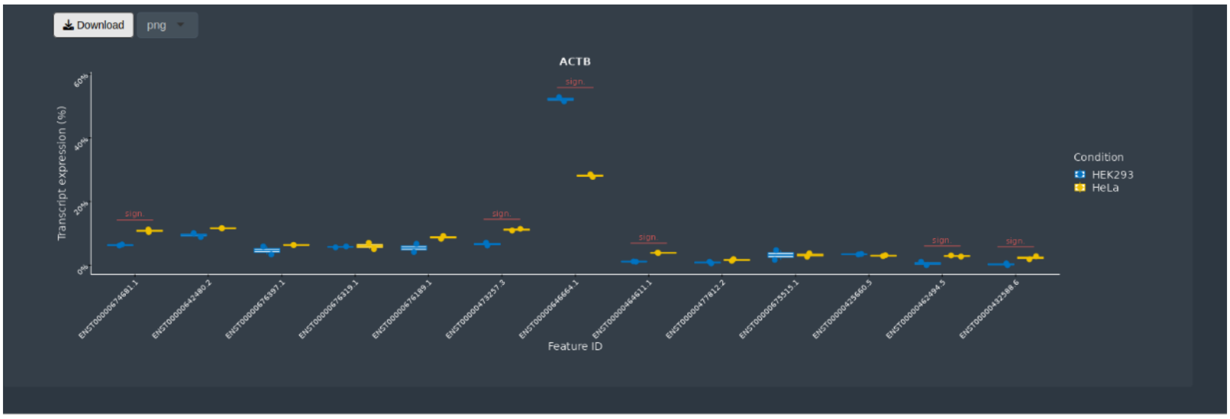


8) ***Gene-wise analysis***

- In the "*Gene-wise analysis*" tab, users can choose between “Gene Counts” and “Gene Body Coverage” tabs.
- In the "Gene Counts" - users can select multiple genes of interest by searching for the gene in the table and clicking on it. The selected gene will then be transferred to the "Selected genes" table, where users can click "Submit Genes" to proceed.

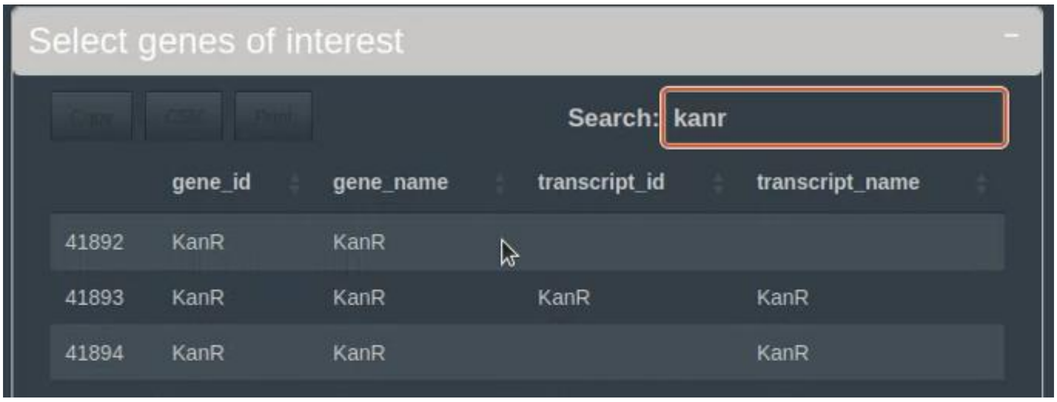


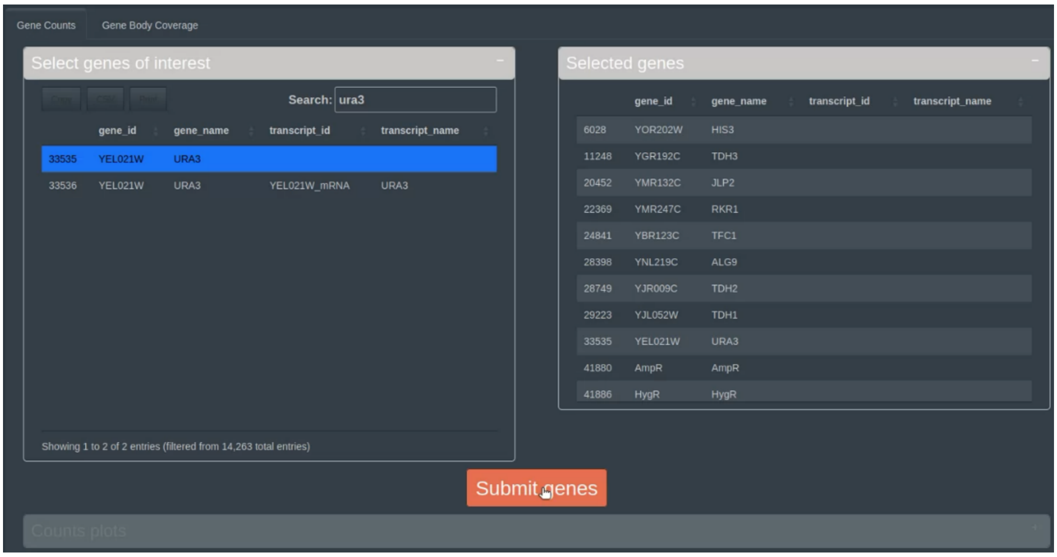

- a figure will appear showing a boxplot/dotplot/violin plot of the raw and normalized read counts per condition. The figures can be exported as png figure by clicking the “download” button.

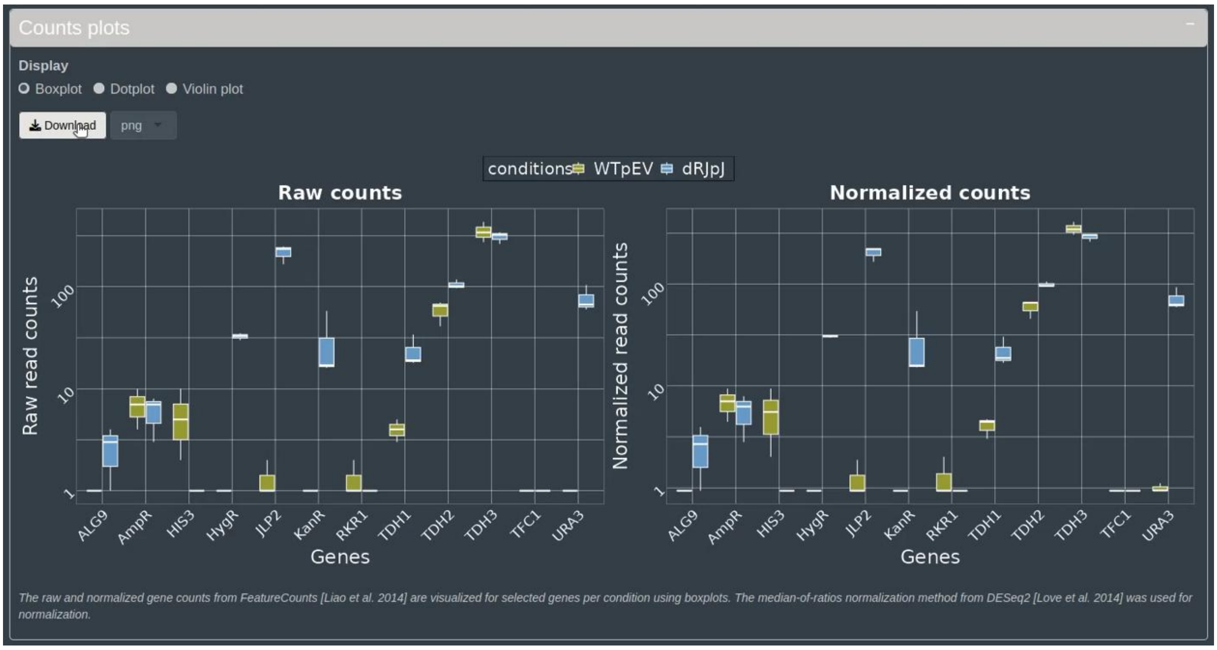


- In the “Gene Body Coverage”, users can select a specific gene of interest and press “Submit gene selection”.
- A figure will display, illustrating the percentage of coverage for exon percentiles for individual samples on the left side, and for the selected conditions on the right side. The figures can be exported as png figure by clicking the “download” button.

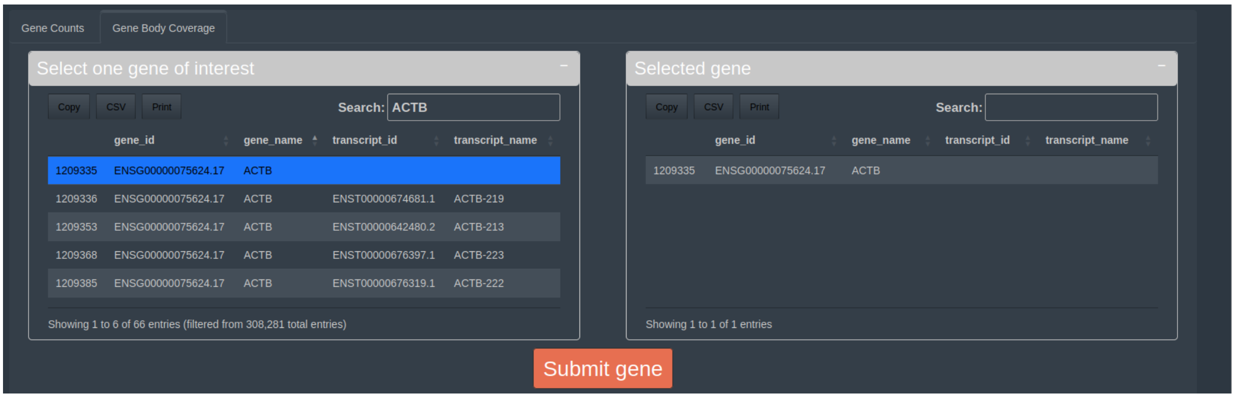


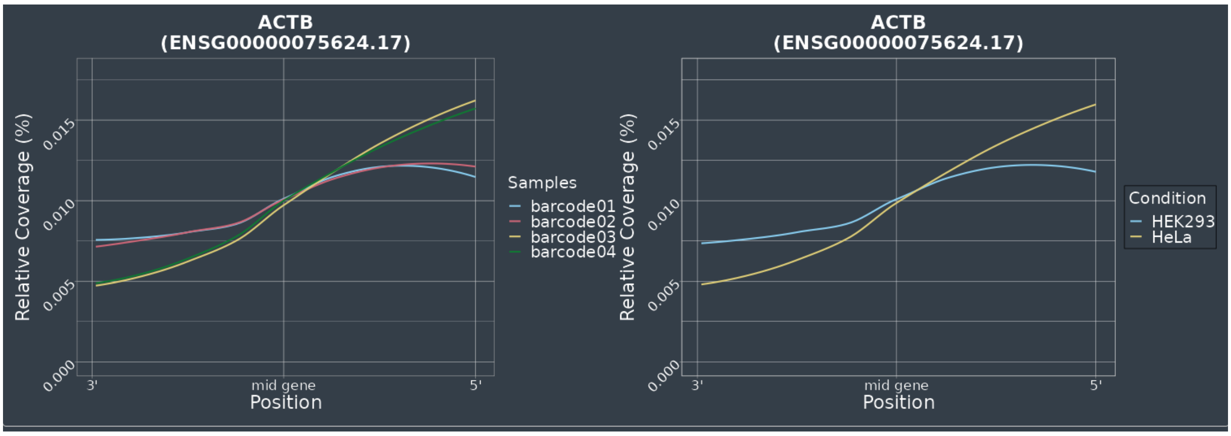

***Supplementary Material 2. Optimizing double-stranded cDNA conversion and loading for Nanopore-Seq.*** During the experimental process, the conversion of double-stranded cDNA (dscDNA) and its subsequent loading into the flow cell were integral steps. Initially, the cDNA library preparation involved using the direct cDNA sequencing kit (SQK-DCS109), but it has since been discontinued. Therefore, we employed an alternative approach utilizing the Maxima H Minus Double-Stranded cDNA Synthesis Kit from Thermo Scientific (K2561), as outlined in the materials and methods. For the HEK and HeLa experiments, NanopoReaTA analysis was assessed on both PromethION (10 replicate and 2 replicate setups) and MinION (2 replicate setup) flow cells (R10 and R9, respectively). The PromethION experimental setup included a 24-hour sequencing period, with data collection intervals at 1hr, 2hr, 5hr, 10hr, and 24hr. In the MinION experimental setup, sequencing extended for 72 hours.

During dscDNA library preparation, two strategies were implemented to evaluate sequencing efficiency for each condition. In the PromethION 10-replicate setup, 300 ng of dscDNA from HEK293 and HeLa cells were used (**Table 1**), while the 2-replicate setup utilized 400 ng of dscDNA (**Table 2**). This variation in cDNA input was based on the total amount of dscDNA required for optimal nanopore sequencing using the PromethION flow cell. For the MinION setup (2 replicates), all available converted dscDNA for HEK293 (~850 ng) and HeLa cells was used (**Table 1**).

Upon utilizing NanopoReaTA's "number of detected gene" utility, we noticed differences in the total number of detected genes between the different setups and flow cells. The 10-replicate PromethION setup showed a relatively balanced number of detected genes (**Figure S1**). However, in the 2-replicate PromethION setup, the number of detected genes was lower in HEK293 compared to HeLa despite loading similar amounts of cDNA (**Figure S2**). In contrast, the MinION setup, which utilized all converted cDNA per replicate, demonstrated a more consistent gene detection rate between conditions (**Figure S3**).

Similarly, the cDNA conversion efficiency differed between RiboM and RiboP samples (**Table 2**). While 400ng of cDNA was taken for the library preparation, RiboM samples did not reach this threshold, hence all cDNA was utilized. Nonetheless, analysis using NanopoReaTA's "number of detected gene" tool revealed that RiboM samples exhibited the highest number of detected genes compared to TotalR and RiboP (**Figure S4**).

These observations raise the question of whether loading a similar amount of cDNA is a suitable strategy for generating even counts between distinct conditions and replicates, or if loading all the converted cDNA into the flow cell, leading to a more even number of reads generated by all replicates, would be preferable. While NanopoReaTA's analysis tools handle read normalization in both cases, addressing these differences is essential for refining best practices in long-read Nanopore RNA-seq experiments.

*Table 1. HEK293 versus HeLa experimental setup (10 replicates per condition). Total Amount of cDNA Converted from 2µg RNA and utilized in PromethION and MinION Sequencing.*

| | | HEK293 | | | | | | | | | | HeLa | | | | | | | | | | cDNA taken for sequencing |
|---|---|---|---|---|---|---|---|---|---|---|---|---|---|---|---|---|---|---|---|---|---|---|
| | | R1 | R2 | R3 | R4 | R5 | R6 | R7 | R8 | R9 | R10 | R1 | R2 | R3 | R4 | R5 | R6 | R7 | R8 | R9 | R10 | |
| *Total amount of cDNA generated (ng)* | **PromethION** | 516 | 496 | 480 | 516 | 508 | 512 | 464 | 480 | 456 | 532 | 348 | 404 | 424 | 420 | 508 | 492 | 440 | 468 | 488 | 432 | 300ng (both HEK293 and HeLa) |

*Table 2. HEK293 versus HeLa experimental setup (2 replicates per condition). Total Amount of cDNA Converted from 2µg RNA and utilized in PromethION and MinION Sequencing.*

| | | HEK293 | | HeLa | | cDNA taken for sequencing |
|---|---|---|---|---|---|---|
| | | R1 | R2 | R1 | R2 | |
| *Total amount of cDNA generated (ng)* | **PromethION** | 1,116 (bc1) | 1,172 (bc2) | 492 (bc3) | 440 (bc4 | 400ng (both HEK293 and HeLa) |
| | **MinION** | 888 (bc4) | 824 (bc5) | 452 (bc6) | 564 (bc7) | All cDNA per replicate |

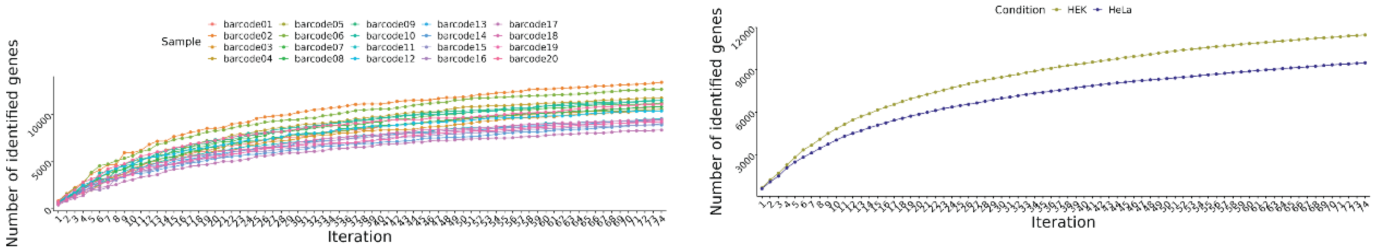


**Figure S1. PromethION 10 replicate experimental setup**

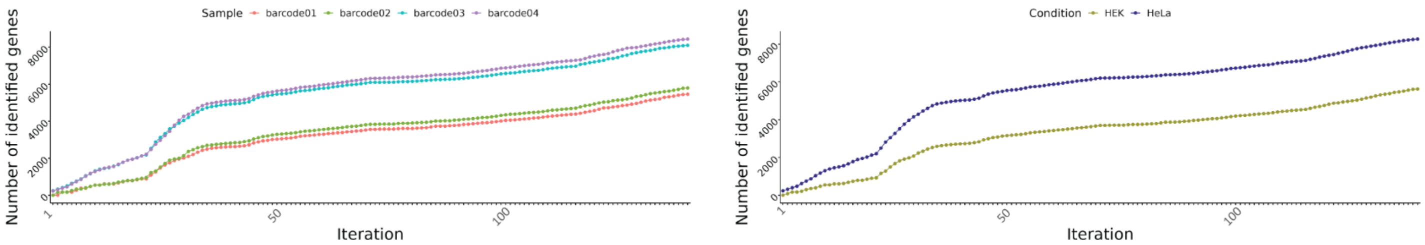


**Figure S2. PromethION 2 replicate experimental setup**

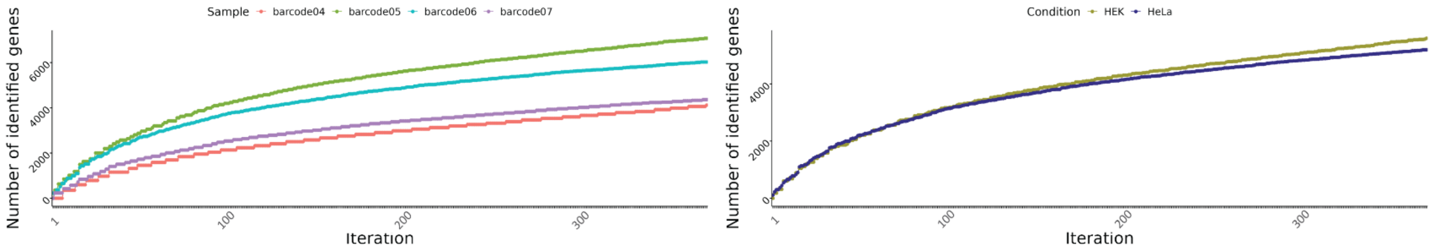


**Figure S3. MinION 2 replicate experimental setup**

*Table 3. rRNA-depleted and rRNA-enriched transcripts setup. Total amount of cDNA converted from 2µg RNA (500ng for RiboM) and utilized in PromethION sequencing.*

| | **Total R** | | **Ribominus** | | **Riboplus** | |
|---|---|---|---|---|---|---|
| | **R1** | **R2** | **R1** | **R2** | R1 | R2 |
| *Total amount of cDNA generated (ng)* | 1,116 (bc1) | 1,172 (bc2) | 132 (bc5) | 163 (bc6) | 364 (bc7) | 2,120 (bc8) |
| *cDNA taken for sequencing* | 400ng | 400ng | 132ng | 163ng | 364ng | 400ng |

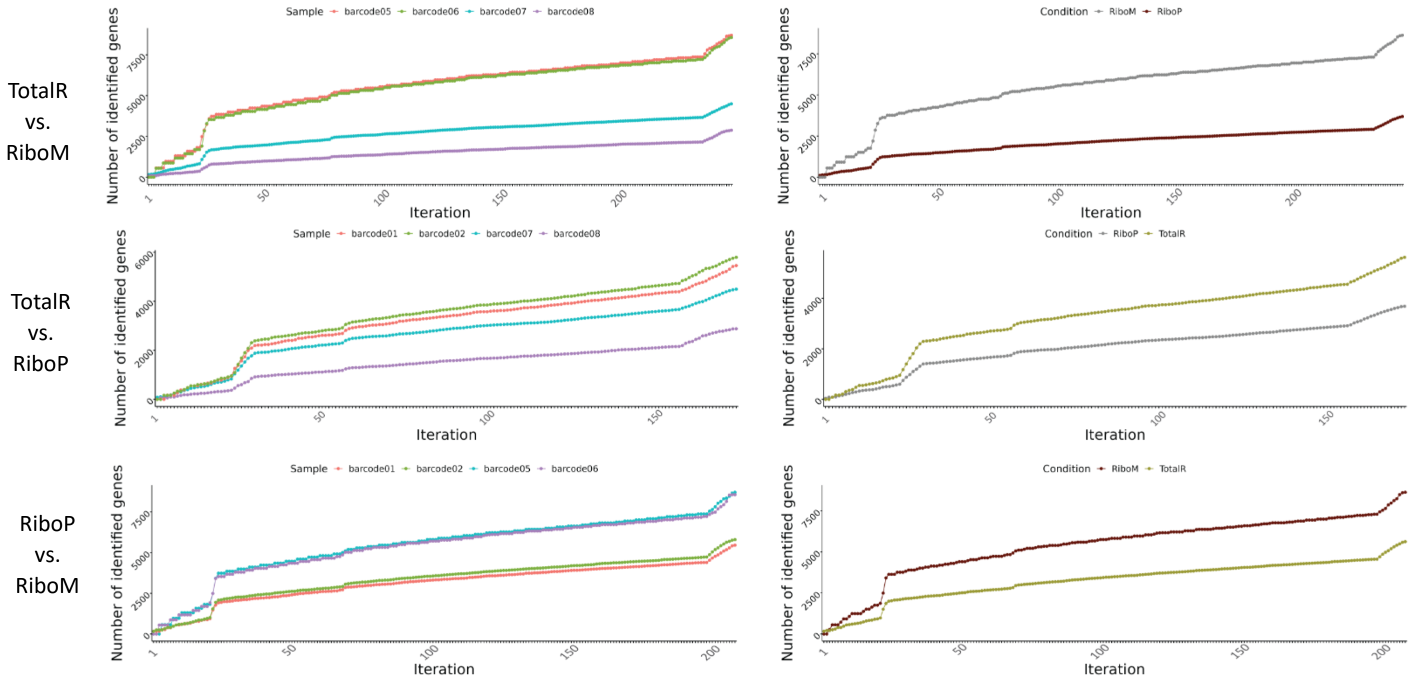


**Figure S4. Comparison of # genes detected in TotalR/Ribominus/Riboplus**

***Supplementary Material 3. Pooling of cDNA libraries after barcoding.***
In the yeast experimental setups, we observed low, yet noticeable abundance of deleted genes by examining the normalized read count. This concerned notably *NEW1* in Yeast setup 1, and *JLP2* in Yeast setup 2. Additionally, we detected the presence of antibiotic resistance genes in WT conditions where the expression of *KanR* or *HygR* should not have been present. Evidence of this discrepancy can be seen in the IGV snapshot, where a few reads were captured in conditions where they should not be expressed (Supplementary Figure 56, see figure below). As one example, in Yeast setup 1, *HygR* was not detected in any sample, since none of the strains expressed HygR within this sequencing setup. However, in yeast setup 2, the identical WT-pEV(*HIS3*), (an aliquot of the identical RNA sample used in setup 1) was included in the library preparation and sequenced alongside the *rkr1Δ::HphMX* strains expressing *HygR*. As a result, we observed a few reads for *HygR* that mapped to the WT-pEV(*HIS3*) samples. This strongly suggests that detection of sequences that should not be present within strains is due to barcode cross-contamination during library preparation (Explanation described below). The same phenomenon can be seen for all other genes mentioned above. Only for the high detection of *HIS3* in all samples of setup 2 there is a different reason: The WT strain BY4741 bears the *his3Δ1* allele, which contains an internal deletion of 187-bp within the *HIS3* gene (Daniel et al. 2006). Beyond this internal deletion, which is clearly visible in Supplementary Figure 56 (see figure below), the remainder of the gene is still expressed at high levels and therefore detected and aligned to the *HIS3* gene, which is expected in this case.

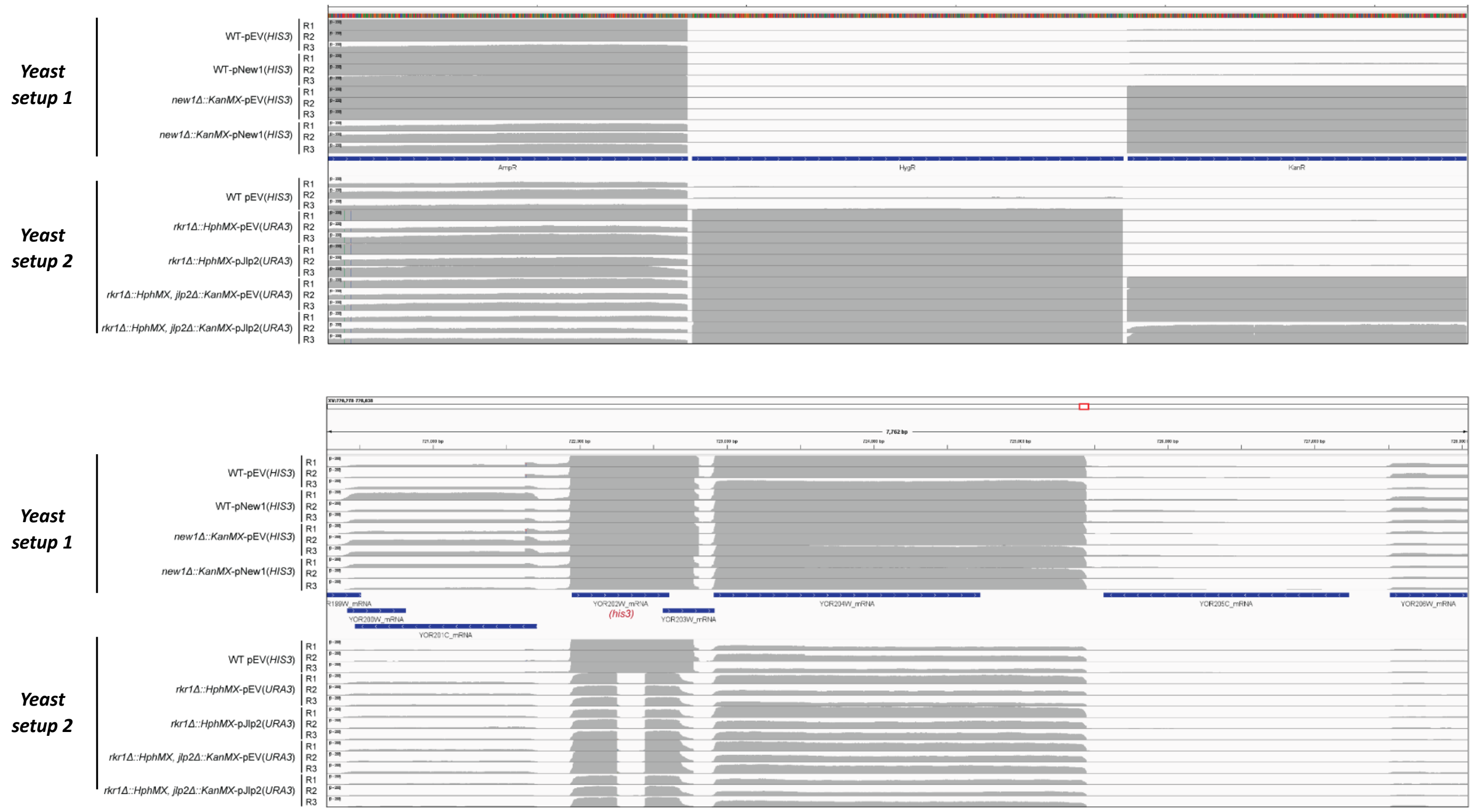


We believe that one possible explanation of these discrepancies is potential barcode mixup that occurs during library preparation in the sample barcoding step. During the barcoding stage, a Blunt/TA Ligase is used to ligate the barcodes to the dscDNA (for 20min). The following step includes incubation of the sample with EDTA (supplied by the kit, EXP-NBD104, ONT) to inhibit the reaction and pooling all the barcodes together. We think that during this step of pooling, despite samples being supplemented with EDTA, it could be that some samples are still barcoded by other barcodes from the pool during mixing and therefore, we see a small, yet visible detection of genes that should not be present in the group. EDTA enzymatic inhibition occurs in a time-dependent manner (Gonzálvo et al., 1997), whereas in the protocol these steps occur relatively quickly with no longer incubation of samples with EDTA. Thus, we suggest two strategies to overcome this challenge. 1. The samples could be incubated with EDTA for at least 20min to ensure proper inhibition of the enzymatic reaction so they will not cross to other samples. 2. Alternatively, each sample could be individually isolated after the barcoding step using Ampure XP bead purification. However, this approach carries the risk of material loss due to the small amount of cDNA being handled. Nonetheless, leveraging the rapid detection capabilities of NanopoReaTA, we quickly identified these discrepancies, showcasing its potential as an effective quality control tool for various experimental setups.

**References**

Wierczeiko A, Pastore S, Mündnich S, Busch AM, Dietrich V, Helm M, Butto T, Gerber S. 2023. NanopoReaTA: a user-friendly tool for nanopore-seq real-time transcriptional analysis. Bioinformatics, 39(8).

Daniel JA, Yoo J, Bettinger BT, Amberg DC, Burke DJ. 2006. Eliminating gene conversion improves high-throughput genetics in Saccharomyces cerevisiae. Genetics. 172(1):709-11.

Gonzálvo MC, Gil F, Hernández AF, Villanueva E, Pla A. 1997. Inhibition of paraoxonase activity in human liver microsomes by exposure to EDTA, metals and mercurials. Chemico-Biological Interactions, 105(3), 169–179.
