## Supplementary Document S1 for "Real-time transcriptomic profiling in distinct experimental conditions"

### Supplementary Figures 1-10, related to Figure 2

#### HEK293 vs HeLa Comparison

**Supplementary Figure 1 |** Sequencing overview HEK293 vs HeLa. .......... 2
**Supplementary Figure 2 |** PromethION (10 Rep) - General overview HEK293 vs HeLa. .......... 4
**Supplementary Figure 3 |** PromethION (2 Rep) - General overview HEK293 vs HeLa. .......... 6
**Supplementary Figure 4 |** MinION (2 Rep) - General overview HEK293 vs HeLa. .......... 8
**Supplementary Figure 5 |** PromethION (10 Rep) - Gene expression HEK293 vs HeLa. .......... 9
**Supplementary Figure 6 |** PromethION (2 Rep) - Gene expression HEK293 vs HeLa. .......... 10
**Supplementary Figure 7 |** MinION (2 Rep) - Gene and Transcript expression HEK293 vs HeLa. .......... 11
**Supplementary Figure 8 |** PromethION (10 Rep) - Transcript expression HEK293 vs HeLa .......... 12
**Supplementary Figure 9 |** PromethION (2 Rep) - Transcript expression HEK293 vs HeLa .......... 13
**Supplementary Figure 10 |** PromethION (2 Rep) - Transcript expression HEK293 vs HeLa .......... 14

#### *General supplementary figures description*

**Sequencing overview** – Refers to sequencing metrics including total reads generated per barcode/sample, mapped reads, gene counts and transcript counts.
**General overview –** Refers to data collected from NanopoReaTA regaring the general sample/condition overview including: Read length overview (per sample and condition), gene expression variability (per sample and condition), change in gene composition (per sample and condition) and Process time.
**Gene expression** - Refers to data collected from NanopoReaTA regaring differential gene expression analysis including: Principal Component Analysis (PCA), Volcano plots, Sample-to-sample variaiblity and Heatmaps.
**Transcript expression** - Refers to data collected from NanopoReaTA regaring differential transcript expression analysis including: Principal Component Analysis (PCA), Volcano plot, Sample-to-sample variaiblity and Heatmaps.

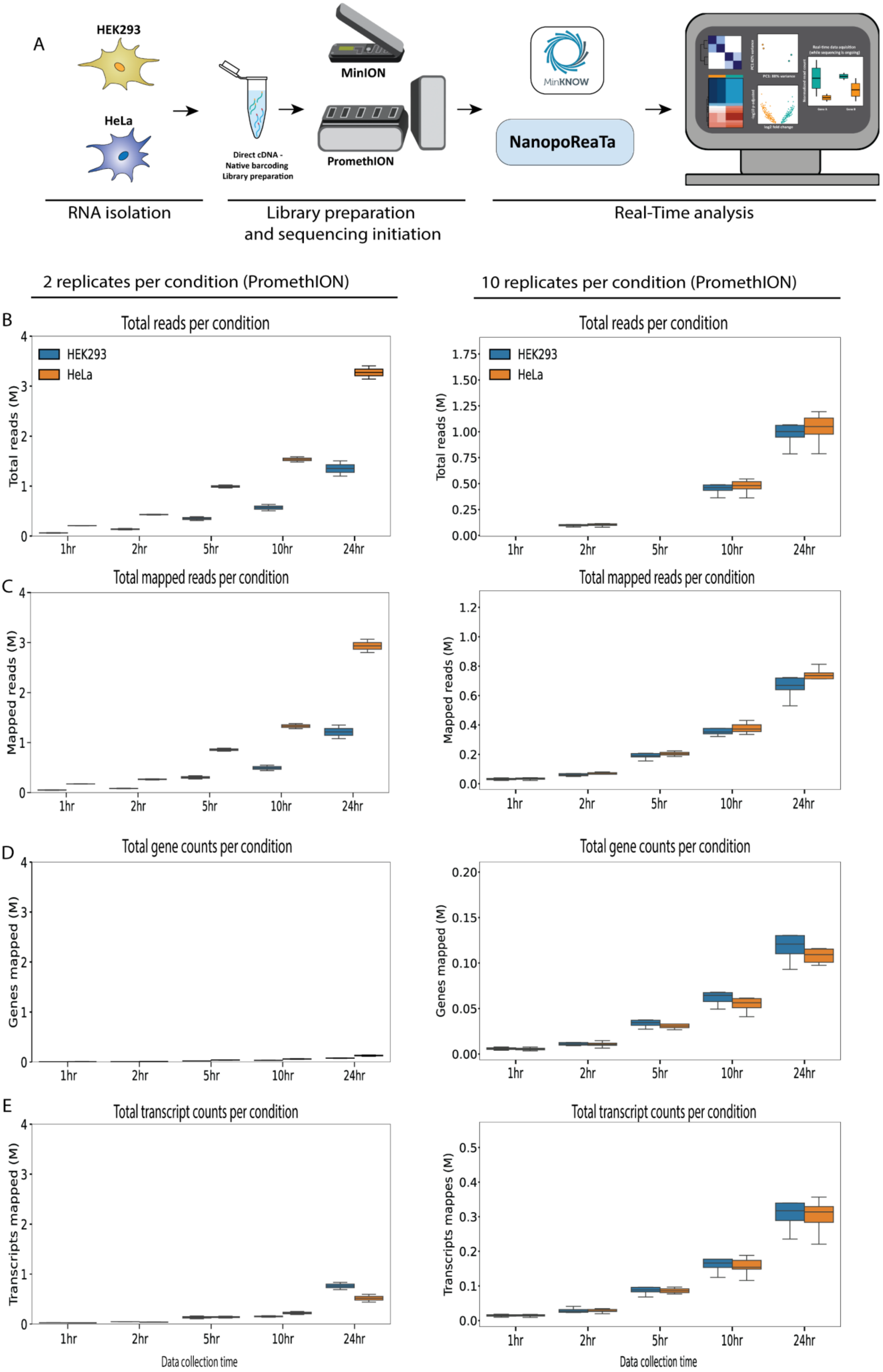
A
HEK293
HeLa
Direct cDNA - Native barcoding Library preparation
MinION
PromethION
MinKNOW
NanopoReaTa
RNA isolation
Library preparation and sequencing initiation
Real-Time analysis
2 replicates per condition (PromethION)
10 replicates per condition (PromethION)
B
Total reads per condition
Total reads (M)
C
Total mapped reads per condition
Mapped reads (M)
D
Total gene counts per condition
Genes mapped (M)
E
Total transcript counts per condition
Transcripts mapped (M)
Transcripts mappes (M)
1hr
2hr
5hr
10hr
24hr
Data collection time

**Supplementary Figure 1. Experimental strategy of HEK293 versus HeLa and sequencing overview. A.** RNA was isolated from HEK293 and HeLa cells dscDNA library was prepared which included samples barcoding and adapter ligation. Samples were loaded and sequenced using a PromethION R10 flow cell or MinION R9 flow cell. NanopoReaTA was activated shortly after sequencing initiation and data was collected 1hr, 2hr, 5hr, 10hr, and 24hr post-sequencing initiation. **B-E.** The box plot represent total read generated (**B**), total mapped reads (**C**), total gene counts (**D**) and total transcript count (**E**) per condition. The data for 2 replicates and 10 replicates per condition are shown in left and right, respectively.

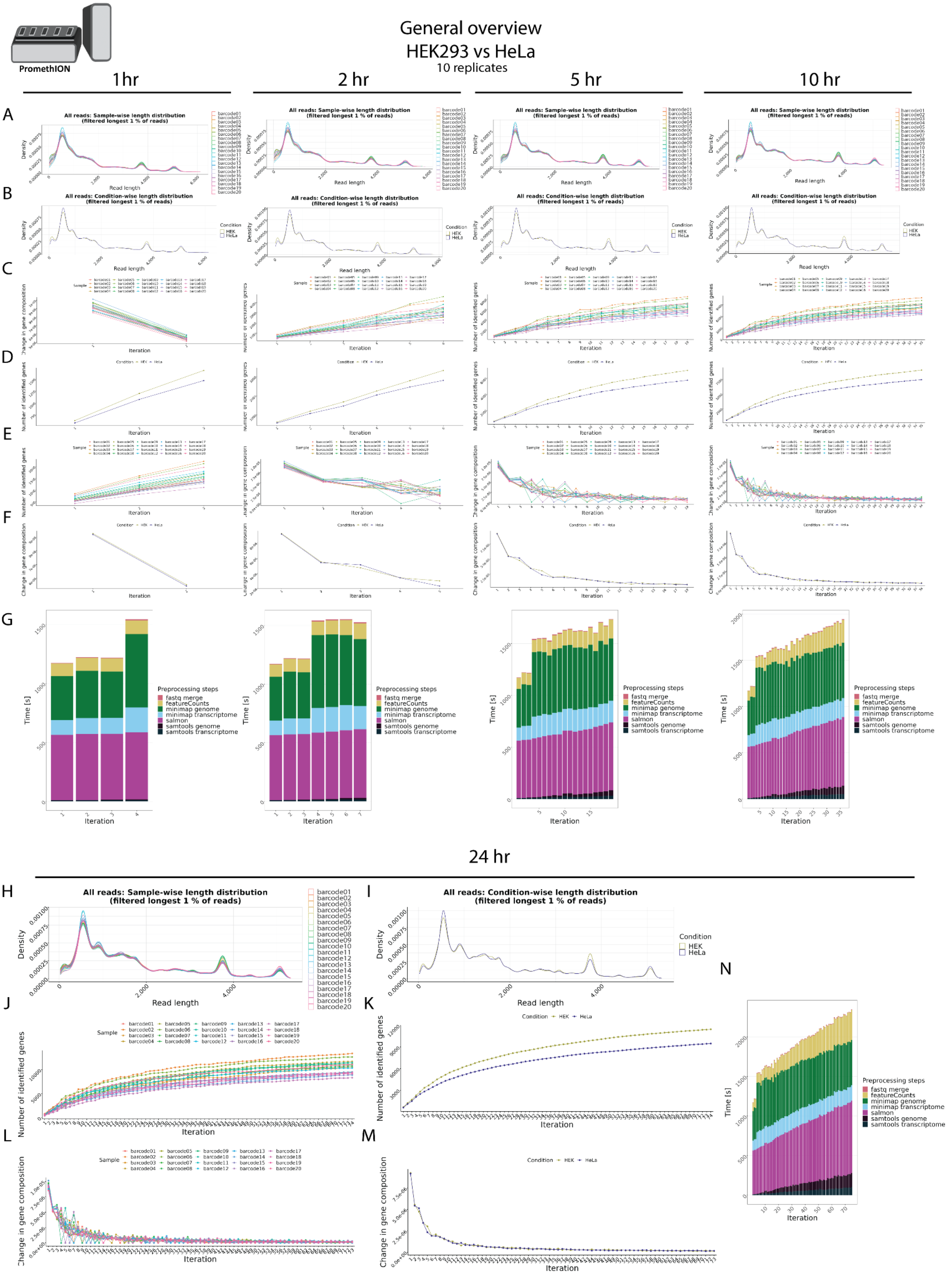
PromethION
General overview
HEK293 vs HeLa
10 replicates
1 hr
2 hr
5 hr
10 hr
24 hr
All reads: Sample-wise length distribution
(filtered longest 1 % of reads)
All reads: Condition-wise length distribution
(filtered longest 1 % of reads)
Density
Read length
Condition
HEK
HeLa
Sample
Number of identified genes
Change in gene composition
Iteration
Time [s]
Preprocessing steps
fastq merge
featureCounts
minimap genome
minimap transcriptome
salmon
samtools genome
samtools transcriptome

**Supplementary Figure 2. PromethION (10 rep) - General overview in HEK293 vs HeLa**. **A-B**. Read length overview. The distribution of read lengths derived from generated fastq files is plotted per sample (A) and per condition (B). All reads of length over the 99 % quantile of all lengths are removed from these visualizations. **C-D**. Gene expression variability. The number of identified genes (> 0 reads counted) is plotted after each iteration per sample (C) and per condition (D). When the detection of additional genes finishes, the lines reach a plateau. **E-F**. Change in gene composition. The change in gene composition (CGC) plot shows the sum of absolute differences of relative gene (%gi) counts per individual gene (g) in iteration (i) to the relative gene counts in iteration (i-1) for each respective sample (E) and condition (F). CGC = sum(|(%g(i-1))-(%g(i))|). **G.** Process time. The bar plot shows the time in seconds all preprocessing steps needed per iteration. One iteration process at maximum 30 files from all samples. This plot updates automatically as a new process finish. **H-N**. Final general overview. Read length (H-I), Gene expression variability (J-K), Change in gene composition (L-M), and processing time (N) for the 24hr time point. All plots are organized according to their respective time points of collection.

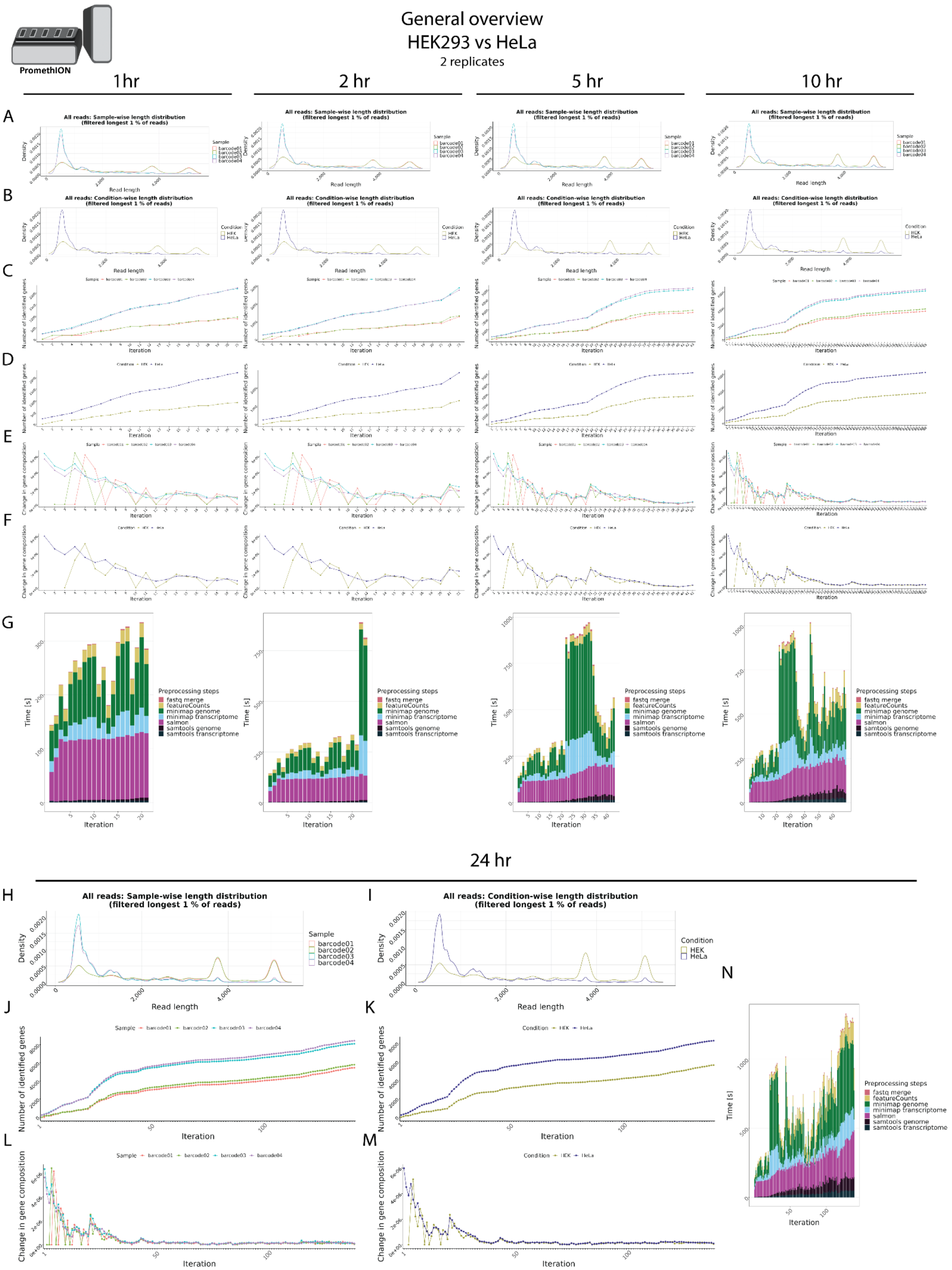

PromethION
General overview
HEK293 vs HeLa
2 replicates
1 hr
2 hr
5 hr
10 hr
A
B
C
D
E
F
G
All reads: Sample-wise length distribution
(filtered longest 1 % of reads)
All reads: Condition-wise length distribution
(filtered longest 1 % of reads)
Density
Read length
Sample
barcode01
barcode02
barcode03
barcode04
Condition
HEK
HeLa
Number of identified genes
Change in gene composition
Iteration
Time [s]
Preprocessing steps
fastq merge
featureCounts
minimap genome
minimap transcriptome
salmon
samtools genome
samtools transcriptome
24 hr
H
I
J
K
L
M
N

**Supplementary Figure 3. PromethION (2 rep)- General overview in HEK293 vs HeLa**. **A-B**. Read length overview. The distribution of read lengths derived from generated fastq files is plotted per sample (A) and per condition (B). All reads of length over the 99 % quantile of all lengths are removed from these visualizations. **C-D**. Gene expression variability. The number of identified genes (> 0 reads counted) is plotted after each iteration per sample (C) and per condition (D). When the detection of additional genes finishes, the lines reach a plateau. **E-F**. Change in gene composition. The change in gene composition (CGC) plot shows the sum of absolute differences of relative gene (%gi) counts per individual gene (g) in iteration (i) to the relative gene counts in iteration (i-1) for each respective sample (E) and condition (F). CGC = sum(|(%g(i-1))-(%g(i))|). **G.** Process time. The bar plots show the time in seconds all preprocessing steps needed per iteration. One iteration process at maximum 30 files from all samples. This plot updates automatically as a new process finish. **H-N**. Final general overview. Read length (H-I), Gene expression variability (J-K), Change in gene composition (L-M), and processing time (N) for the 24hr time point. All plots are organized according to their respective time points of collection.

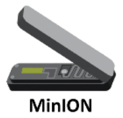


General overview
HEK293 vs HeLa
2 replicates

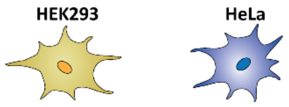


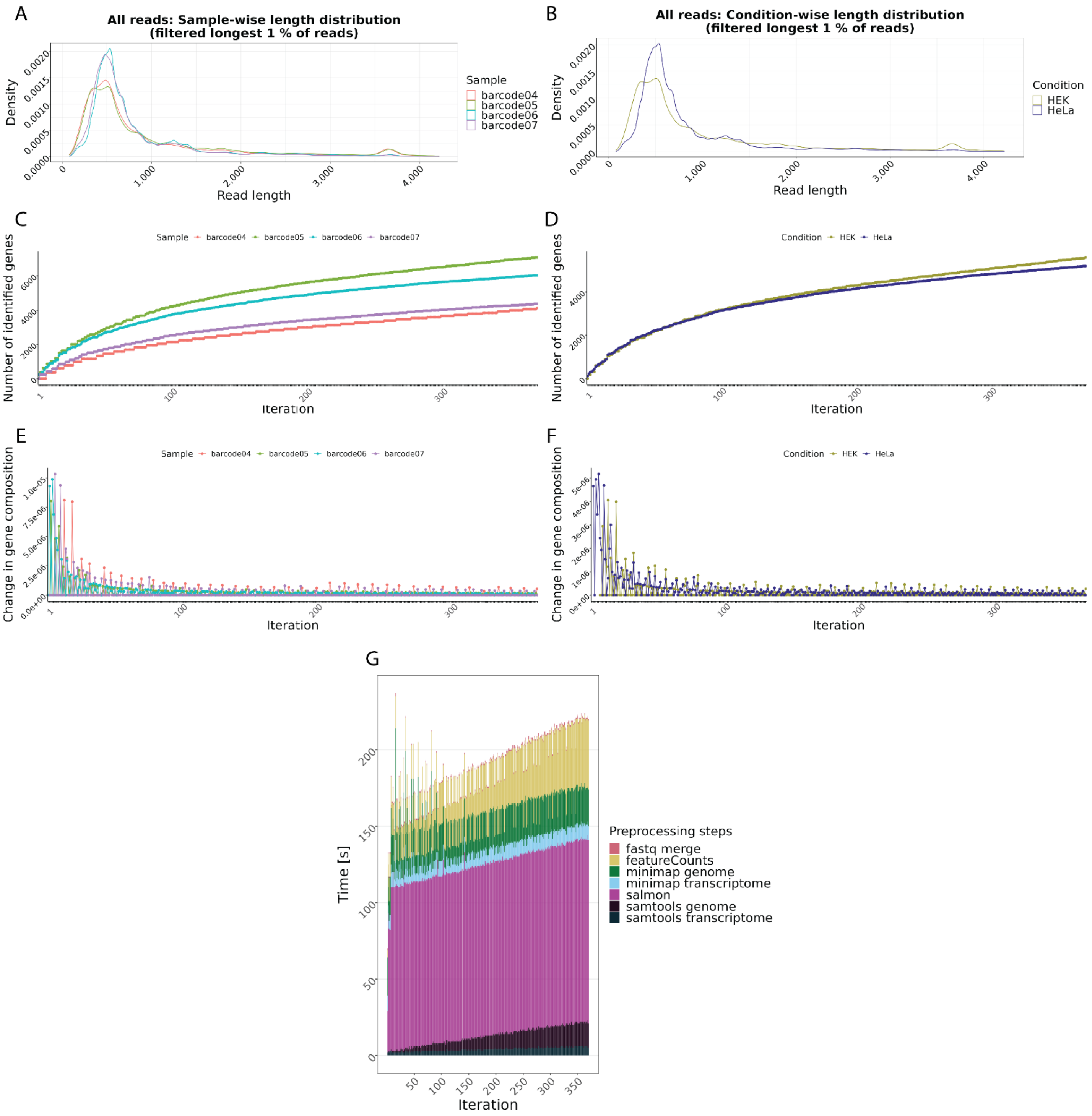


**Supplementary Figure 4. MinION - General overview HEK293 vs HeLa. A-B**. Read length overview. The distribution of read lengths derived from generated fastq files is plotted per sample (A) and per condition (B). All reads of length over the 99 % quantile of all lengths are removed from these visualizations. **C-D**. Gene expression variability. The number of identified genes (> 0 reads counted) is plotted after each iteration per sample (C) and per condition (D). When the detection of additional genes finishes, the lines reach a plateau. **E-F**. Change in gene composition. The change in gene composition (CGC) plot shows the sum of absolute differences of relative gene (%gi) counts per individual gene (g) in iteration (i) to the relative gene counts in iteration (i-1) for each respective sample (E) and condition (F). CGC = sum(|(%g(i-1))-(%g(i))|). **G.** Process time. The bar plots show the time in seconds all preprocessing steps needed per iteration. One iteration process at maximum 30 files from all samples. This plot updates automatically as a new process finish.

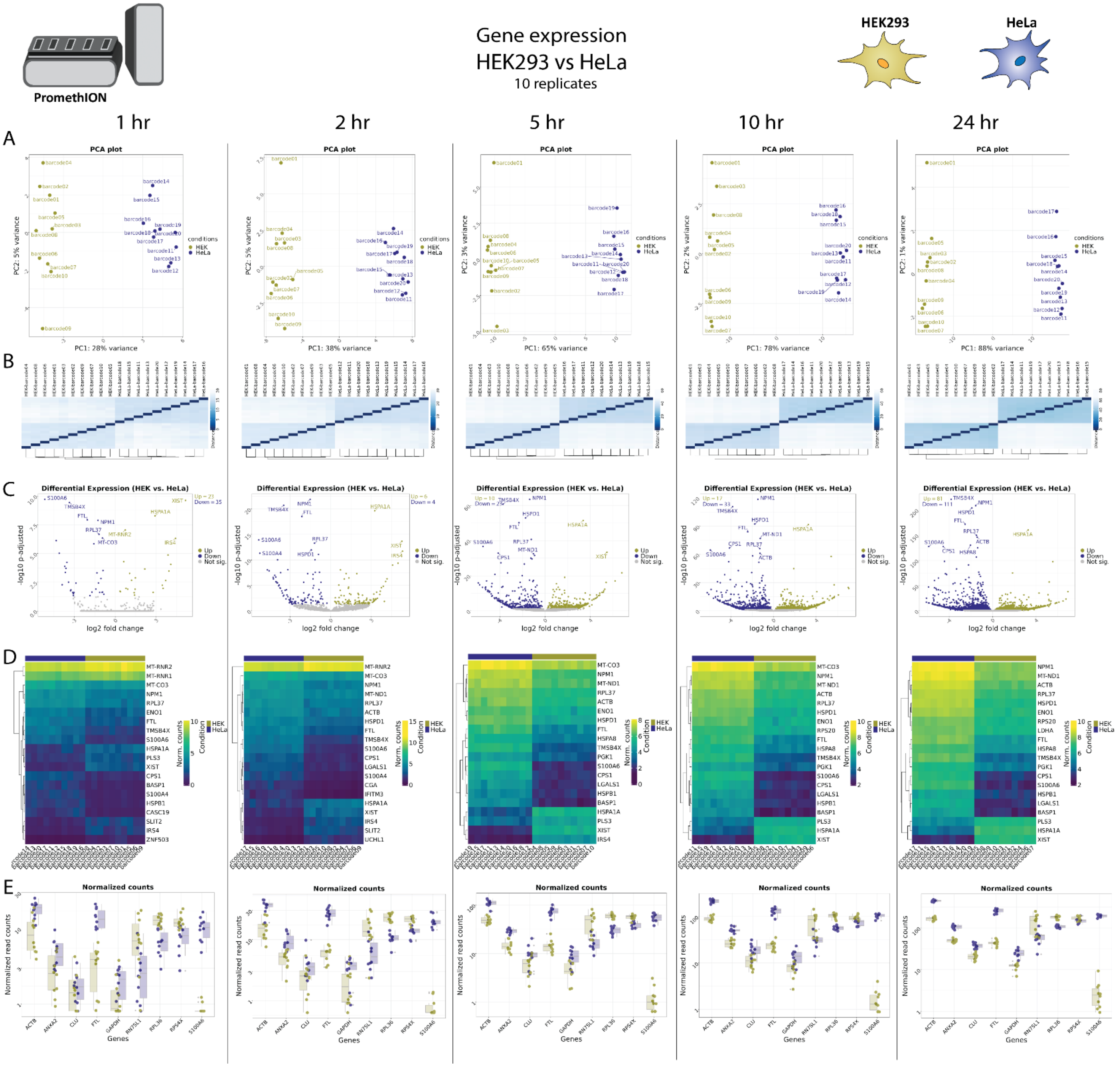


**Supplementary Figure 5. PromethION (10 rep) - Gene expression analysis in HEK293 vs HeLa. A.** Principal Component Analysis (PCA) of the top 500 genes with the highest variance across all samples, the two Principal Components. Each dot corresponds to one sample and is colored by the respective condition. **B.** Sample2sample distance plot.The euclidean distance between the gene expression patterns of all samples to each other is plotted using heatmap. Normalized gene counts were used for distance computation. **C.** Volcano plots. The differential gene expression analysis results from DESeq2 are shown by plotting the log2FoldChange between the conditions of interest against the –log10 adjusted p-value per gene observed from the Wald-Test integrated in DESeq2 [Love, Huber, and Anders et al. 2014]. The top 10 significant genes are labeled by their symbol. **D.** Heatmaps. The expression of the top 20 differentially expressed genes is plotted using Heatmap by coloring the number of reads per gene. The legend on the right side describes which colors correspond to highly and lowly expressed genes. **E.** Normalized gene count for selected genes. Normalized gene counts from FeatureCounts [Liao et al. 2014] are visualized for selected genes per condition using boxplots. The median-of-ratios normalization method from DESeq2 [Love et al. 2014] was used for normalization.

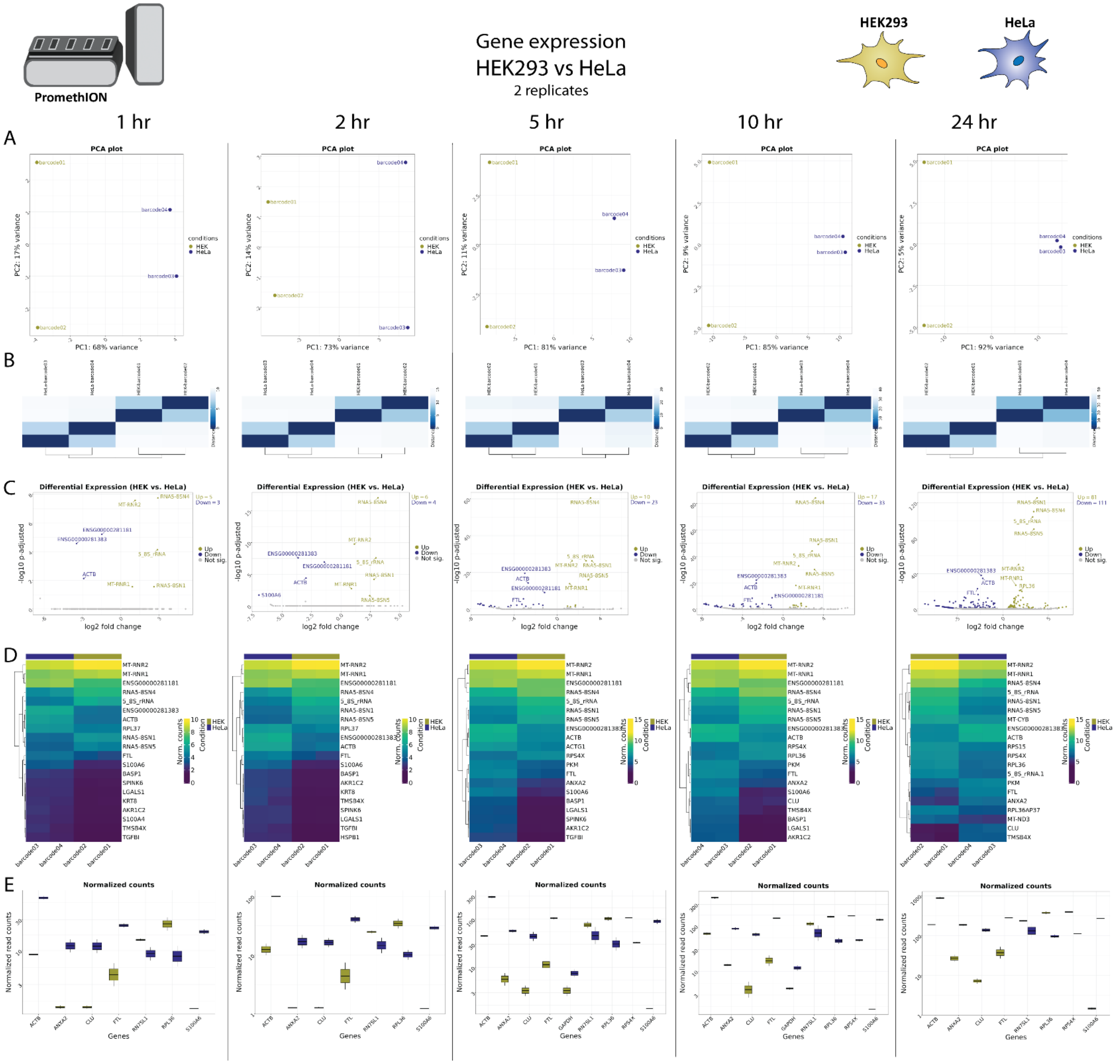


**Supplementary Figure 6. PromethION - Gene expression analysis in HEK293 vs HeLa. A.** Principal Component Analysis (PCA) of the top 500 genes with the highest variance across all samples, the two Principal Components. Each dot corresponds to one sample and is colored by the respective condition. **B.** Sample2sample distance plot.The euclidean distance between the gene expression patterns of all samples to each other is plotted using heatmap. Normalized gene counts were used for distance computation. **C.** Volcano plots. The differential gene expression analysis results from DESeq2 are shown by plotting the log2FoldChange between the conditions of interest against the –log10 adjusted p-value per gene observed from the Wald-Test integrated in DESeq2 [Love, Huber, and Anders et al. 2014]. The top 10 significant genes are labeled by their symbol. **D.** Heatmaps. The expression of the top 20 differentially expressed genes is plotted using Heatmap by coloring the number of reads per gene. The legend on the right side describes which colors correspond to highly and lowly expressed genes. **E**. Normalized gene count for selected genes. Normalized gene counts from FeatureCounts [Liao et al. 2014] are visualized for selected genes per condition using boxplots. The median-of-ratios normalization method from DESeq2 [Love et al. 2014] was used for normalization.

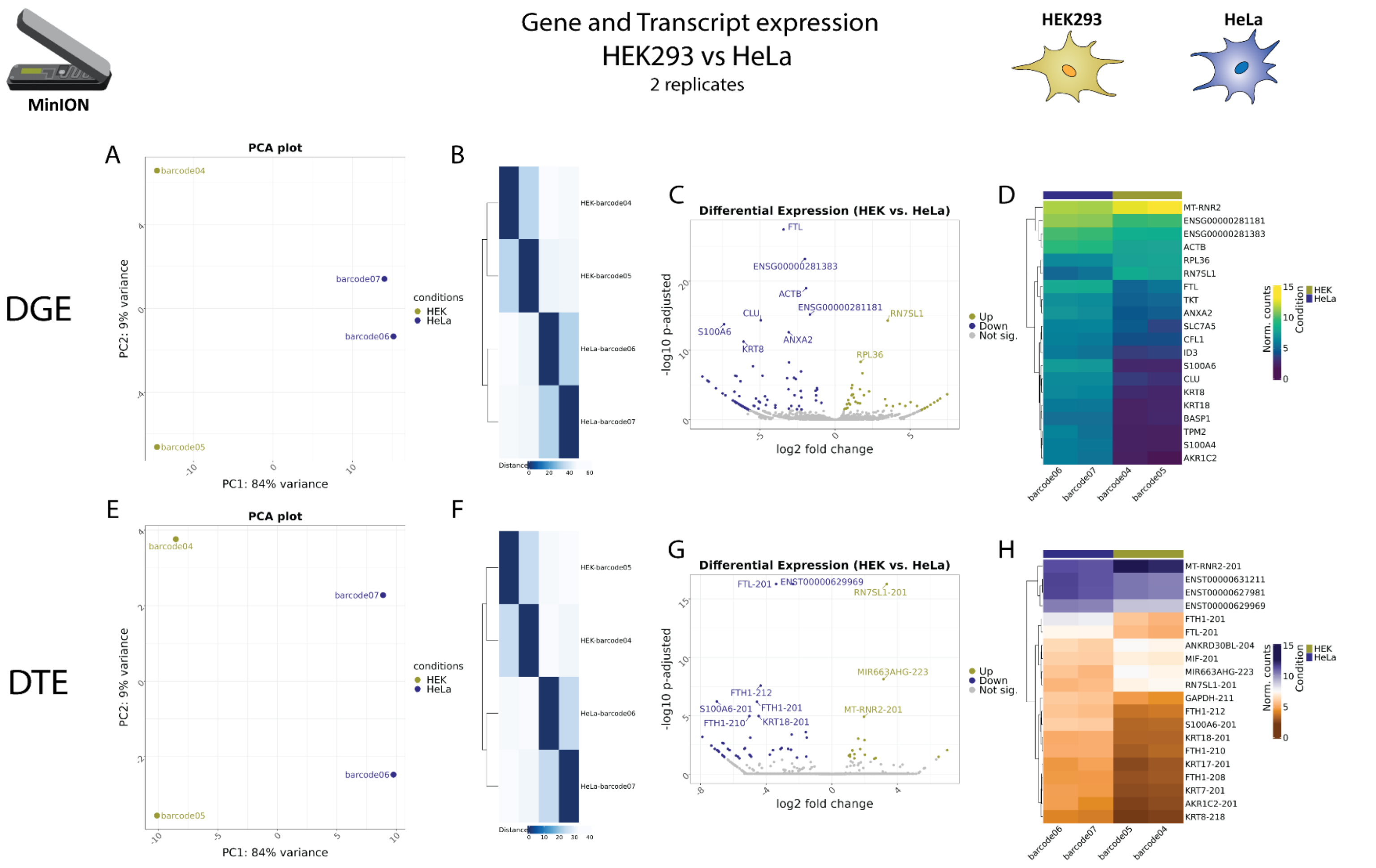


**Supplementary Figure 7. MinION – Gene and Transcript expression in HEK293 vs HeLa. A-D** DGE analysis. **A.** Principal Component Analysis (PCA) of the top 500 genes with the highest variance across all samples. **B.** The euclidean distance between the gene expression patterns of all samples to each other is plotted using heatmap. Normalized gene counts were used for distance computation. **C.** The differential gene expression analysis results from DESeq2 are shown by plotting the log2FoldChange between the conditions of interest against the –log10 adjusted p-value per gene observed from the Wald-Test integrated in DESeq2 **D.** The expression of the top 20 differentially expressed genes is plotted using Heatmap by coloring the number of reads per gene. **E-H** DTE analysis. **E.** Principal Component Analysis (PCA) of the top 500 transcripts with the highest variance across all samples. **F.** The euclidean distance between the transcript expression patterns of all samples to each other is plotted using heatmap. Normalized transcript counts were used for distance computation. **G.** The differential transcript expression analysis are shown by plotting the log2FoldChange between the conditions of interest against the –log10 adjusted p-value per transcript observed from the Wald-Test integrated in DESeq2 **H.** The expression of the top 20 differentially expressed transcripts is plotted using Heatmap by coloring the number of reads per gene.

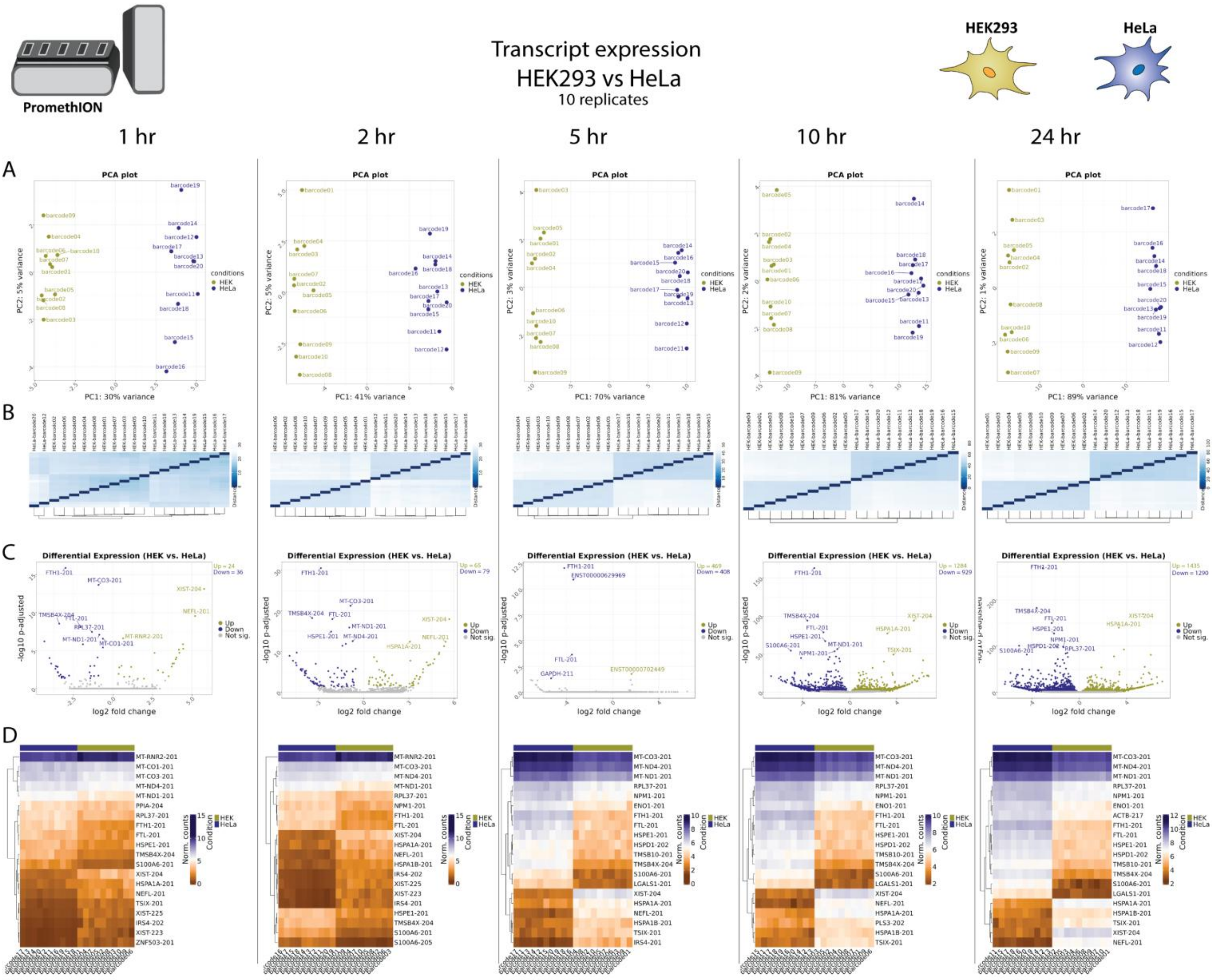


**Supplementary Figure 8. Transcript expression in HEK293 vs HeLa (10 rep). A.** Principal Component Analysis (PCA) of the top 500 transcripts with the highest variance across all samples, the two Principal Components, PC1 and PC2 - that explain the most variance in the dataset are plotted against each other. Each dot corresponds to one sample and is colored by the respective condition. **B.** Sample2sample plot. The euclidean distance between the transcript expression patterns of all samples to each other is plotted using heatmap. Normalized gene counts were used for distance computation. **C.** Volcano plots. The differential transcript expression analysis results from DESeq2 are shown by plotting the log2FoldChange between the conditions of interest against the –log10 adjusted p-value per gene observed from the Wald-Test integrated in DESeq2. The top 10 significant genes are labeled by their symbol. **D.** Heatmaps. The expression of the top 20 differentially expressed transcripts is plotted using Heatmap by coloring the number of reads per transcript. The legend on the right side describes which colors correspond to highly and lowly expressed transcripts. **E**. Differential transcript usage volcano plot. The log2FoldChange values of the transcripts from DEXSeq (Anders et al. 2012) are plotted against the –log10 adjusted p-value.

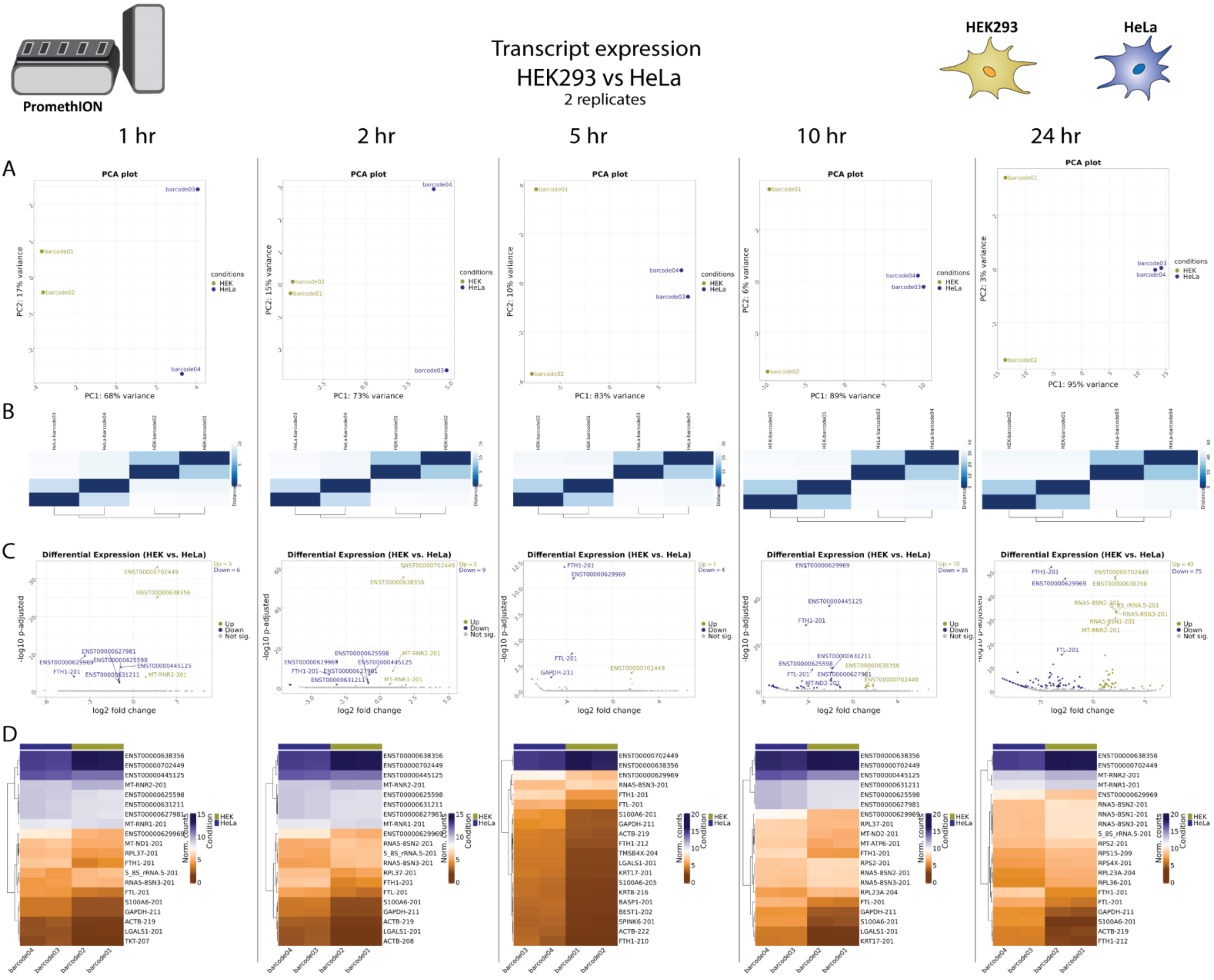


**Supplementary Figure 9. Transcript expression in HEK293 vs HeLa. A.** Principal Component Analysis (PCA) of the top 500 transcripts with the highest variance across all samples, the two Principal Components, PC1 and PC2 - that explain the most variance in the dataset are plotted against each other. Each dot corresponds to one sample and is colored by the respective condition. **B.** Sample2sample plot. The euclidean distance between the transcript expression patterns of all samples to each other is plotted using heatmap. Normalized gene counts were used for distance computation. **C.** Volcano plots. The differential transcript expression analysis results from DESeq2 are shown by plotting the log2FoldChange between the conditions of interest against the –log10 adjusted p-value per gene observed from the Wald-Test integrated in DESeq2. The top 10 significant genes are labeled by their symbol. **D.** Heatmaps. The expression of the top 20 differentially expressed transcripts is plotted using Heatmap by coloring the number of reads per transcript. The legend on the right side describes which colors correspond to highly and lowly expressed transcripts

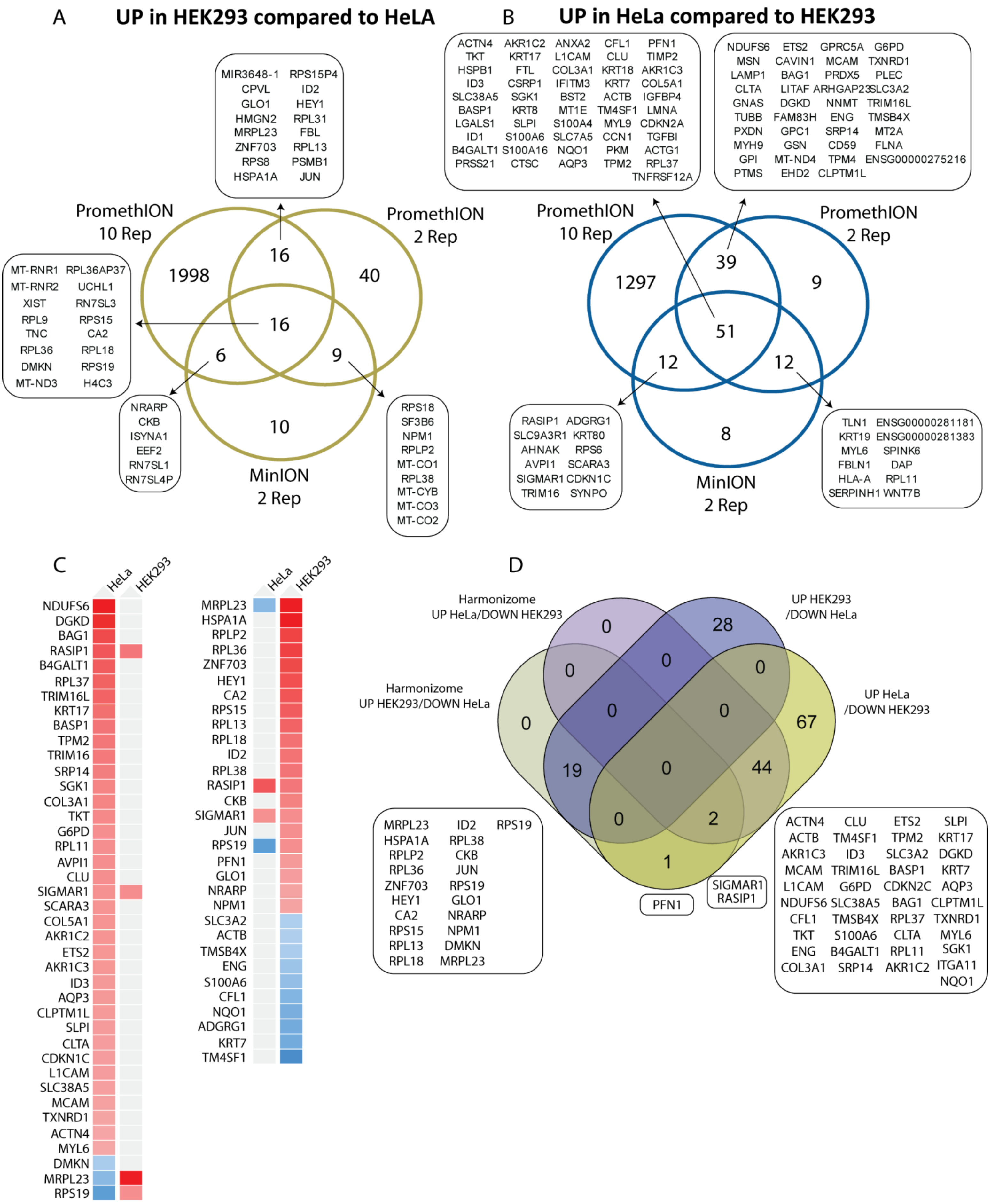
A
UP in HEK293 compared to HeLA
MIR3648-1 RPS15P4
CPVL ID2
GLO1 HEY1
HMGN2 RPL31
MRPL23 FBL
ZNF703 RPL13
RPS8 PSMB1
HSPA1A JUN
PromethION 10 Rep
PromethION 2 Rep
MinION 2 Rep
1998
16
40
16
6
9
10
MT-RNR1 RPL36AP37
MT-RNR2 UCHL1
XIST RN7SL3
RPL9 RPS15
TNC CA2
RPL36 RPL18
DMKN RPS19
MT-ND3 H4C3
NRARP
CKB
ISYNA1
EEF2
RN7SL1
RN7SL4P
RPS18
SF3B6
NPM1
RPLP2
MT-CO1
RPL38
MT-CYB
MT-CO3
MT-CO2
B
UP in HeLa compared to HEK293
ACTN4 AKR1C2 ANXA2 CFL1 PFN1
TKT KRT17 L1CAM CLU TIMP2
HSPB1 FTL COL3A1 KRT18 AKR1C3
ID3 CSRP1 IFITM3 KRT7 COL5A1
SLC38A5 SGK1 BST2 ACTB IGFBP4
BASP1 KRT8 MT1E TM4SF1 LMNA
LGALS1 SLPI S100A4 MYL9 CDKN2A
ID1 S100A6 SLC7A5 CCN1 TGFBI
B4GALT1 S100A16 NQO1 PKM ACTG1
PRSS21 CTSC AQP3 TPM2 RPL37
TNFRSF12A
NDUFS6 ETS2 GPRC5A G6PD
MSN CAVIN1 MCAM TXNRD1
LAMP1 BAG1 PRDX5 PLEC
CLTA LITAF ARHGAP23 SLC3A2
GNAS DGKD NNMT TRIM16L
TUBB FAM83H ENG TMSB4X
PXDN GPC1 SRP14 MT2A
MYH9 GSN CD59 FLNA
GPI MT-ND4 TPM4 ENSG00000275216
PTMS EHD2 CLPTM1L
PromethION 10 Rep
PromethION 2 Rep
MinION 2 Rep
1297
39
9
51
12
12
8
RASIP1 ADGRG1
SLC9A3R1 KRT80
AHNAK RPS6
AVPI1 SCARA3
SIGMAR1 CDKN1C
TRIM16 SYNPO
TLN1 ENSG00000281181
KRT19 ENSG00000281383
MYL6 SPINK6
FBLN1 DAP
HLA-A RPL11
SERPINH1 WNT7B
C
HeLa HEK293
NDUFS6
DGKD
BAG1
RASIP1
B4GALT1
RPL37
TRIM16L
KRT17
BASP1
TPM2
TRIM16
SRP14
SGK1
COL3A1
TKT
G6PD
RPL11
AVPI1
CLU
SIGMAR1
SCARA3
COL5A1
AKR1C2
ETS2
AKR1C3
ID3
AQP3
CLPTM1L
SLPI
CLTA
CDKN1C
L1CAM
SLC38A5
MCAM
TXNRD1
ACTN4
MYL6
DMKN
MRPL23
RPS19
HeLa HEK293
MRPL23
HSPA1A
RPLP2
RPL36
ZNF703
HEY1
CA2
RPS15
RPL13
RPL18
ID2
RPL38
RASIP1
CKB
SIGMAR1
JUN
RPS19
PFN1
GLO1
NRARP
NPM1
SLC3A2
ACTB
TMSB4X
ENG
S100A6
CFL1
NQO1
ADGRG1
KRT7
TM4SF1
D
Harmonizome UP HeLa/DOWN HEK293
UP HEK293 /DOWN HeLa
Harmonizome UP HEK293/DOWN HeLa
UP HeLa /DOWN HEK293
0
28
0
0
0
0
0
0
67
19
0
44
0
2
1
MRPL23 ID2 RPS19
HSPA1A RPL38
RPLP2 CKB
RPL36 JUN
ZNF703 RPS19
HEY1 GLO1
CA2 NRARP
RPS15 NPM1
RPL13 DMKN
RPL18 MRPL23
PFN1
SIGMAR1
RASIP1
ACTN4 CLU ETS2 SLPI
ACTB TM4SF1 TPM2 KRT17
AKR1C3 ID3 SLC3A2 DGKD
MCAM TRIM16L BASP1 KRT7
L1CAM G6PD CDKN2C AQP3
NDUFS6 SLC38A5 BAG1 CLPTM1L
CFL1 TMSB4X RPL37 TXNRD1
TKT S100A6 CLTA MYL6
ENG B4GALT1 RPL11 SGK1
COL3A1 SRP14 AKR1C2 ITGA11
NQO1

**Supplementary Figure 10. Comparative gene expression analysis between HEK293 and HeLa cells across PromethION and MinION sequencing platforms. A-B.**Venn diagrams showing genes upregulated in HEK293 compared to HeLa (A) and in HeLa compared to HEK293 (B) across PromethION (10-replicate and 2-replicate setups) and MinION (2-replicate setup). Overlaps indicate genes consistently detected across platforms, highlighting shared and unique DEGs. **C.** Heatmaps showing the relative expression levels of selected DEGs enriched in HEK293 or HeLa according to through Harmonizome database (Rouillard et al. 2016). The Harmonizome database contains various datasets, including the "HPA Cell Line Gene Expression Profiles" which evaluates differential gene expression across distinct cell lines (Uhlén et al. 2015). We examined whether the DEGs aligned with the observations in the database. **D**. Venn diagram comparing DEGs identified in HEK293 and HeLa against the Harmonizome database. Overlapping regions validate the observed transcriptional profiles, confirming consistency with known cell line gene expression patterns.
