## Supplementary Document S2 for "Real-time transcriptomic profiling in distinct experimental conditions"

### Supplementary Figures 11-20, related to Figure 3

#### Experimental enriched transcript expression


**Supplementary Figure 11 |** Sequencing overview in RiboM/RiboP/TotalR. ....................2
**Supplementary Figure 12 |** General overview RiboM vs TotalR. ....................4
**Supplementary Figure 13 |** Gene expression RiboM vs TotalR. ....................6
**Supplementary Figure 14 |** Transcript expression RiboM vs TotalR. ....................8
**Supplementary Figure 15 |** General overview RiboP vs TotalR. ....................9
**Supplementary Figure 16 |** Gene expression RiboP vs TotalR. ....................11
**Supplementary Figure 17 |** Transcript expression RiboP vs TotalR. ....................13
**Supplementary Figure 18 |** General overview RiboP vs RiboM. ....................14
**Supplementary Figure 19 |** Gene expression RiboP vs RiboM. ....................16
**Supplementary Figure 20 |** Transcript expression RiboP vs RiboM ....................18


#### *General supplementary figures description*

**Sequencing overview** – Refers to sequencing metrics including total reads generated per barcode/sample, mapped reads, gene counts and transcript counts.
**General overview –** Refers to data collected from NanopoReaTA regaring the general sample/condition overview including: Read length overview (per sample and condition), gene expression variability (per sample and condition), change in gene composition (per sample and condition) and Process time.
**Gene expression** - Refers to data collected from NanopoReaTA regaring differential gene expression analysis including: Principal Component Analysis (PCA), Volcano plots, Sample-to-sample variaiblity and Heatmaps.
**Transcript expression** - Refers to data collected from NanopoReaTA regaring differential transcript expression analysis including: Principal Component Analysis (PCA), Volcano plot, Sample-to-sample variaiblity and Heatmaps.

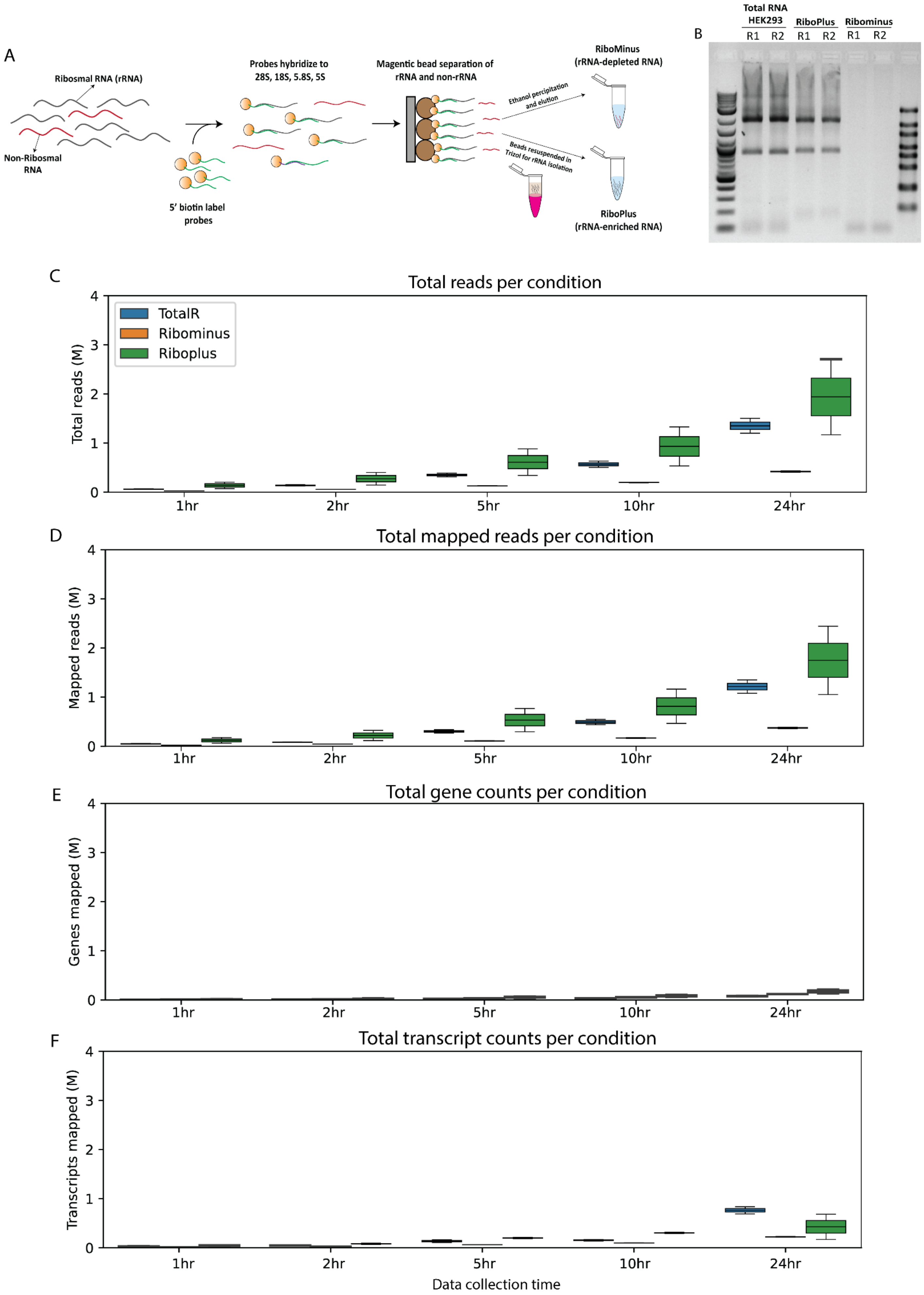
A
Ribosmal RNA (rRNA)
Non-Ribosmal RNA
5' biotin label probes
Probes hybridize to 28S, 18S, 5.8S, 5S
Magentic bead separation of rRNA and non-rRNA
Ethanol percipitation and elution
Beads resuspended in Trizol for rRNA isolation
RiboMinus (rRNA-depleted RNA)
RiboPlus (rRNA-enriched RNA)
B
Total RNA HEK293
RiboPlus
Ribominus
R1
R2
C
Total reads per condition
TotalR
Ribominus
Riboplus
Total reads (M)
1hr
2hr
5hr
10hr
24hr
D
Total mapped reads per condition
Mapped reads (M)
E
Total gene counts per condition
Genes mapped (M)
F
Total transcript counts per condition
Transcripts mapped (M)
Data collection time

**Supplementary Figure 11. Selective purification ribosomal-depleted (RiboMinus) and ribosomal-enriched (RiboPlus) transcripts and sequencing overview. A.** Selective purification of distinct RNA populations was performed using the Ribominus™ Eukaryote kit for RNA-seq (#Ambion, A10837-08) according to the manufacturer's "standard protocol" instructions. Total RNA (containing rRNA and other RNA transcripts) is incubated with 5' biotin lable probes complementary to rRNA RNA transcripts (28S, 18S, 5.8S and 5S). Probes hybrdize to rRNA and seperated using magentic bead seperation. Non-rRNA are purified using ethanol percipitation whereas bead-bound rRNA are purified using Trizol RNA isolation. B. 1 µg of Total RNA (HEK293) and RiboPlus and 150ng of Ribominus were loaded on 1% TBE agarose gel stained with ethidium bromide (in dublicates). Both total RNA and RiboPlus (rRNA-enriched) contained bands corresponding to 28S, 18S and 5S rRNA wheras RiboMinus contained a band correspoing to lower fragment length. **C-F.** The box plot represent total read generated (**C**), total mapped reads (**D**), total gene counts (**E**) and total transcript count (**F**) per condition (from 2 replicates), per time point.

### General overview
### Ribominus vs TotalR

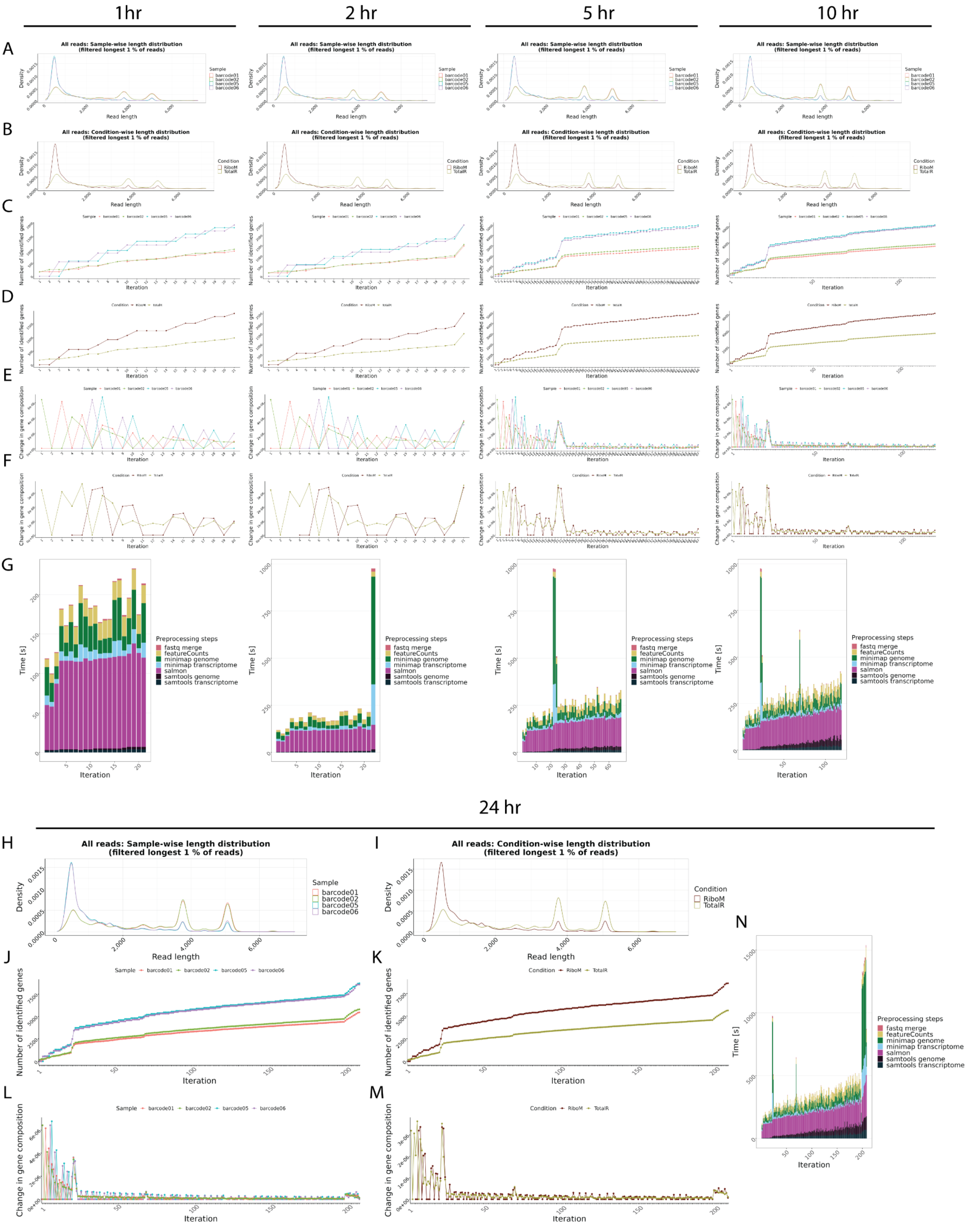

**Supplementary Figure 12. General overview in RiboMinus vs Total RNA**. **A-B**. Read length overview. The distribution of read lengths derived from generated fastq files is plotted per sample (A) and per condition (B). All reads of length over the 99 % quantile of all lengths are removed from these visualizations. **C-D**. Gene expression variability. The number of identified genes (> 0 reads counted) is plotted after each iteration per sample (C) and per condition (D). When the detection of additional genes finishes, the lines reach a plateau. **E-F**. Change in gene composition. The change in gene composition (CGC) plot shows the sum of absolute differences of relative gene (%gi) counts per individual gene (g) in iteration (i) to the relative gene counts in iteration (i-1) for each respective sample (E) and condition (F). CGC = sum(|(%g(i-1))-(%g(i))|). **G.** Process time. The bar plots show the time in seconds all preprocessing steps needed per iteration. One iteration process at maximum 30 files from all samples. This plot updates automatically as a new process finish. **H-N**. Final general overview. Read length (H-I), Gene expression variability (J-K), Change in gene composition (L-M), and processing time (N) for the 24hr time point. All plots are organized according to their respective time points of collection.

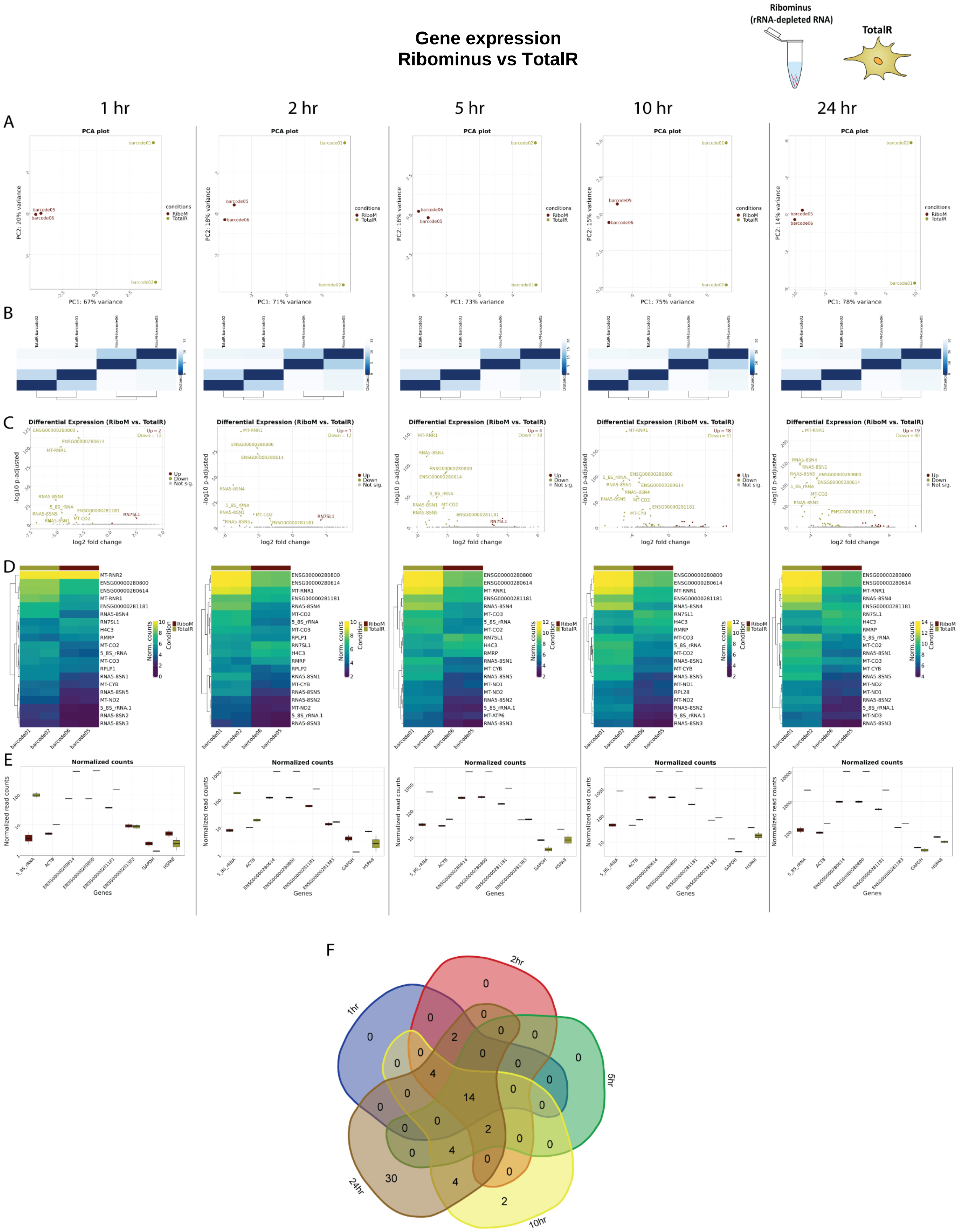

Gene expression
Ribominus vs TotalR
Ribominus
(rRNA-depleted RNA)
TotalR
1 hr
2 hr
5 hr
10 hr
24 hr
A
PCA plot
PC1: 67% variance
PC1: 71% variance
PC1: 73% variance
PC1: 75% variance
PC1: 78% variance
conditions
RiboM
TotalR
B
C
Differential Expression (RiboM vs. TotalR)
log2 fold change
-log10 p-adjusted
Up
Down
Not sig.
D
Norm. counts
Condition
E
Normalized counts
Normalized read counts
Genes
F
1hr
2hr
5hr
10hr
24hr
0
0
0
0
2
0
0
0
0
0
0
4
0
14
0
0
0
0
0
0
2
0
0
0
4
0
0
30
4
2

**Supplementary Figure 13. Gene expression analysis in RiboMinus vs Total RNA. A.** Principal Component Analysis (PCA) of the top 500 genes with the highest variance across all samples, the two Principal Components. Each dot corresponds to one sample and is colored by the respective condition. **B.** Sample2sample distance plot. The euclidean distance between the gene expression patterns of all samples to each other is plotted using heatmap. Normalized gene counts were used for distance computation. **C.** Volcano plots. The differential gene expression analysis results from DESeq2 are shown by plotting the log2FoldChange between the conditions of interest against the –log10 adjusted p-value per gene observed from the Wald-Test integrated in DESeq2 [Love, Huber, and Anders et al. 2014]. The top 10 significant genes are labeled by their symbol. **D.** Heatmaps. The expression of the top 20 differentially expressed genes is plotted using Heatmap by coloring the number of reads per gene. The legend on the right side describes which colors correspond to highly and lowly expressed genes. **E**. Normalized gene count for selected genes. Normalized gene counts from FeatureCounts [Liao et al. 2014] are visualized for selected genes per condition using boxplots. The median-of-ratios normalization method from DESeq2 [Love et al. 2014] was used for normalization. **F**. 5-way Venn diagram showing the differentially expressed gene overlaps between the distinct collected time points.

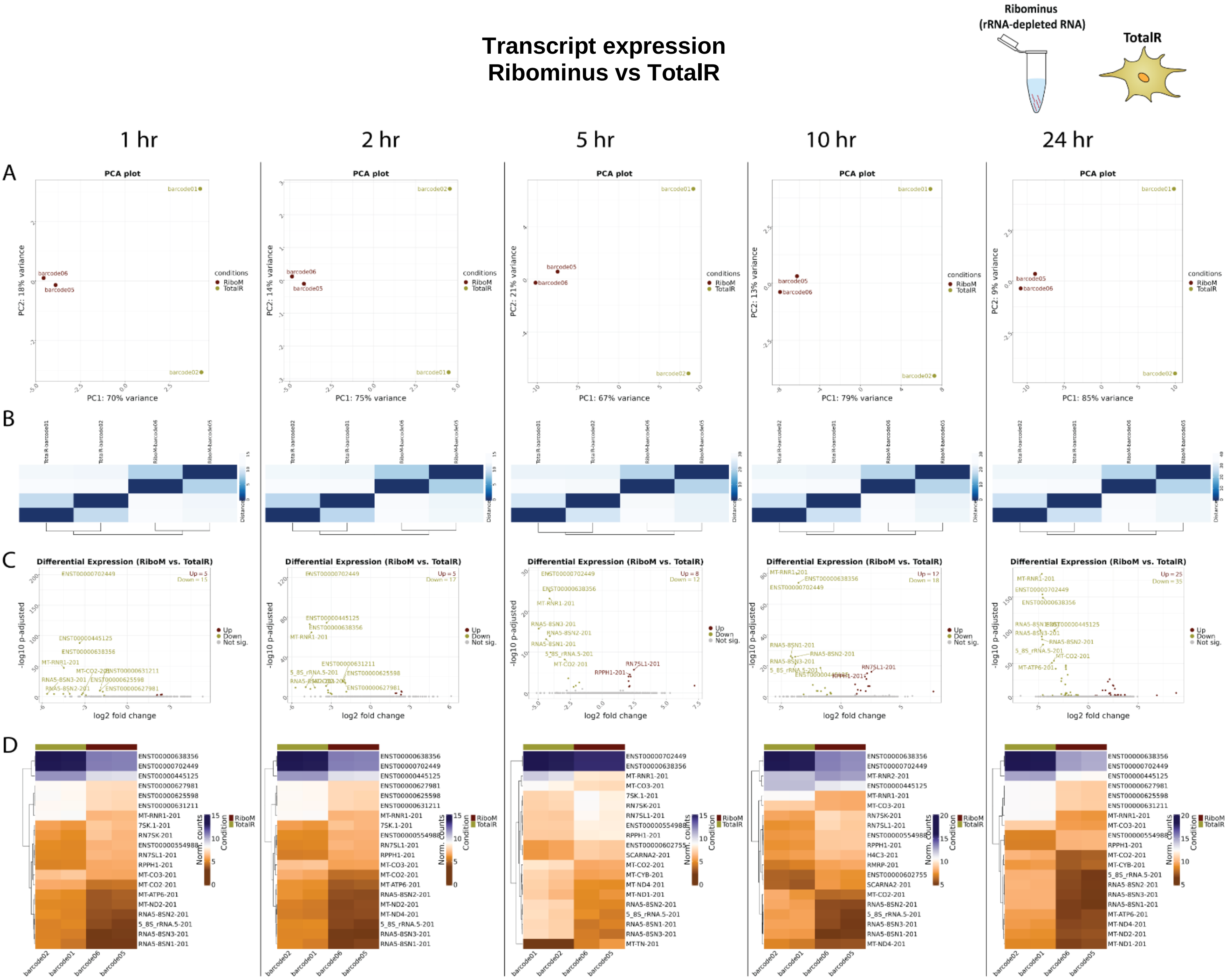


**Supplementary Figure 14. Transcript expression in RiboMinus vs Total RNA. A.** Principal Component Analysis (PCA) of the top 500 transcripts with the highest variance across all samples, the two Principal Components, PC1 and PC2 - that explain the most variance in the dataset are plotted against each other. Each dot corresponds to one sample and is colored by the respective condition. **B.** Sample2sample plot. The euclidean distance between the transcript expression patterns of all samples to each other is plotted using heatmap. Normalized gene counts were used for distance computation. **C.** Volcano plots. The differential transcript expression analysis results from DESeq2 are shown by plotting the log2FoldChange between the conditions of interest against the –log10 adjusted p-value per gene observed from the Wald-Test integrated in DESeq2. The top 10 significant genes are labeled by their symbol. **D.** Heatmaps. The expression of the top 20 differentially expressed transcripts is plotted using Heatmap by coloring the number of reads per transcript. The legend on the right side describes which colors correspond to highly and lowly expressed transcripts.

### General overview
### Riboplus vs TotalR

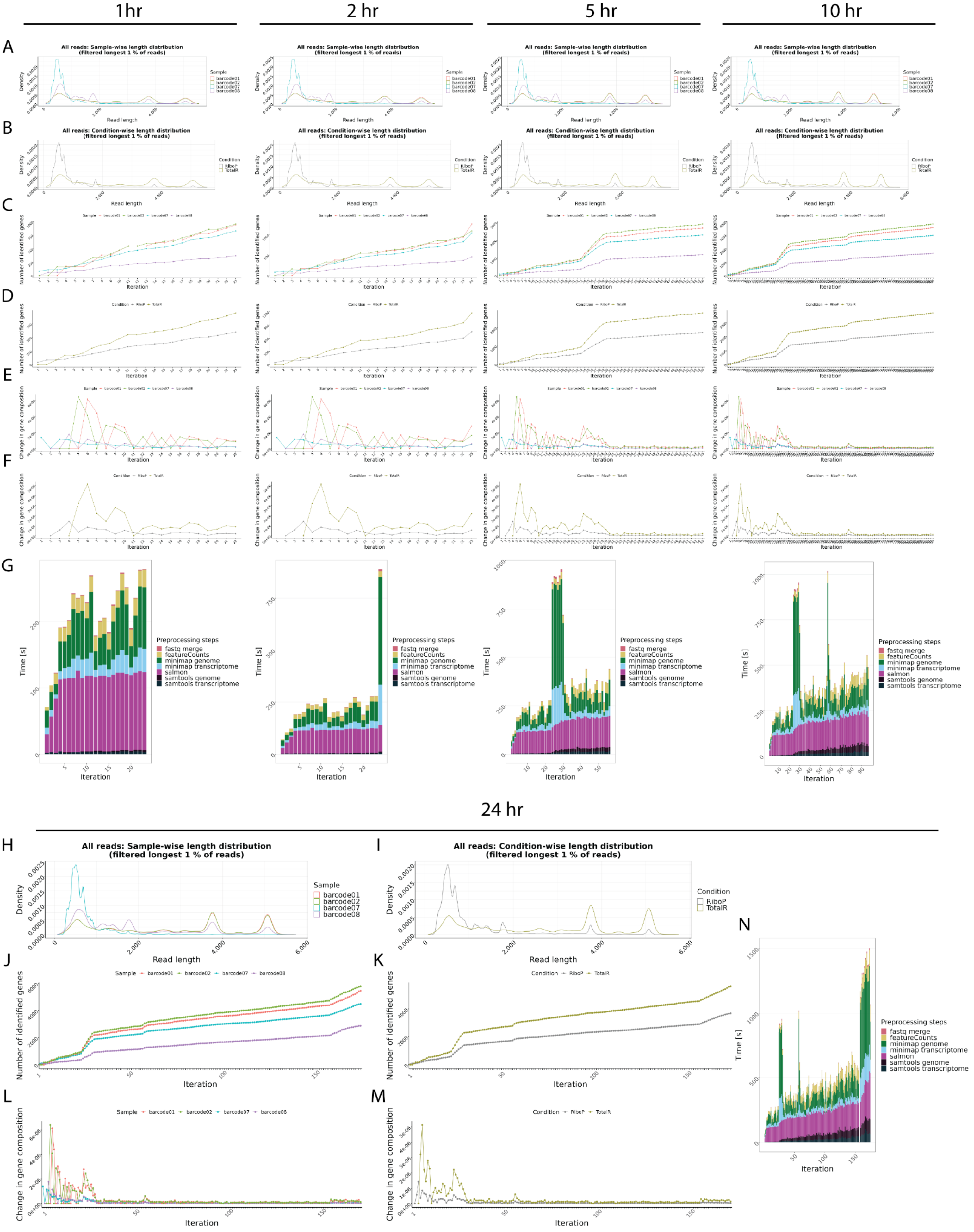

**Supplementary Figure 15. General overview in RiboPlus vs Total RNA**. **A-B**. Read length overview. The distribution of read lengths derived from generated fastq files is plotted per sample (A) and per condition (B). All reads of length over the 99 % quantile of all lengths are removed from these visualizations. **C-D**. Gene expression variability. The number of identified genes (> 0 reads counted) is plotted after each iteration per sample (C) and per condition (D). When the detection of additional genes finishes, the lines reach a plateau. **E-F**. Change in gene composition. The change in gene composition (CGC) plot shows the sum of absolute differences of relative gene (%gi) counts per individual gene (g) in iteration (i) to the relative gene counts in iteration (i-1) for each respective sample (E) and condition (F). CGC = sum(|(%g(i-1))-(%g(i))|). **G.** Process time. The bar plots show the time in seconds all preprocessing steps needed per iteration. One iteration process at maximum 30 files from all samples. This plot updates automatically as a new process finish. **H-N**. Final general overview. Read length (H-I), Gene expression variability (J-K), Change in gene composition (L-M), and processing time (N) for the 24hr time point. All plots are organized according to their respective time points of collection.

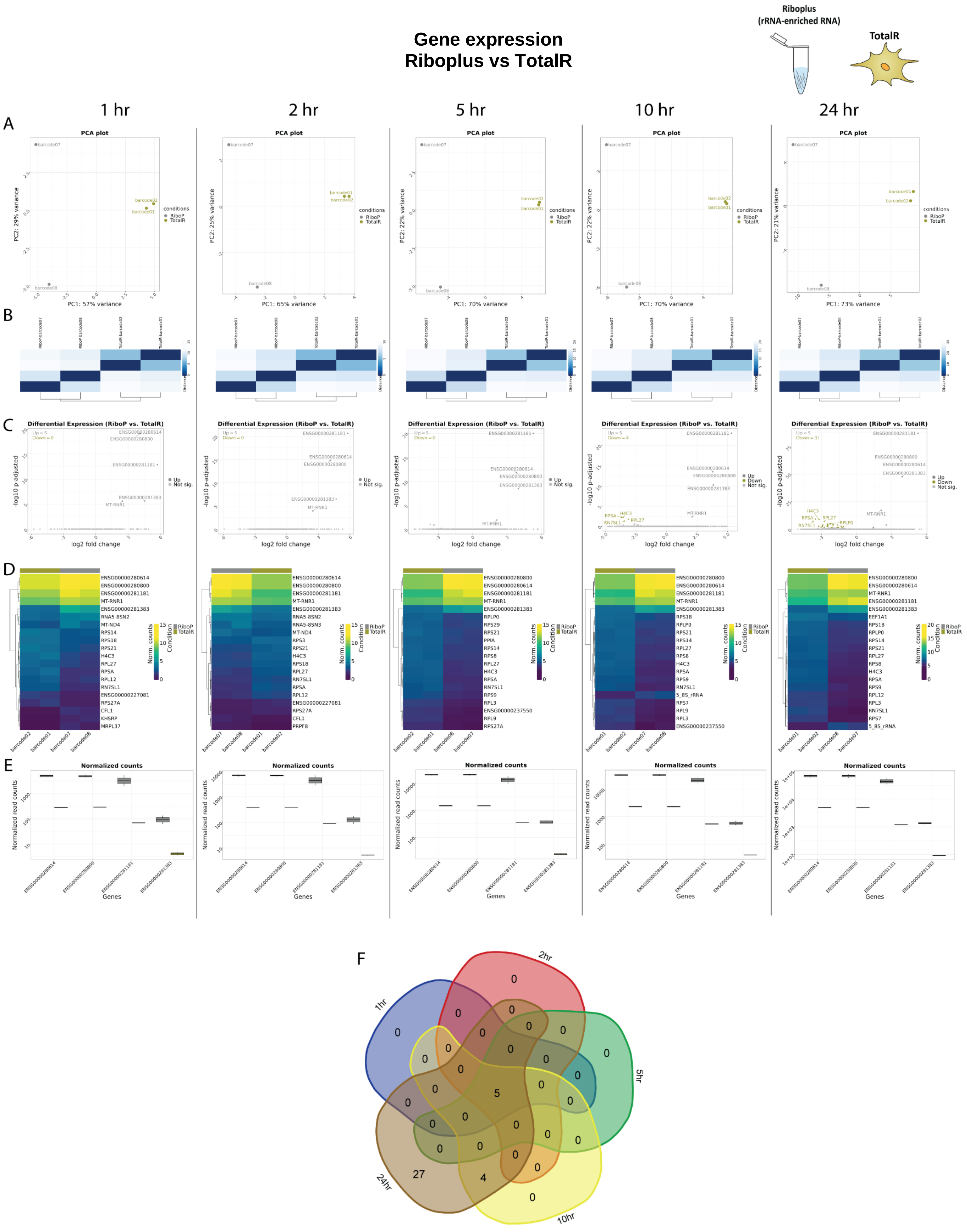
Gene expression
Riboplus vs TotalR
Riboplus
(rRNA-enriched RNA)
TotalR
1 hr
2 hr
5 hr
10 hr
24 hr
A
PCA plot
PC1: 57% variance
PC1: 65% variance
PC1: 70% variance
PC1: 70% variance
PC1: 73% variance
PC2: 29% variance
PC2: 25% variance
PC2: 22% variance
PC2: 22% variance
PC2: 21% variance
conditions
RiboP
TotalR
B
C
Differential Expression (RiboP vs. TotalR)
log2 fold change
-log10 p-adjusted
Up
Down
Not sig.
D
Norm. counts
Condition
RiboP
TotalR
E
Normalized counts
Normalized read counts
Genes
F
1hr
2hr
5hr
10hr
24hr

**Supplementary Figure 16. Gene expression analysis in RiboPlus vs Total RNA. A.** Principal Component Analysis (PCA) of the top 500 genes with the highest variance across all samples, the two Principal Components. Each dot corresponds to one sample and is colored by the respective condition. **B.** Sample2sample distance plot.The euclidean distance between the gene expression patterns of all samples to each other is plotted using heatmap. Normalized gene counts were used for distance computation. **C.** Volcano plots. The differential gene expression analysis results from DESeq2 are shown by plotting the log2FoldChange between the conditions of interest against the –log10 adjusted p-value per gene observed from the Wald-Test integrated in DESeq2 [Love, Huber, and Anders et al. 2014]. The top 10 significant genes are labeled by their symbol. **D.** Heatmaps. The expression of the top 20 differentially expressed genes is plotted using Heatmap by coloring the number of reads per gene. The legend on the right side describes which colors correspond to highly and lowly expressed genes. **E**. Normalized gene count for selected genes. Normalized gene counts from FeatureCounts [Liao et al. 2014] are visualized for selected genes per condition using boxplots. The median-of-ratios normalization method from DESeq2 [Love et al. 2014] was used for normalization. **F**. 5-way Venn diagram showing the differentially expressed gene overlaps between the distinct collected time points.

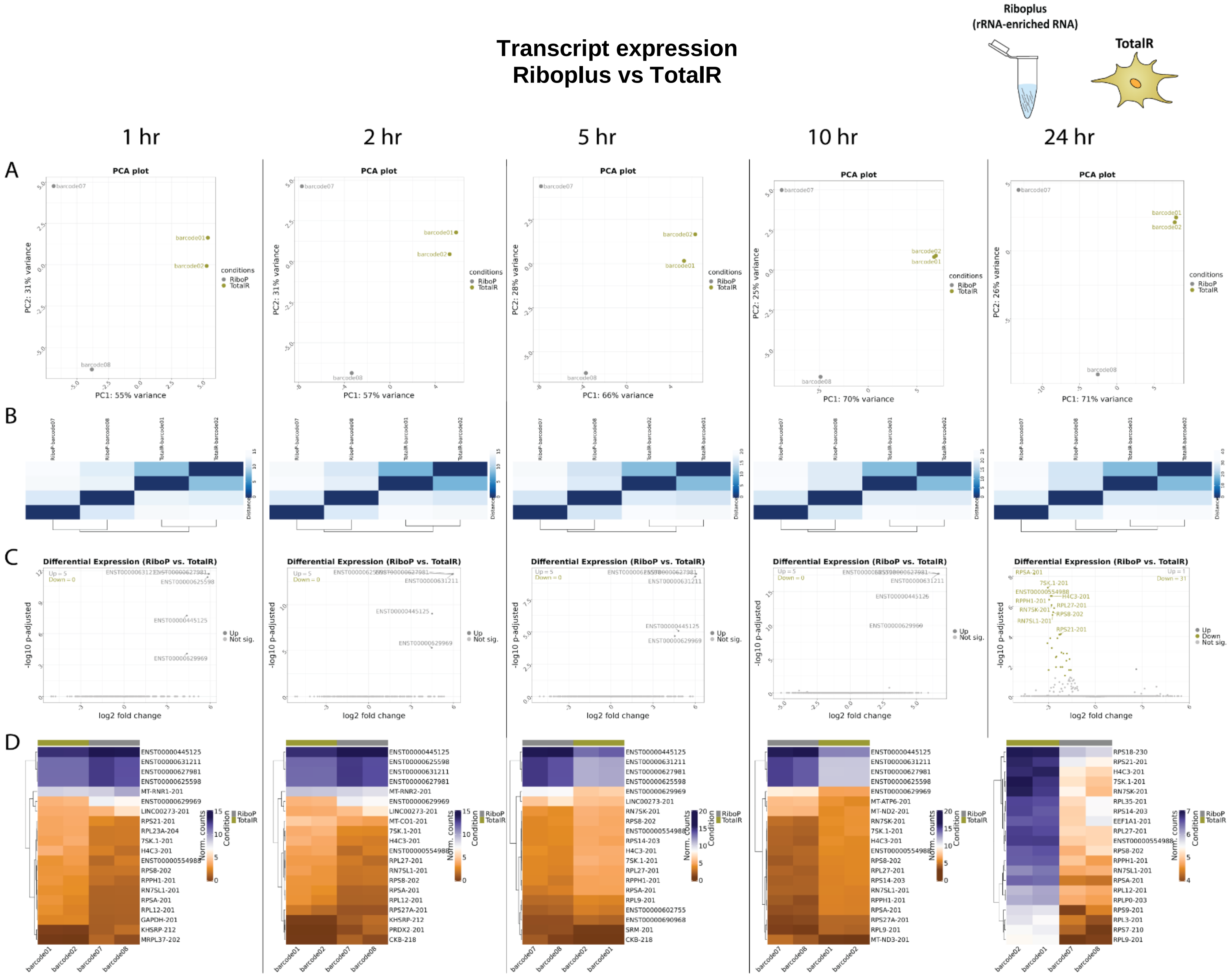


**Supplementary Figure 17. Transcript expression in RiboPlus vs Total RNA. A.** Principal Component Analysis (PCA) of the top 500 transcripts with the highest variance across all samples, the two Principal Components. Each dot corresponds to one sample and is colored by the respective condition. **B.** Sample2sample plot. The euclidean distance between the transcript expression patterns of all samples to each other is plotted using heatmap. Normalized gene counts were used for distance computation. **C.** Volcano plots. The differential transcript expression analysis results from DESeq2 are shown by plotting the log2FoldChange between the conditions of interest against the –log10 adjusted p-value per gene observed from the Wald-Test integrated in DESeq2. The top 10 significant genes are labeled by their symbol. **D.** Heatmaps. The expression of the top 20 differentially expressed transcripts is plotted using Heatmap by coloring the number of reads per transcript. The legend on the right side describes which colors correspond to highly and lowly expressed transcripts.

### General overview
### Riboplus vs Ribominus

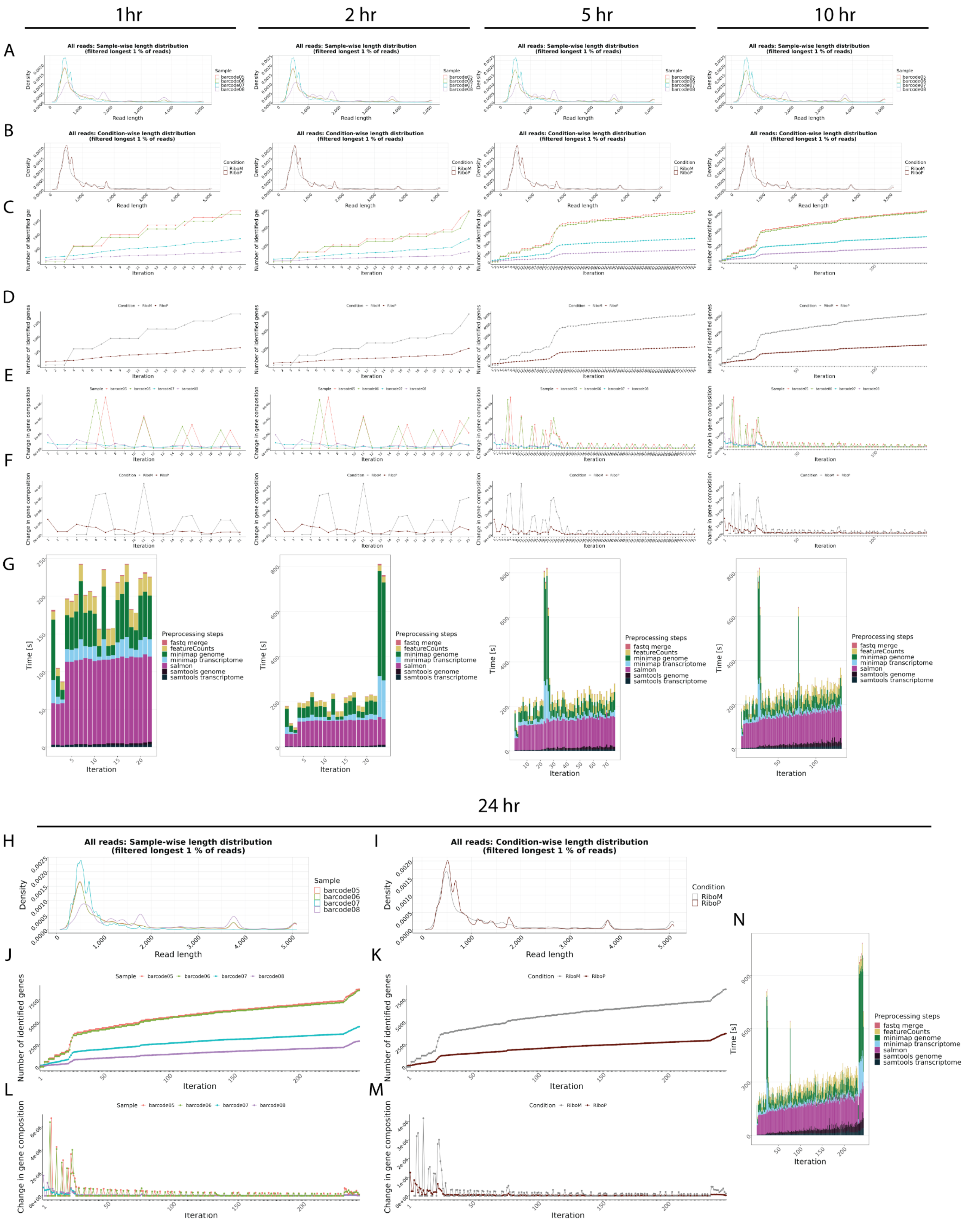

**Supplementary Figure 18. General overview in RiboPlus vs RiboMinus**. **A-B**. Read length overview. The distribution of read lengths derived from generated fastq files is plotted per sample (A) and per condition (B). All reads of length over the 99 % quantile of all lengths are removed from these visualizations. **C-D**. Gene expression variability. The number of identified genes (> 0 reads counted) is plotted after each iteration per sample (C) and per condition (D). When the detection of additional genes finishes, the lines reach a plateau. **E-F**. Change in gene composition. The change in gene composition (CGC) plot shows the sum of absolute differences of relative gene (%gi) counts per individual gene (g) in iteration (i) to the relative gene counts in iteration (i-1) for each respective sample (E) and condition (F). CGC = sum(|(%g(i-1))-(%g(i))|). **G.** Process time. The bar plots show the time in seconds all preprocessing steps needed per iteration. One iteration process at maximum 30 files from all samples. This plot updates automatically as a new process finish. **H-N**. Final general overview. Read length (H-I), Gene expression variability (J-K), Change in gene composition (L-M), and processing time (N) for the 24hr time point. All plots are organized according to their respective time points of collection.

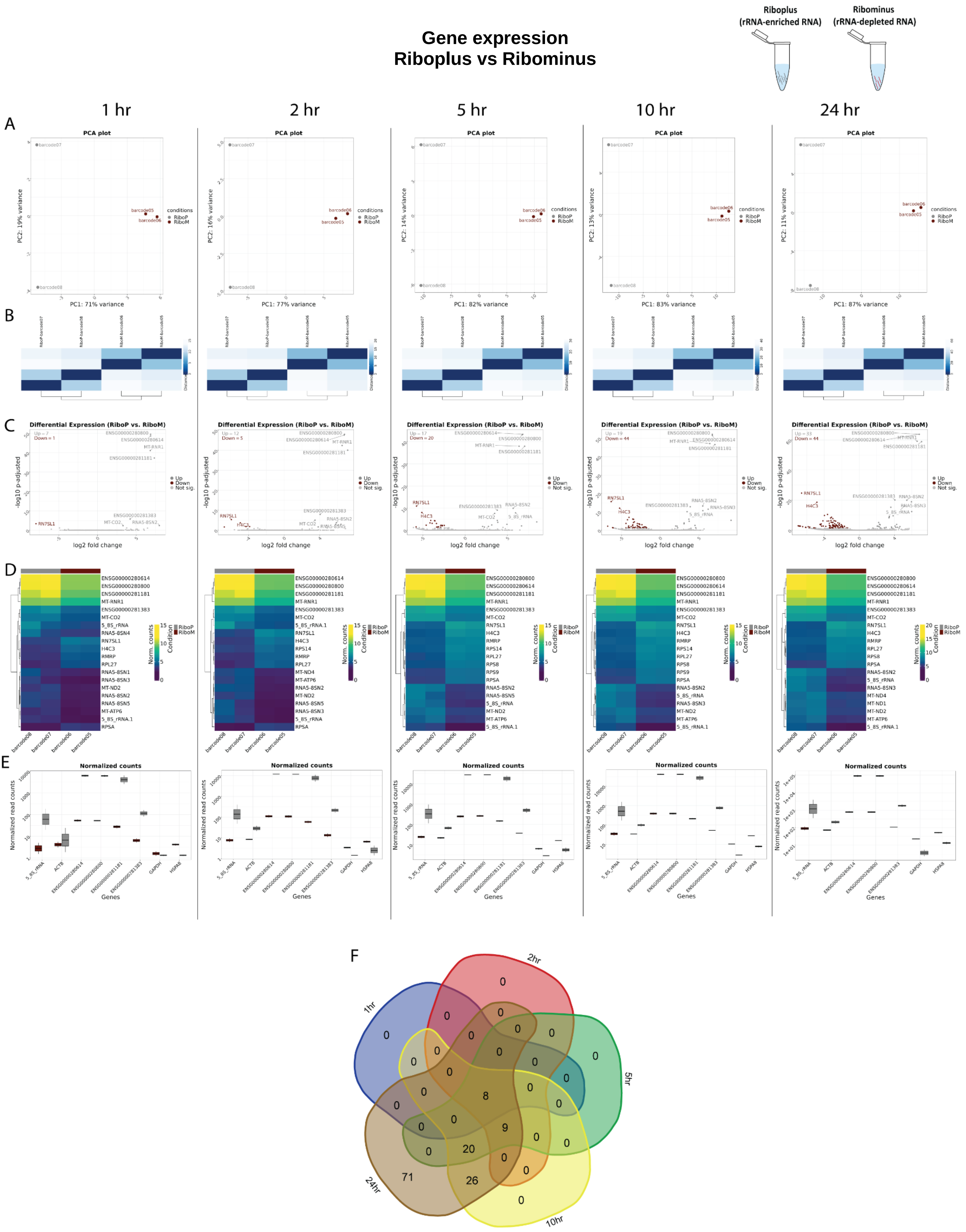
Gene expression
Riboplus vs Ribominus
Riboplus
(rRNA-enriched RNA)
Ribominus
(rRNA-depleted RNA)
1 hr
2 hr
5 hr
10 hr
24 hr
A
PCA plot
PC1: 71% variance
PC1: 77% variance
PC1: 82% variance
PC1: 83% variance
PC1: 87% variance
PC2: 19% variance
PC2: 16% variance
PC2: 14% variance
PC2: 13% variance
PC2: 11% variance
conditions
RiboP
RiboM
B
C
Differential Expression (RiboP vs. RiboM)
-log10 p-adjusted
log2 fold change
Up
Down
Not sig.
D
Norm. counts
Condition
E
Normalized counts
Normalized read counts
Genes
F
1hr
2hr
5hr
10hr
24hr

**Supplementary Figure 19. Gene expression analysis in RiboPlus vs RiboMinus. A.** Principal Component Analysis (PCA) of the top 500 genes with the highest variance across all samples, the two Principal Components. Each dot corresponds to one sample and is colored by the respective condition. **B.** Sample2sample distance plot. The euclidean distance between the gene expression patterns of all samples to each other is plotted using heatmap. Normalized gene counts were used for distance computation. **C.** Volcano plots. The differential gene expression analysis results from DESeq2 are shown by plotting the log2FoldChange between the conditions of interest against the –log10 adjusted p-value per gene observed from the Wald-Test integrated in DESeq2 [Love, Huber, and Anders et al. 2014]. The top 10 significant genes are labeled by their symbol. **D.** Heatmaps. The expression of the top 20 differentially expressed genes is plotted using Heatmap by coloring the number of reads per gene. The legend on the right side describes which colors correspond to highly and lowly expressed genes. **E**. Normalized gene count for selected genes. Normalized gene counts from FeatureCounts [Liao et al. 2014] are visualized for selected genes per condition using boxplots. The median-of-ratios normalization method from DESeq2 [Love et al. 2014] was used for normalization. **F**. 5-way Venn diagram showing the differentially expressed gene overlaps between the distinct collected time points.

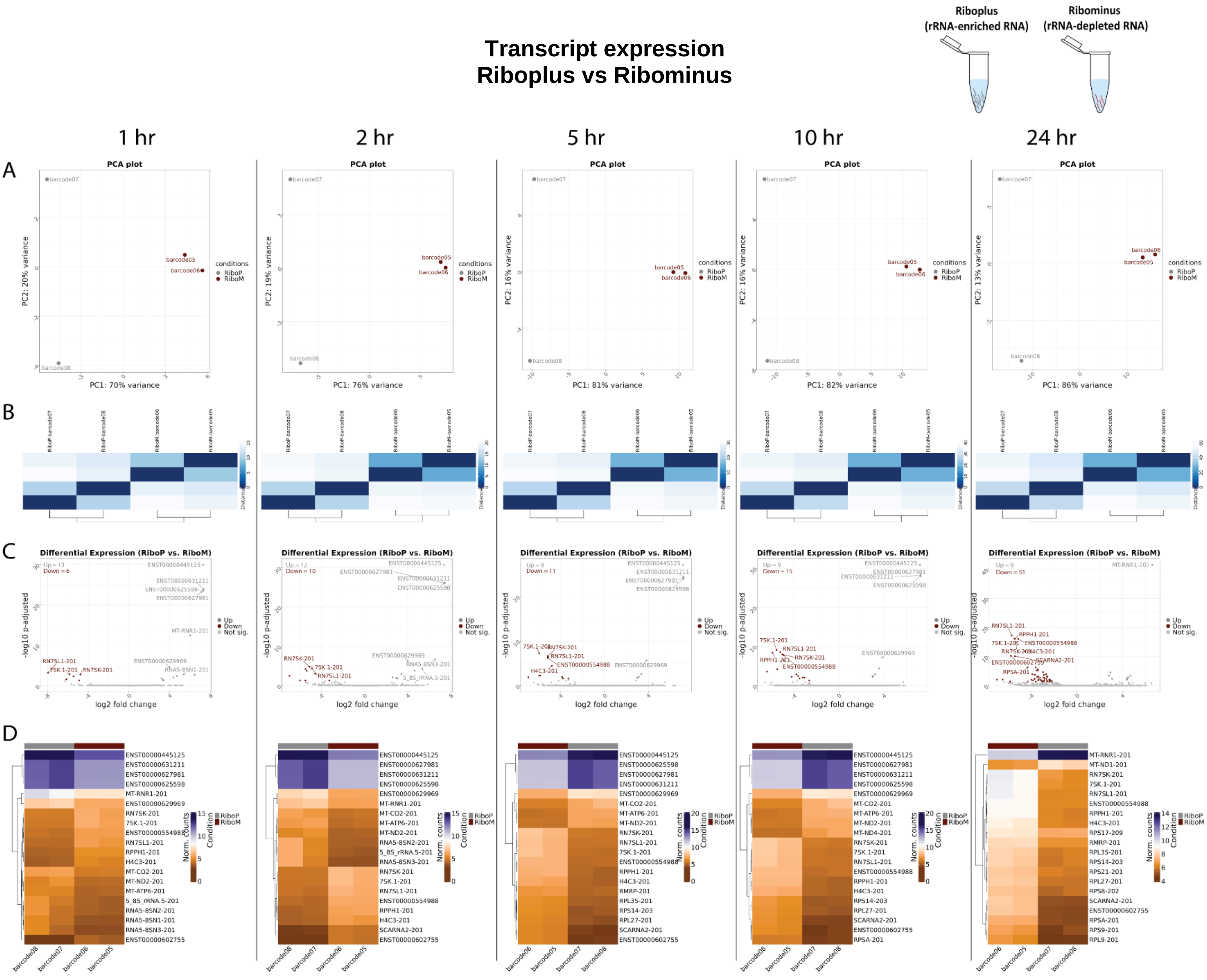


**Supplementary Figure 20. Transcript expression in RiboPlus vs RiboMinus. A.** Principal Component Analysis (PCA) of the top 500 transcripts with the highest variance across all samples, the two Principal Components. Each dot corresponds to one sample and is colored by the respective condition. **B.** Sample2sample plot. The euclidean distance between the transcript expression patterns of all samples to each other is plotted using heatmap. Normalized gene counts were used for distance computation. **C.** Volcano plots. The differential transcript expression analysis results from DESeq2 are shown by plotting the log2FoldChange between the conditions of interest against the –log10 adjusted p-value per gene observed from the Wald-Test integrated in DESeq2. The top 10 significant genes are labeled by their symbol. **D.** Heatmaps. The expression of the top 20 differentially expressed transcripts is plotted using Heatmap by coloring the number of reads per transcript. The legend on the right side describes which colors correspond to highly and lowly expressed transcripts.
