## Supplementary Document S3 for "Real-time transcriptomic profiling in distinct experimental conditions"

### Supplementary Figures 21-24, related to Figure 4

#### Heat-shock (HS) vs non-heat shock (NHS) comparison

**Supplementary Figure 21 |** Sequencing overview HS vs NHS. ....................2
**Supplementary Figure 22 |** PromethION - General overview HS vs NHS. ....................3
**Supplementary Figure 23 |** PromethION - Gene expression HS vs NHS. ....................5
**Supplementary Figure 24 |** PromethION - Transcript expression HS vs NHS. ....................7

#### *General supplementary figures description*

**Sequencing overview** – Refers to sequencing metrics including total reads generated per barcode/sample, mapped reads, gene counts and transcript counts.
**General overview –** Refers to data collected from NanopoReaTA regaring the general sample/condition overview including: Read length overview (per sample and condition), gene expression variability (per sample and condition), change in gene composition (per sample and condition) and Process time.
**Gene expression** - Refers to data collected from NanopoReaTA regaring differential gene expression analysis including: Principal Component Analysis (PCA), Volcano plots, Sample-to-sample variaiblity and Heatmaps.
**Transcript expression** - Refers to data collected from NanopoReaTA regaring differential transcript expression analysis including: Principal Component Analysis (PCA), Volcano plot, Sample-to-sample variaiblity and Heatmaps.

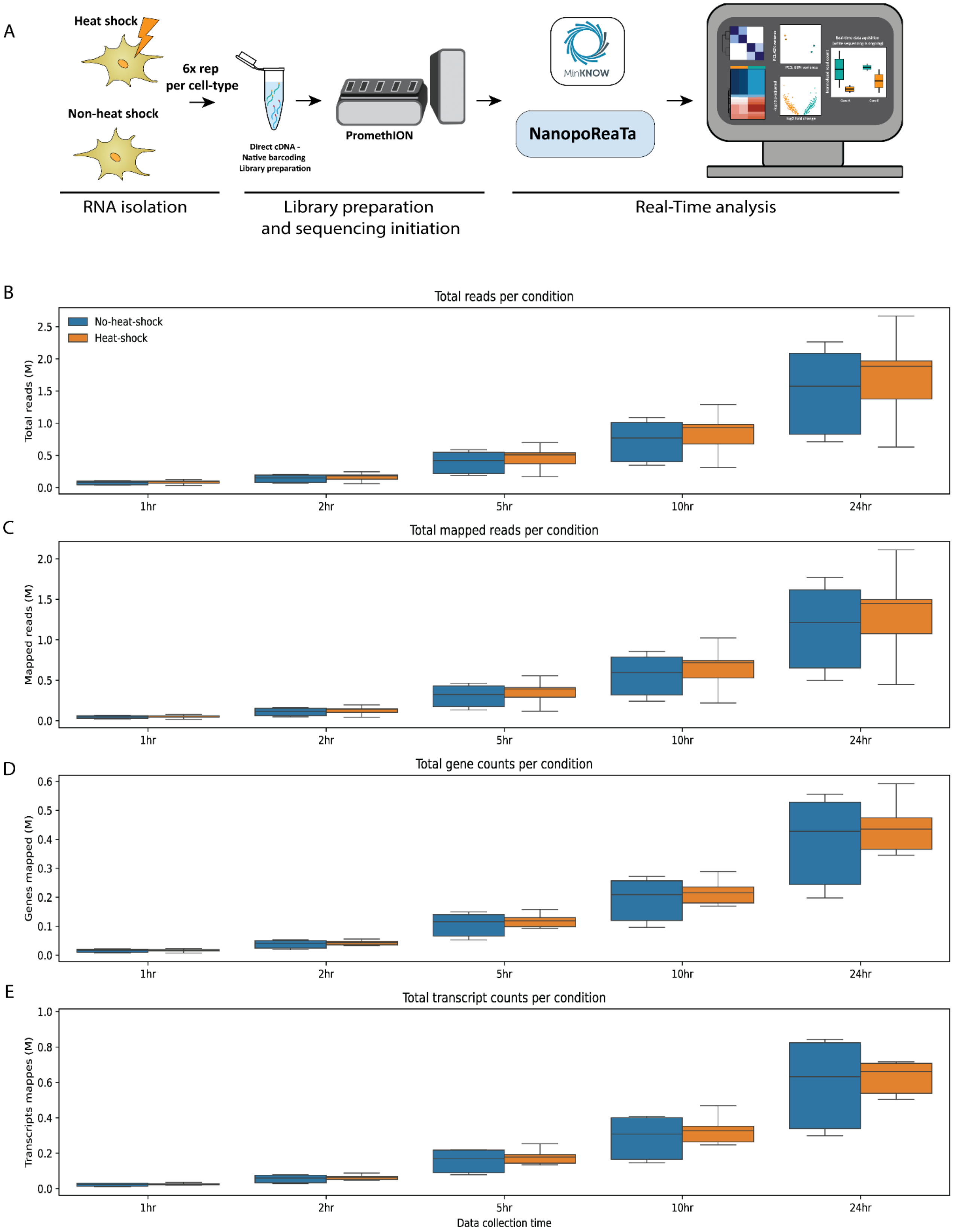


**Supplementary Figure 21. Experimental strategy of heat-shock experiments and sequencing overview. A.** RNA was isolated from HEK293 and HeLa cells (n=6) dscDNA library was prepared which included samples barcoding and adapter ligation. Samples were loaded and sequenced using a PromethION R10 flow cell or MinION R9 flow cell. NanopoReaTA was activated shortly after sequencing initiation and data was collected 1hr, 2hr, 5hr, 10hr, and 24hr post-sequencing initiation. **B-E.** The box plot represent total read generated (**B**), total mapped reads (**C**), total gene counts (**D**) and total transcript count (**E**) per condition (from 2 replicates), per time point.

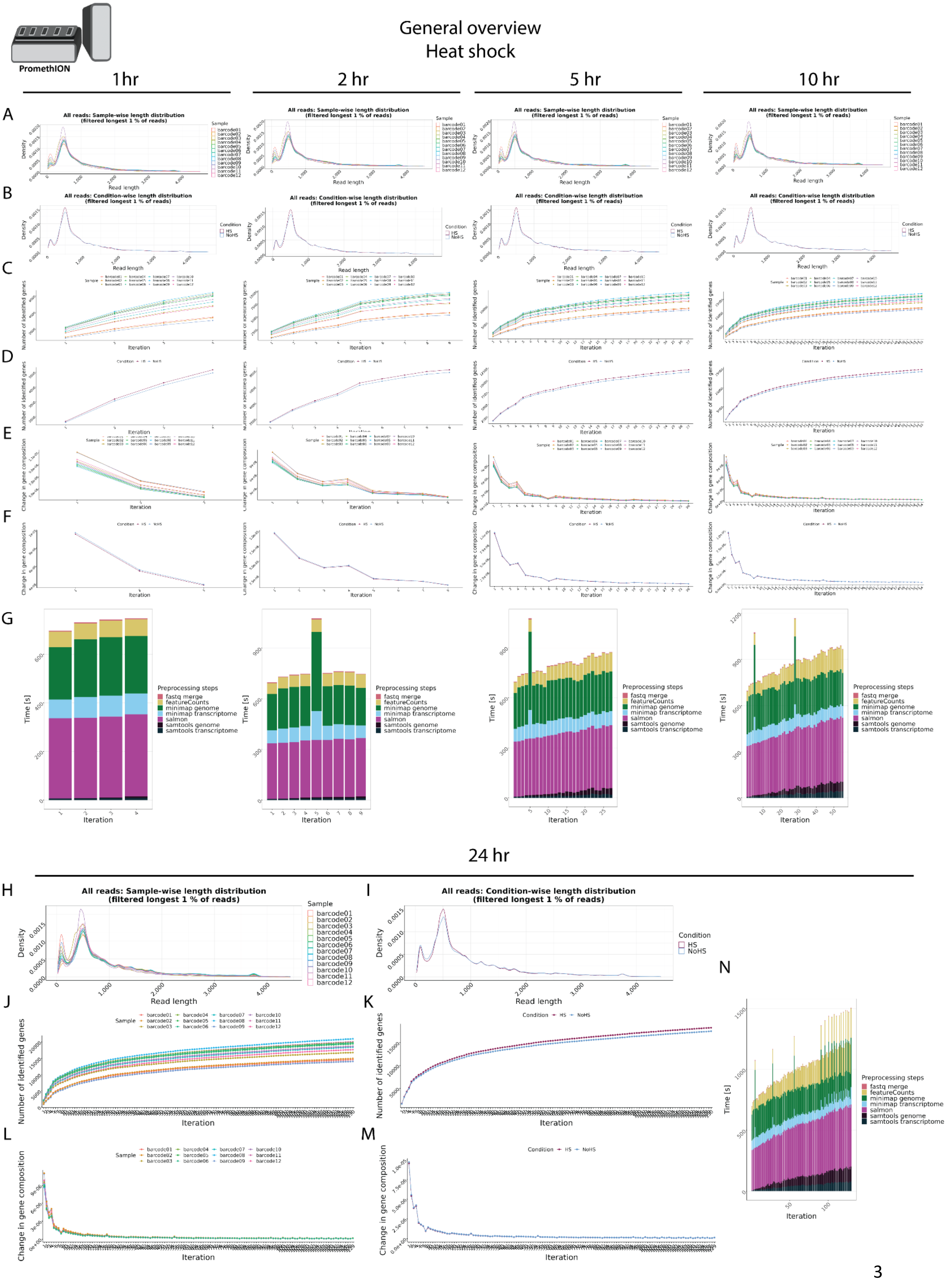
PromethION
General overview
Heat shock
1 hr
2 hr
5 hr
10 hr
24 hr
A
B
C
D
E
F
G
H
I
J
K
L
M
N
All reads: Sample-wise length distribution
(filtered longest 1 % of reads)
All reads: Condition-wise length distribution
(filtered longest 1 % of reads)
Density
Read length
Sample
barcode01
barcode02
barcode03
barcode04
barcode05
barcode06
barcode07
barcode08
barcode09
barcode10
barcode11
barcode12
Condition
HS
NoHS
Number of identified genes
Change in gene composition
Iteration
Time [s]
Preprocessing steps
fastq merge
featureCounts
minimap genome
minimap transcriptome
salmon
samtools genome
samtools transcriptome

**Supplementary Figure 22. General overview in HS vs NHS**. **A-B**. Read length overview. The distribution of read lengths derived from generated fastq files is plotted per sample (A) and per condition (B). All reads of length over the 99 % quantile of all lengths are removed from these visualizations. **C-D**. Gene expression variability. The number of identified genes (> 0 reads counted) is plotted after each iteration per sample (C) and per condition (D). When the detection of additional genes finishes, the lines reach a plateau. **E-F**. Change in gene composition. The change in gene composition (CGC) plot shows the sum of absolute differences of relative gene (%gi) counts per individual gene (g) in iteration (i) to the relative gene counts in iteration (i-1) for each respective sample (E) and condition (F). CGC = sum(|(%g(i-1))-(%g(i))|). **G.** Process time. The bar plots show the time in seconds all preprocessing steps needed per iteration. One iteration process at maximum 30 files from all samples. This plot updates automatically as a new process finish. **H-N**. Final general overview. Read length (H-I), Gene expression variability (J-K), Change in gene composition (L-M), and processing time (N) for the 24hr time point. All plots are organized according to their respective time points of collection.

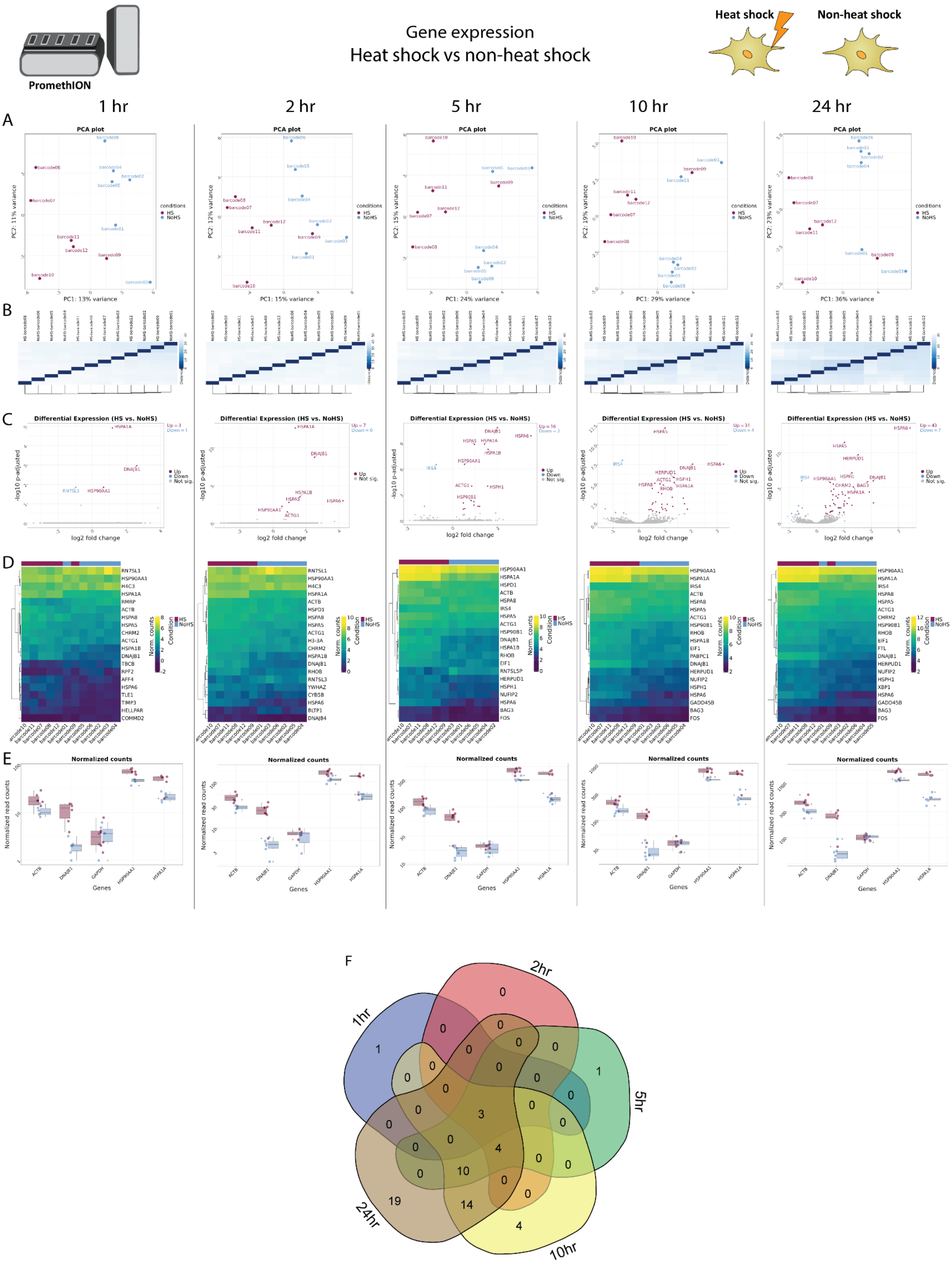
PromethION
Gene expression
Heat shock vs non-heat shock
Heat shock
Non-heat shock
1 hr
2 hr
5 hr
10 hr
24 hr
A
PCA plot
PC1: 13% variance
PC1: 15% variance
PC1: 24% variance
PC1: 29% variance
PC1: 36% variance
conditions
HS
NoHS
B
C
Differential Expression (HS vs. NoHS)
log2 fold change
-log10 p-adjusted
Up
Down
Not sig.
D
Norm. counts
Condition
E
Normalized counts
Normalized read counts
Genes
F
1hr
2hr
5hr
10hr
24hr

**Supplementary Figure 23. Gene expression analysis in in HS vs NHS. A.** Principal Component Analysis (PCA) of the top 500 genes with the highest variance across all samples, the two Principal Components. Each dot corresponds to one sample and is colored by the respective condition. **B.** Sample2sample distance plot. The euclidean distance between the gene expression patterns of all samples to each other is plotted using heatmap. Normalized gene counts were used for distance computation. **C.** Volcano plots. The differential gene expression analysis results from DESeq2 are shown by plotting the log2FoldChange between the conditions of interest against the –log10 adjusted p-value per gene observed from the Wald-Test integrated in DESeq2 [Love, Huber, and Anders et al. 2014]. The top 10 significant genes are labeled by their symbol. **D.** Heatmaps. The expression of the top 20 differentially expressed genes is plotted using Heatmap by coloring the number of reads per gene. The legend on the right side describes which colors correspond to highly and lowly expressed genes. **E**. Normalized gene count for selected genes. Normalized gene counts from FeatureCounts [Liao et al. 2014] are visualized for selected genes per condition using boxplots. The median-of-ratios normalization method from DESeq2 [Love et al. 2014] was used for normalization. **F**. 5-way Venn diagram showing the differentially expressed gene overlaps between the distinct collected time points.

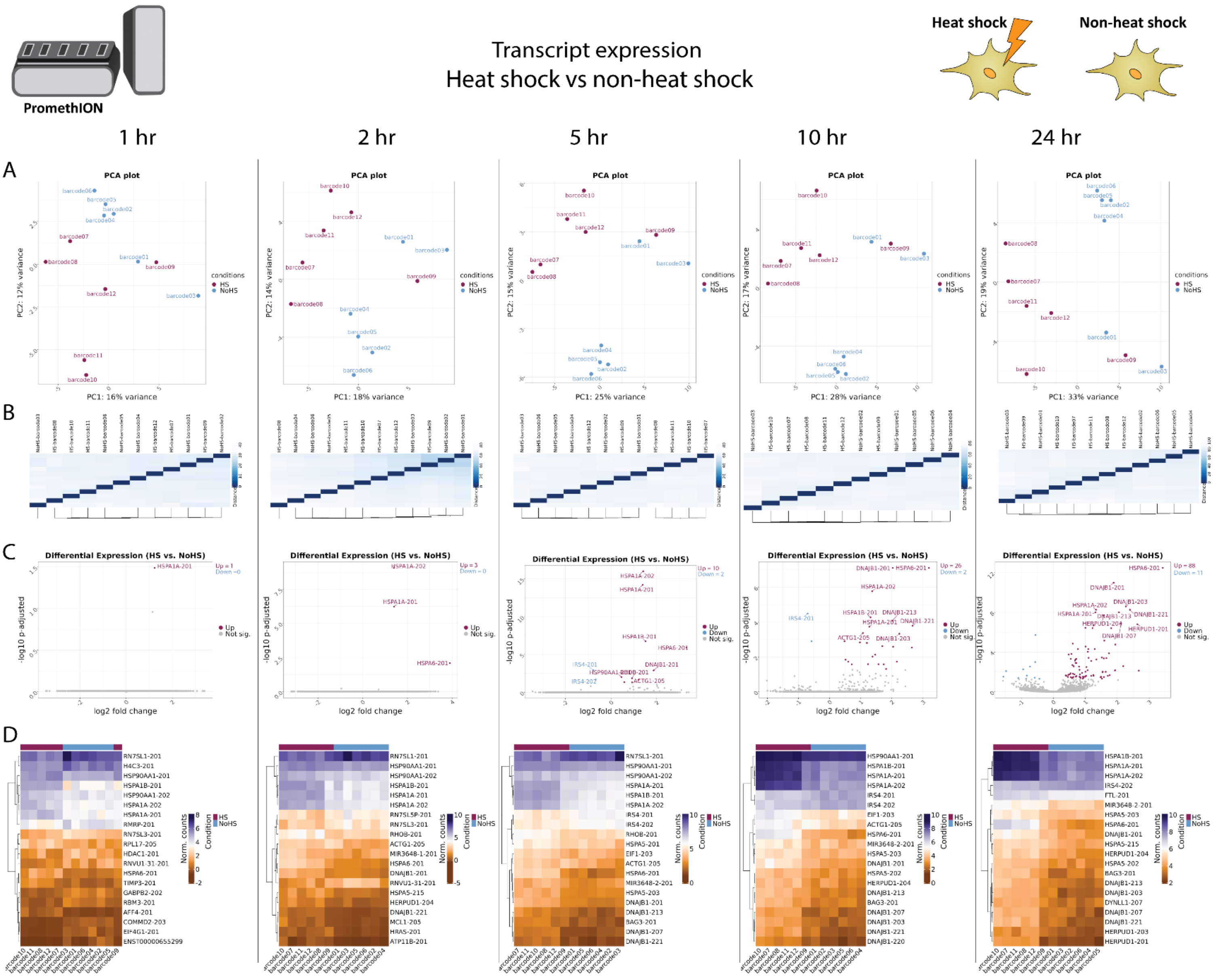


**Supplementary Figure 24. Transcript expression in HS vs NHS. A.** Principal Component Analysis (PCA) of the top 500 transcripts with the highest variance across all samples, the two Principal Components. Each dot corresponds to one sample and is colored by the respective condition. **B.** Sample2sample plot. The euclidean distance between the transcript expression patterns of all samples to each other is plotted using heatmap. Normalized gene counts were used for distance computation. **C.** Volcano plots. The differential transcript expression analysis results from DESeq2 are shown by plotting the log2FoldChange between the conditions of interest against the –log10 adjusted p-value per gene observed from the Wald-Test integrated in DESeq2. The top 10 significant genes are labeled by their symbol. **D.** Heatmaps. The expression of the top 20 differentially expressed transcripts is plotted using Heatmap by coloring the number of reads per transcript. The legend on the right side describes which colors correspond to highly and lowly expressed transcripts.
