## Supplementary Document S4 for "Real-time transcriptomic profiling in distinct experimental conditions"

### Supplementary Figures 25-41, related to Figure 5

#### Yeat Setup 1

**Supplementary Figure 25 |** Sequencing overview -Yeast setup 1.......................................................................2
**Supplementary Figure 26 |** Real-time transcriptomic analysis in yeast setup 1 samples using NanopoReaTA. ....4
**new1Δ-pEV(HIS3) vs WT-pEV(HIS3)**
**Supplementary Figure 27 |** General overview *new1Δ*-pEV*(HIS3) vs* WT-pEV*(HIS3)* ..............................................6
**Supplementary Figure 28 |** Gene expression *new1Δ*-pEV*(HIS3) vs* WT-pEV*(HIS3)*. ...............................................8
**Supplementary Figure 29 |** Transcript expression *new1Δ*-pEV*(HIS3) vs* WT-pEV*(HIS3)*. .....................................10
**WT-pNew1(HIS3) vs WT-pEV(HIS3)**
**Supplementary Figure 30 |** General overview WT-pNew1*(HIS3) vs* WT-pEV*(HIS3)*............................................11
**Supplementary Figure 31 |** Gene expression WT-pNew1*(HIS3) vs* WT-pEV*(HIS3)*. ...............................................13
**Supplementary Figure 32 |** Transcript expression WT-pNew1*(HIS3) vs* WT-pEV*(HIS3)*.......................................15
**new1Δ-pNew1(HIS3) vs new1Δ-pEV(HIS3)**
**Supplementary Figure 33 |** General overview *new1Δ*-pNew1(*HIS3*) vs *new1Δ*-pEV(*HIS3*) ................................16
**Supplementary Figure 34 |** Gene expression *new1Δ*-pNew1(*HIS3*) vs *new1Δ*-pEV(*HIS3*) ...................................18
**Supplementary Figure 35|** Transcript expression *new1Δ*-pNew1(*HIS3*) vs *new1Δ*-pEV(*HIS3*) ............................20
**new1Δ-pNew1(HIS3) vs WT-pEV(HIS3)**
**Supplementary Figure 36 |** General overview *new1Δ*-pNew1*(HIS3) vs* WT-pEV*(HIS3)* .......................................21
**Supplementary Figure 37 |** Gene expression *new1Δ*-pNew1*(HIS3) vs* WT-pEV*(HIS3)*........................................23
**Supplementary Figure 38 |** Transcript expression *new1Δ*-pNew1*(HIS3) vs* WT-pEV*(HIS3)*. ................................25
**new1Δ-pNew1(HIS3) vs WT-pNew1(HIS3)**
**Supplementary Figure 39 |** General overview *new1Δ*-pNew1*(HIS3) vs* WT-pNew1*(HIS3)* ..................................26
**Supplementary Figure 40 |** Gene expression *new1Δ*-pNew1*(HIS3) vs* WT-pNew1*(HIS3)*....................................28
**Supplementary Figure 41 |** Transcript expression *new1Δ*-pNew1*(HIS3) vs* WT-pNew1*(HIS3)*............................30

#### *General supplementary figures description*

**Sequencing overview** – Refers to sequencing metrics including total reads generated per barcode/sample, mapped reads, gene counts and transcript counts.

**General overview –** Refers to data collected from NanopoReaTA regaring the general sample/condition overview including: Read length overview (per sample and condition), gene expression variability (per sample and condition), change in gene composition (per sample and condition) and Process time.

**Gene expression** - Refers to data collected from NanopoReaTA regaring differential gene expression analysis including: Principal Component Analysis (PCA), Volcano plots, Sample-to-sample variaiblity and Heatmaps.

**Transcript expression** - Refers to data collected from NanopoReaTA regaring differential transcript expression analysis including: Principal Component Analysis (PCA), Volcano plot, Sample-to-sample variaiblity and Heatmaps.

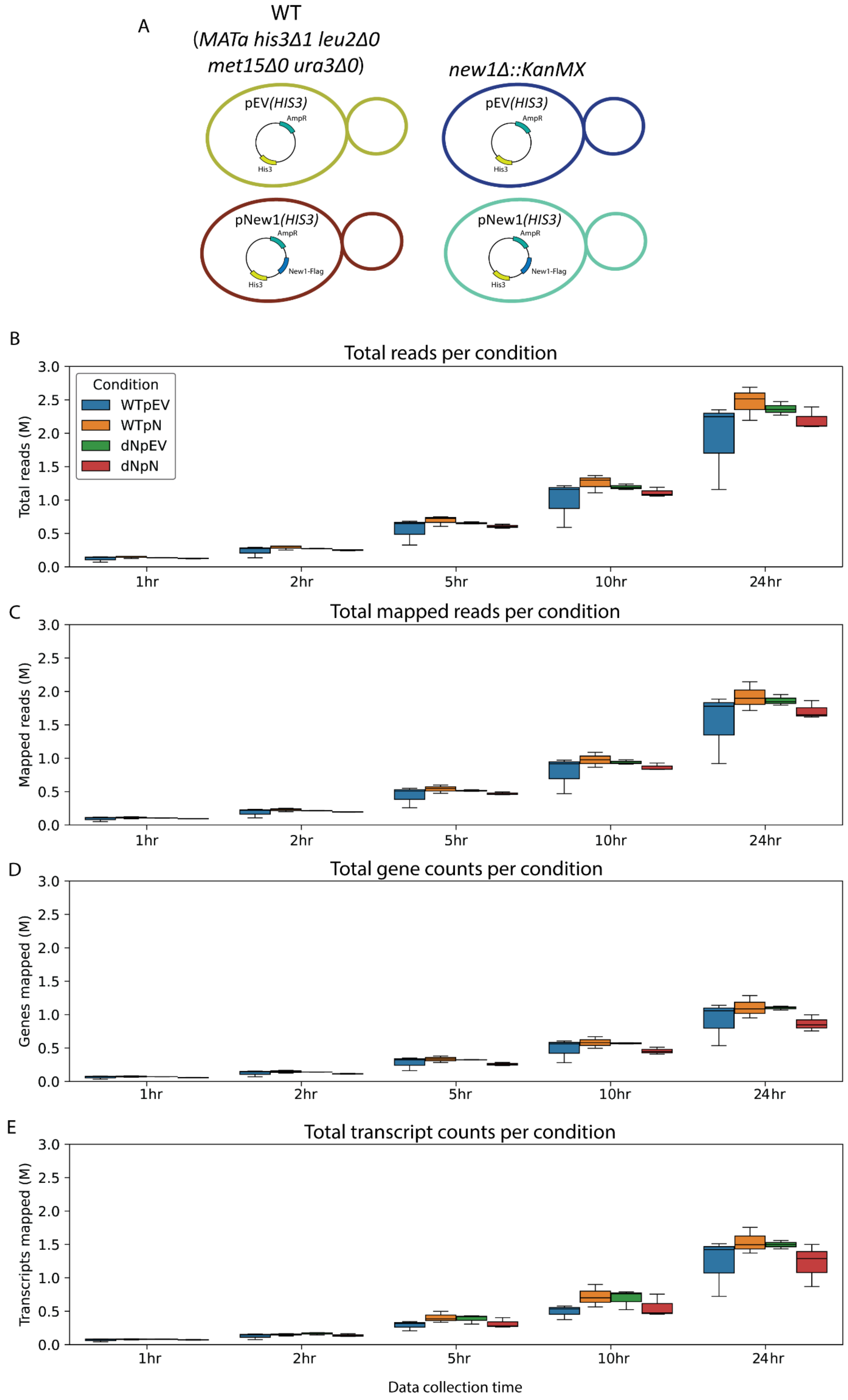
A
WT
(*MATa his3Δ1 leu2Δ0 met15Δ0 ura3Δ0*)
*new1Δ::KanMX*
pEV(*HIS3*)
AmpR
His3
pNew1(*HIS3*)
New1-Flag
B
Total reads per condition
Total reads (M)
Condition
WTpEV
WTpN
dNpEV
dNpN
1hr
2hr
5hr
10hr
24hr
C
Total mapped reads per condition
Mapped reads (M)
D
Total gene counts per condition
Genes mapped (M)
E
Total transcript counts per condition
Transcripts mapped (M)
Data collection time
0.0
0.5
1.0
1.5
2.0
2.5
3.0

**Supplementary Figure 25.** Sequencing Overview Yeast Setup 1. **A.** For Yeast setup 1, WT strain (BY4741*, MATa, his3Δ1, leu2Δ0, met15Δ0, ura3Δ0*) and *new1Δ::KanMX*, where the *NEW1* gene was replaced with the KanMX cassette which contains the Kanamycin resistance gene (*KanR*). These strains were transformed with either an empty vector with the *HIS3* selection marker (pEV*(HIS3)*) or an overexpression vector for C-terminally FLAG-tagged New1 with the *HIS3* selection marker (pNew1*(HIS3)*). **B-E.** The box plots represent total read generated (**B**), total mapped reads (**C**), total gene counts (**D**) and total transcript count (**E**) per condition, per time point (3 biological replicates per condition). Abbreviations: WTpEV - WTpEV*(HIS3)*, WTpN - WTpNew1*(HIS3)*, dNpEV - *new1Δ*pEV*(HIS3)*, dNpN - *new1Δ*pNew1*(HIS3).*

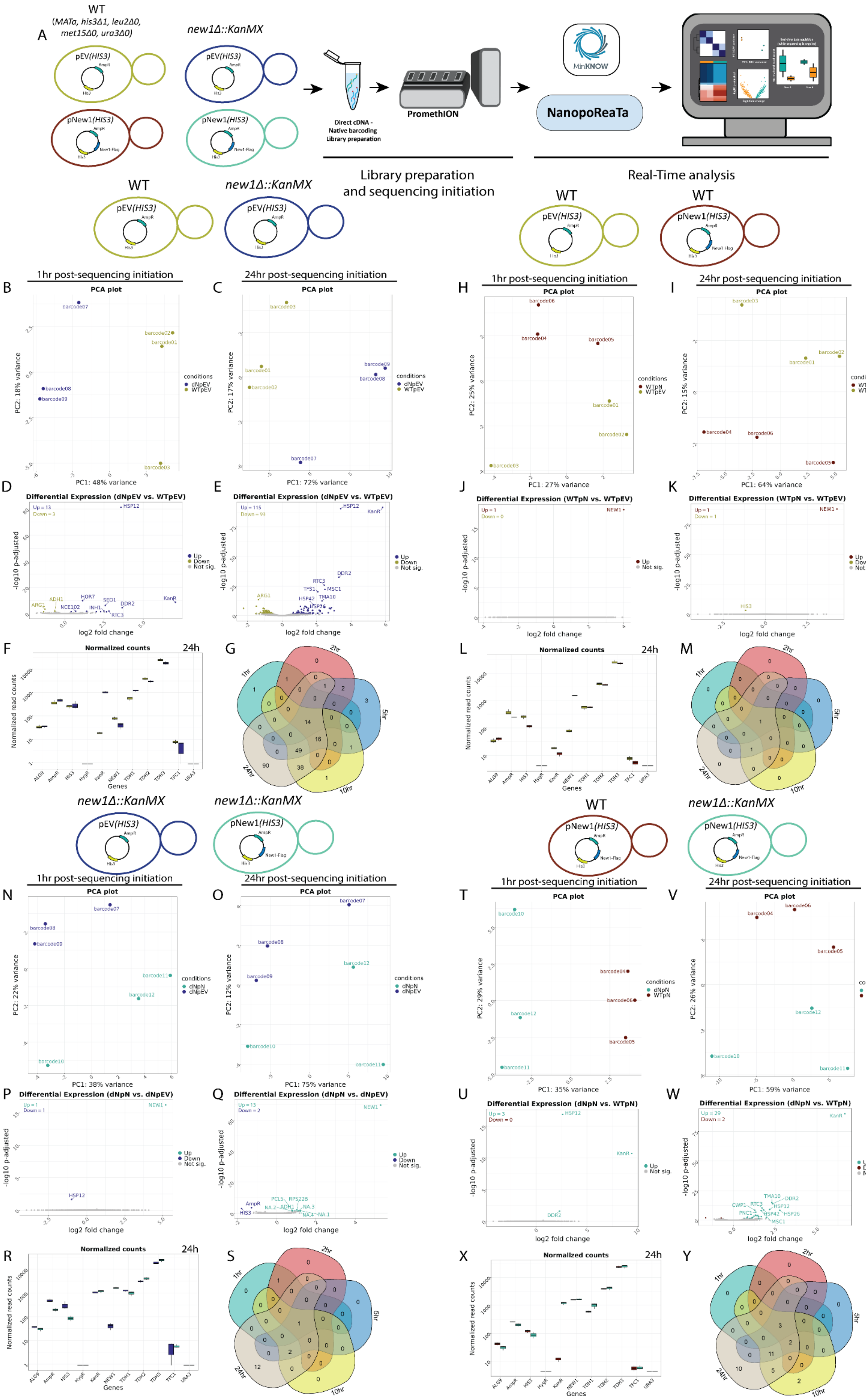
A
WT
(MATa, his3Δ1, leu2Δ0, met15Δ0, ura3Δ0)
new1Δ::KanMX
pEV(HIS3)
pNew1(HIS3)
Direct cDNA - Native barcoding Library preparation
PromethION
MinKNOW
NanopoReaTa
Library preparation and sequencing initiation
Real-Time analysis
1hr post-sequencing initiation
24hr post-sequencing initiation
PCA plot
PC1: 48% variance
PC2: 18% variance
PC1: 72% variance
PC2: 17% variance
PC1: 27% variance
PC2: 25% variance
PC1: 64% variance
PC2: 15% variance
conditions
dNpEV
WTpEV
WTpN
Differential Expression (dNpEV vs. WTpEV)
Differential Expression (WTpN vs. WTpEV)
Differential Expression (dNpN vs. dNpEV)
Differential Expression (dNpN vs. WTpN)
-log10 p-adjusted
log2 fold change
Up
Down
Not sig.
Normalized counts
24h
Normalized read counts
Genes
1hr
2hr
5hr
10hr
24hr
PC1: 38% variance
PC2: 22% variance
PC1: 75% variance
PC2: 12% variance
PC1: 35% variance
PC2: 29% variance
PC1: 59% variance
PC2: 26% variance
dNpN

**Supplementary Figure 26. Real-time transcriptomic analysis in yeast setup 1 samples using NanopoReaTA. A.** Experimental strategy of yeast setup 1. RNA was isolated from selected yeast strains and dscDNA library was prepared which included samples barcoding and adapter ligation. Samples were loaded and sequenced using a PromethION R10 flow cell. NanopoReaTA was activated shortly after sequencing initiation and data was collected 1hr, 2hr, 5hr, 10hr, and 24hr post-sequencing initiation. **B-G.** Differential gene expression in *new1Δ*-pEV(*HIS3*) versus WT-pEV(*HIS3*). Selected data plots showing PCA (B-C) and volcano plots (D-E) 1hr and 24hr post sequencing initiation. **F.** Normalized gene count for selected genes 24hr post sequencing initiation. Normalized gene counts are visualized for selected genes per condition using boxplots. The median-of-ratios normalization method from DESeq2 was used for normalization. **G.** Five-way Venn diagram showing the differentially expressed gene overlaps between the distinct collected time points. **H-M.** Differential gene expression in WT-pNew1(*HIS3*) versus WT-pEV(*HIS3*). Similar analyses to B-G were conducted for PCA (**H-I**) and volcano plots (**J-K**) as well as normalized gene counts (**L**) and Venn diagram (**M**). **N-S**. Differential gene expression in *new1Δ*-pNew1(*HIS3*) versus *new1Δ*-pEV(*HIS3*). Similar analyses to B-G were conducted for PCA (**N-O**) and volcano plots (**P-Q**) as well as normalized gene counts (**R**) and Venn diagram (**S**). **T-Y.** Differential gene expression in *new1Δ*-pNew1(*HIS3*) versus WT-pNew1(*HIS3*). Similar analyses to B-G were conducted for PCA (**T-V**) and volcano plots (**U-W**) as well as normalized gene counts (**X**) and Venn diagram (**Y**).

#### General overview
#### *new1Δ*-pEV(*HIS3*) *vs* WT-pEV(*HIS3*)

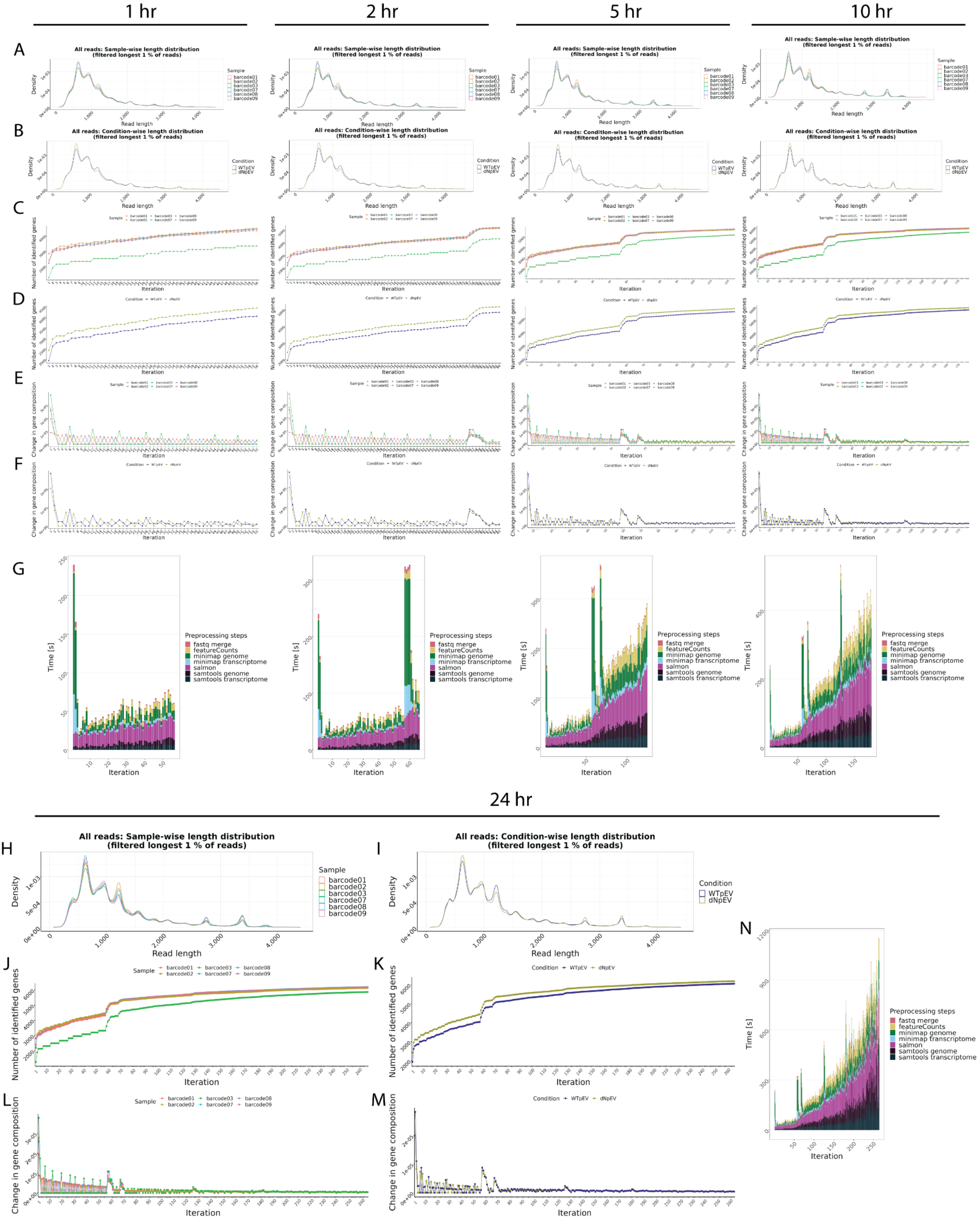

**Supplementary Figure 27. General overview in *new1Δ*-pEV*(HIS3) vs* WT-pEV*(HIS3)*. A-B**. Read length overview. The distribution of read lengths derived from generated fastq files is plotted per sample (A) and per condition (B). All reads of length over the 99 % quantile of all lengths are removed from these visualizations. **C-D**. Gene expression variability. The number of identified genes (> 0 reads counted) is plotted after each iteration per sample (C) and per condition (D). When the detection of additional genes finishes, the lines reach a plateau. **E-F**. Change in gene composition. The change in gene composition (CGC) plot shows the sum of absolute differences of relative gene (%gi) counts per individual gene (g) in iteration (i) to the relative gene counts in iteration (i-1) for each respective sample (E) and condition (F). CGC = sum(|(%g(i-1))-(%g(i))|). **G.** Process time. The bar plots show the time in seconds all preprocessing steps needed per iteration. One iteration process at maximum 30 files from all samples. This plot updates automatically as a new process finish. **H-N**. Final general overview. Read length (H-I), Gene expression variability (J-K), Change in gene composition (L-M), and processing time (N) for the 24hr time point. All plots are organized according to their respective time points of collection.

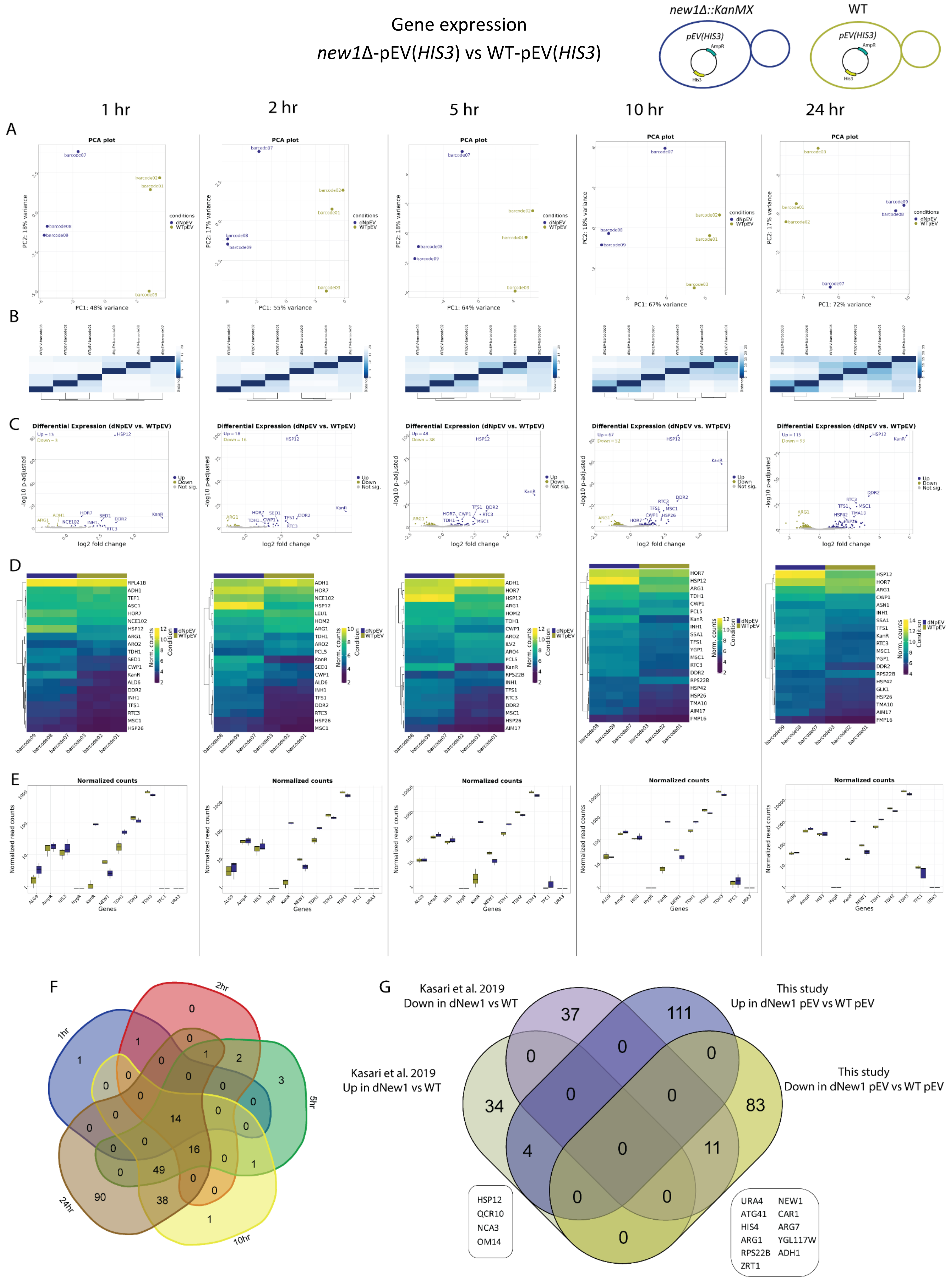
Gene expression
new1Δ-pEV(HIS3) vs WT-pEV(HIS3)
new1Δ::KanMX
WT
pEV(HIS3)
AmpR
His3
1 hr
2 hr
5 hr
10 hr
24 hr
A
PCA plot
PC1: 48% variance
PC1: 55% variance
PC1: 64% variance
PC1: 67% variance
PC1: 72% variance
conditions
dNpEV
WTpEV
B
C
Differential Expression (dNpEV vs. WTpEV)
-log10 p-adjusted
log2 fold change
Up
Down
Not sig.
D
Norm. counts
Condition
E
Normalized counts
Normalized read counts
Genes
F
1hr
2hr
5hr
10hr
24hr
G
Kasari et al. 2019
Down in dNew1 vs WT
This study
Up in dNew1 pEV vs WT pEV
Kasari et al. 2019
Up in dNew1 vs WT
This study
Down in dNew1 pEV vs WT pEV
37
111
0
0
0
34
0
0
83
4
0
11
0
0
0
HSP12
QCR10
NCA3
OM14
URA4
ATG41
HIS4
ARG1
RPS22B
ZRT1
NEW1
CAR1
ARG7
YGL117W
ADH1

**Supplementary Figure 28. Gene expression in *new1Δ*-pEV*(HIS3)* *vs* WT-pEV*(HIS3)*. A.** Principal Component Analysis (PCA) of the top 500 genes with the highest variance across all samples, the two Principal Components. Each dot corresponds to one sample and is colored by the respective condition. **B.** Sample2sample distance plot. The euclidean distance between the gene expression patterns of all samples to each other is plotted using heatmap. Normalized gene counts were used for distance computation. **C.** Volcano plots. The differential gene expression analysis results from DESeq2 are shown by plotting the log2FoldChange between the conditions of interest against the –log10 adjusted p-value per gene observed from the Wald-Test integrated in DESeq2 [Love, Huber, and Anders et al. 2014]. The top 10 significant genes are labeled by their symbol. **D.** Heatmaps. The expression of the top 20 differentially expressed genes is plotted using Heatmap by coloring the number of reads per gene. The legend on the right side describes which colors correspond to highly and lowly expressed genes. **E**. Normalized gene count for selected genes. Normalized gene counts from FeatureCounts [Liao et al. 2014] are visualized for selected genes per condition using boxplots. The median-of-ratios normalization method from DESeq2 [Love et al. 2014] was used for normalization. **F**. 5-way Venn diagram showing the differentially expressed gene overlaps between the distinct collected time points. **G**. Overlap of Up- and Down-regulated DEGs between this study and Kasari et al. 2019.

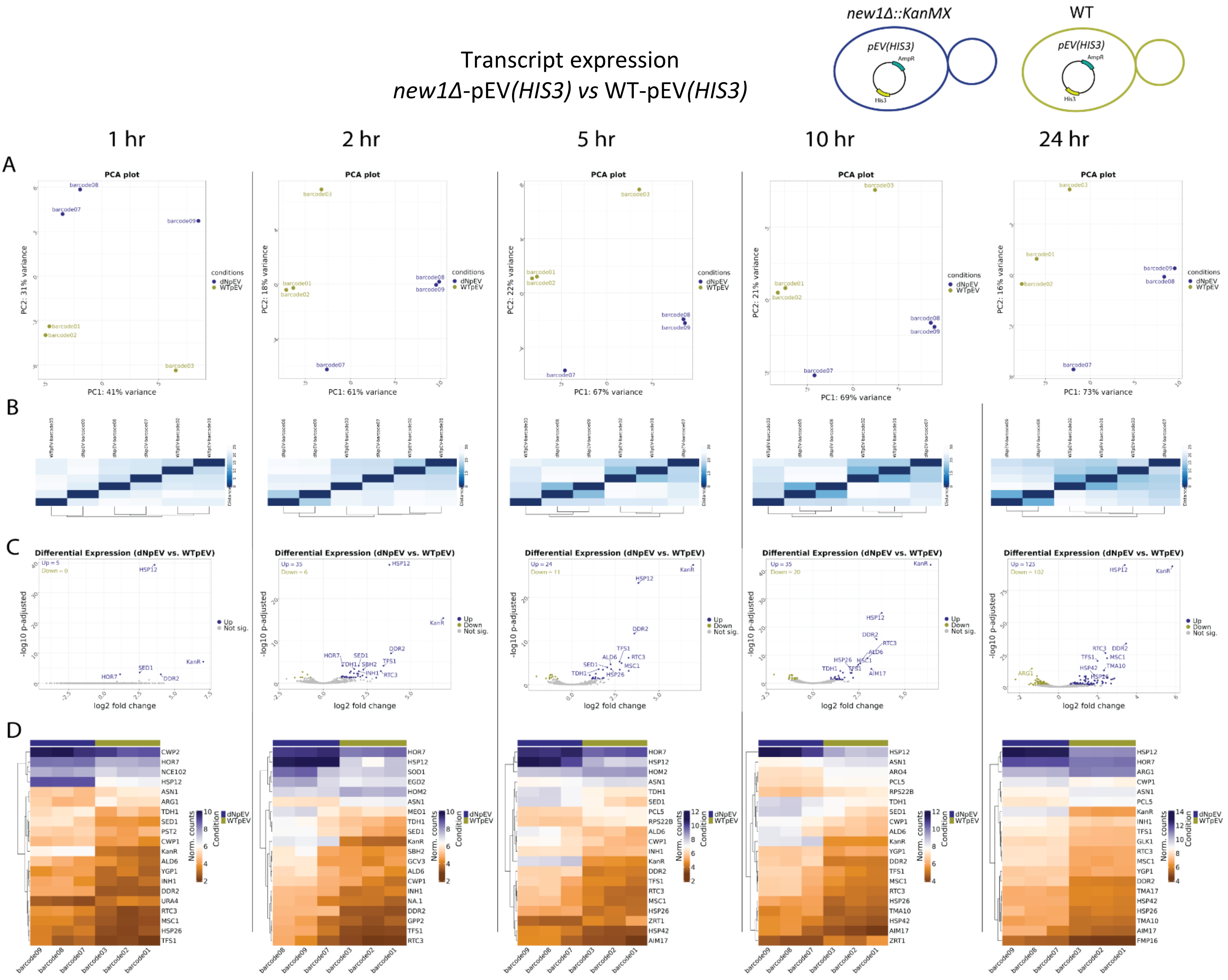


**Supplementary Figure 29. Transcript expression *new1Δ-pEV(HIS3) vs WT-pEV(HIS3)*. A.** Principal Component Analysis (PCA) of the top 500 transcripts with the highest variance across all samples, the two Principal Components. Each dot corresponds to one sample and is colored by the respective condition. **B.** Sample2sample plot. The euclidean distance between the transcript expression patterns of all samples to each other is plotted using heatmap. Normalized gene counts were used for distance computation. **C.** Volcano plots. The differential transcript expression analysis results from DESeq2 are shown by plotting the log2FoldChange between the conditions of interest against the –log10 adjusted p-value per gene observed from the Wald-Test integrated in DESeq2. The top 10 significant genes are labeled by their symbol. **D.** Heatmaps. The expression of the top 20 differentially expressed transcripts is plotted using Heatmap by coloring the number of reads per transcript. The legend on the right side describes which colors correspond to highly and lowly expressed transcripts.

General overview

*new1Δ*-pEV(*HIS3*) *vs* WT-pEV(*HIS3*)

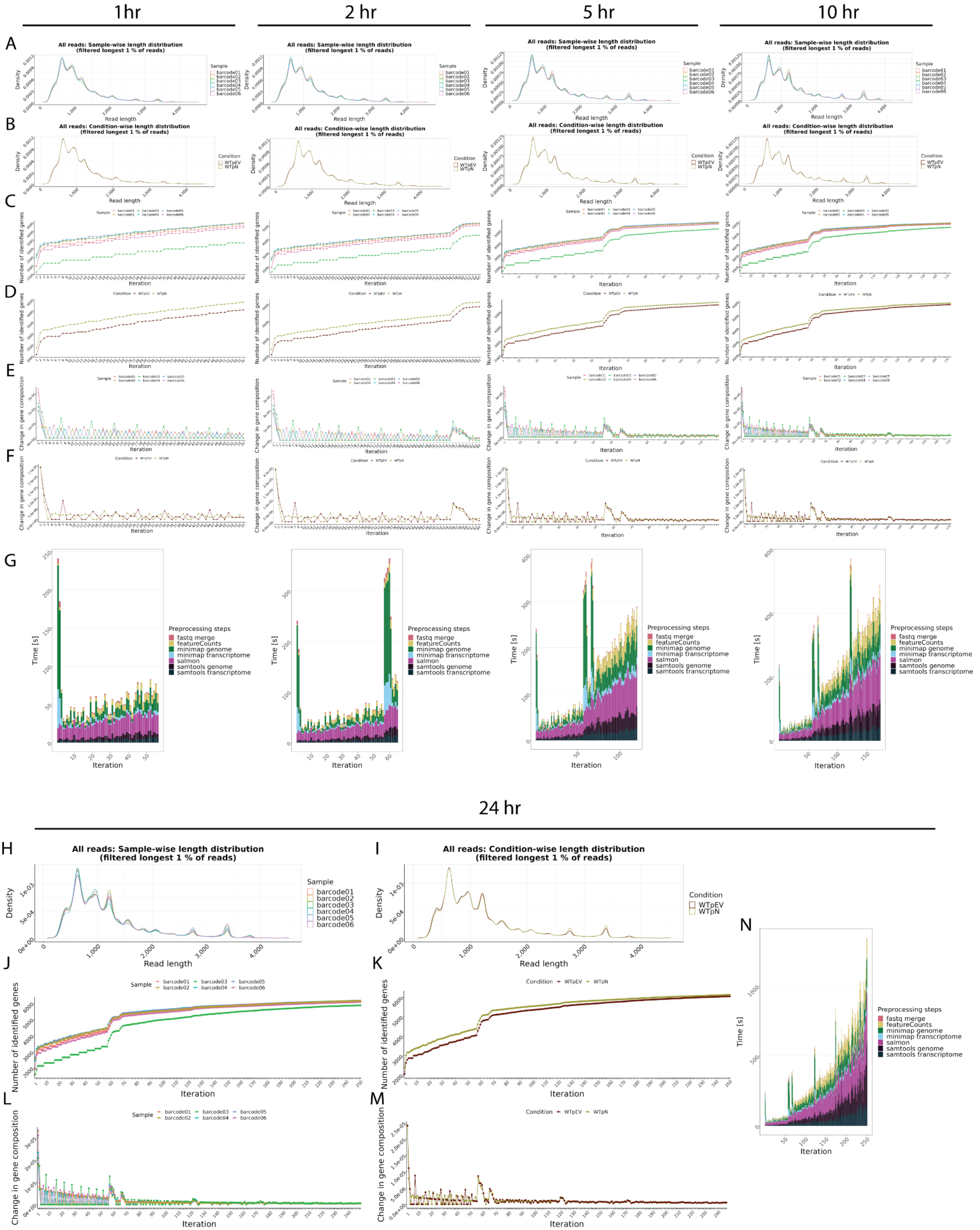

**Supplementary Figure 30. General overview in WT-pNew1*(HIS3)* *vs* WT-pEV*(HIS3)*. A-B**. Read length overview. The distribution of read lengths derived from generated fastq files is plotted per sample (A) and per condition (B). All reads of length over the 99 % quantile of all lengths are removed from these visualizations. **C-D**. Gene expression variability. The number of identified genes (> 0 reads counted) is plotted after each iteration per sample (C) and per condition (D). When the detection of additional genes finishes, the lines reach a plateau. **E-F**. Change in gene composition. The change in gene composition (CGC) plot shows the sum of absolute differences of relative gene (%gi) counts per individual gene (g) in iteration (i) to the relative gene counts in iteration (i-1) for each respective sample (E) and condition (F). CGC = sum(|(%g(i-1))-(%g(i))|). **G.** Process time. The bar plots show the time in seconds all preprocessing steps needed per iteration. One iteration process at maximum 30 files from all samples. This plot updates automatically as a new process finish. **H-N**. Final general overview. Read length (H-I), Gene expression variability (J-K), Change in gene composition (L-M), and processing time (N) for the 24hr time point. All plots are organized according to their respective time points of collection.

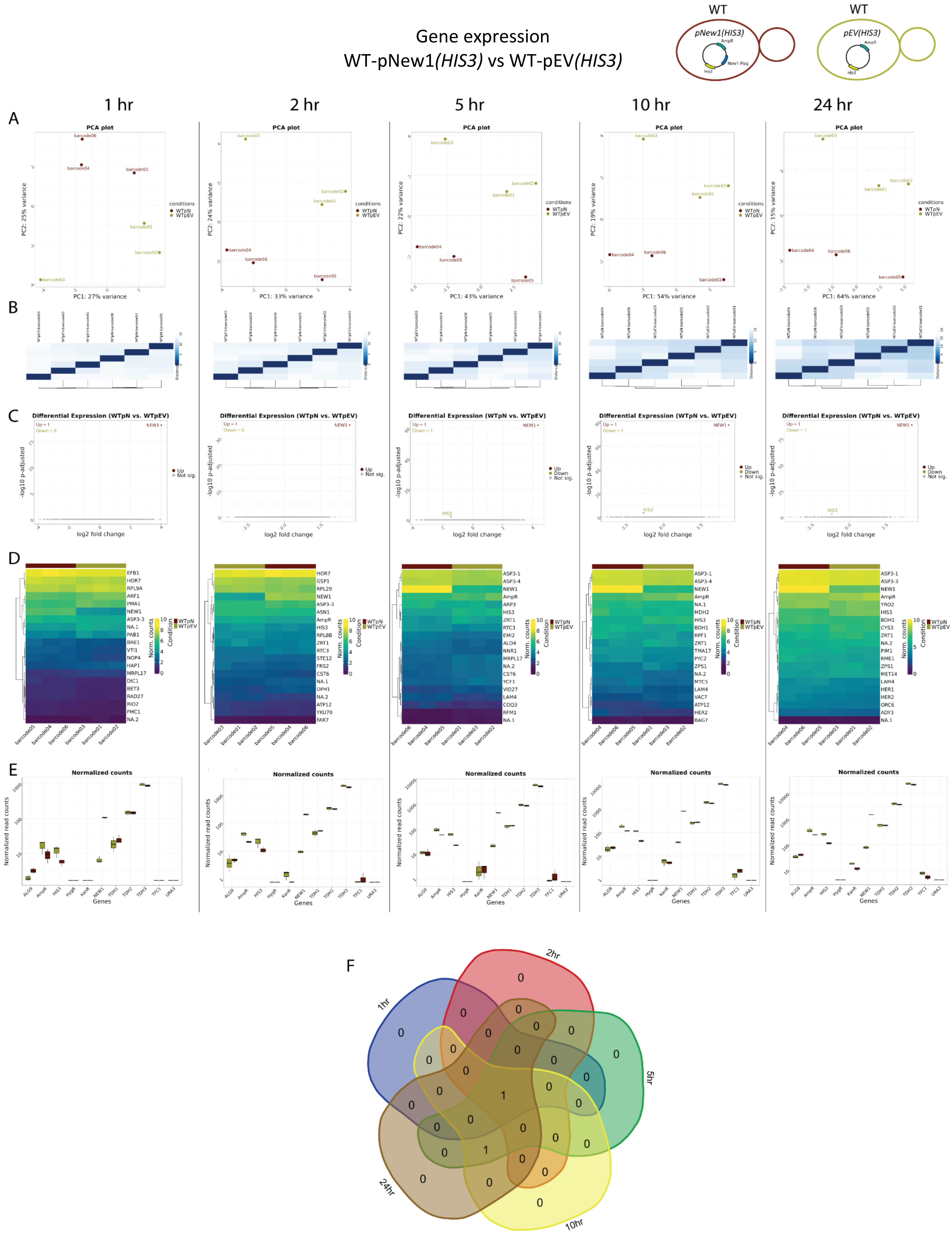
Gene expression
WT-pNew1(HIS3) vs WT-pEV(HIS3)
WT
pNew1(HIS3)
WT
pEV(HIS3)
1 hr
2 hr
5 hr
10 hr
24 hr
A
PCA plot
B
C
Differential Expression (WTpN vs. WTpEV)
D
E
Normalized counts
F
1hr
2hr
5hr
10hr
24hr

**Supplementary Figure 31. Gene expression in WT-pNew1*(HIS3) vs* WT-pEV*(HIS3)*. A.** Principal Component Analysis (PCA) of the top 500 genes with the highest variance across all samples, the two Principal Components. Each dot corresponds to one sample and is colored by the respective condition. **B.** Sample2sample distance plot. The euclidean distance between the gene expression patterns of all samples to each other is plotted using heatmap. Normalized gene counts were used for distance computation. **C.** Volcano plots. The differential gene expression analysis results from DESeq2 are shown by plotting the log2FoldChange between the conditions of interest against the –log10 adjusted p-value per gene observed from the Wald-Test integrated in DESeq2 [Love, Huber, and Anders et al. 2014]. The top 10 significant genes are labeled by their symbol. **D.** Heatmaps. The expression of the top 20 differentially expressed genes is plotted using Heatmap by coloring the number of reads per gene. The legend on the right side describes which colors correspond to highly and lowly expressed genes. **E**. Normalized gene count for selected genes. Normalized gene counts from FeatureCounts [Liao et al. 2014] are visualized for selected genes per condition using boxplots. The median-of-ratios normalization method from DESeq2 [Love et al. 2014] was used for normalization. **F**. 5-way Venn diagram showing the differentially expressed gene overlaps between the distinct collected time points.

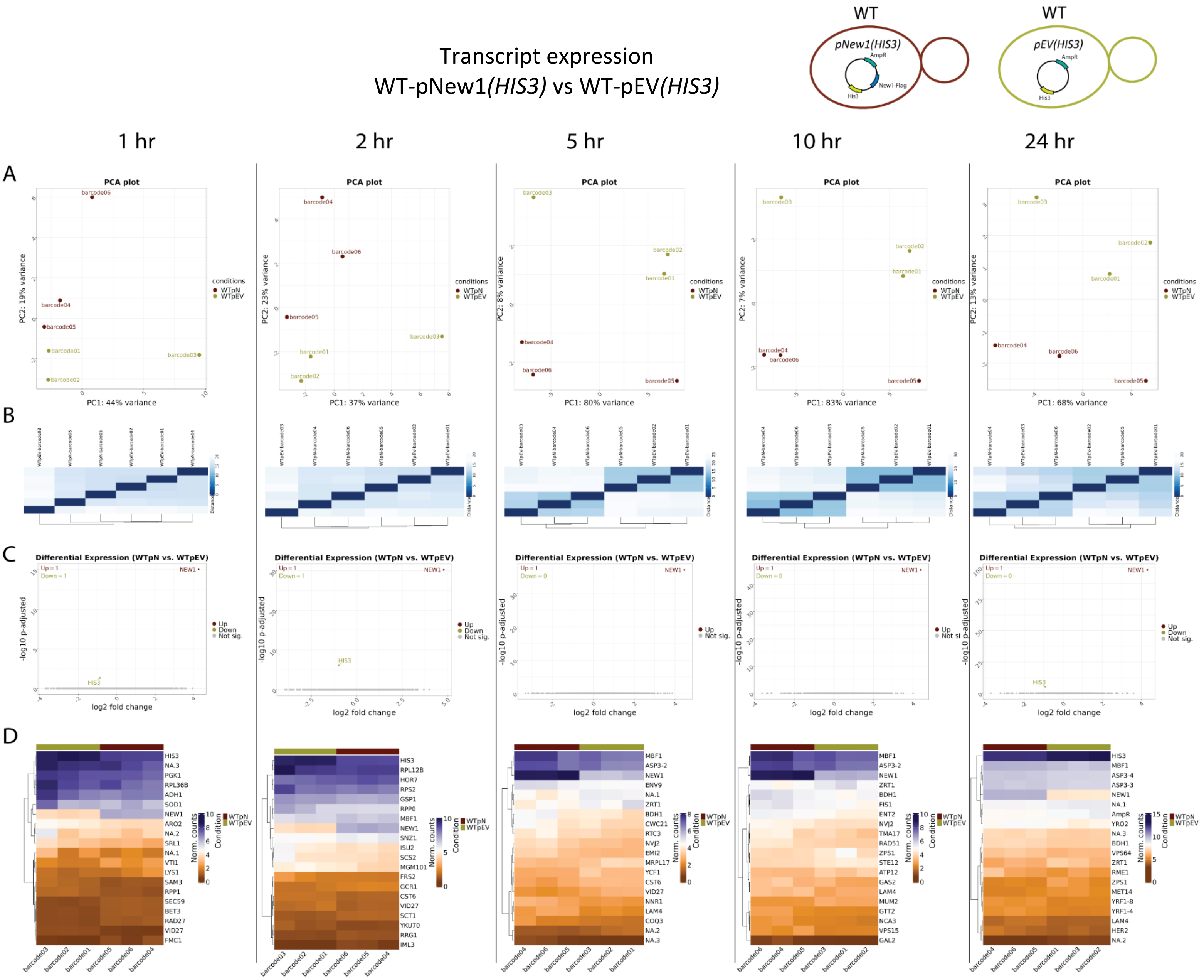


**Supplementary Figure 32. Gene expression in WT-pNew1*(HIS3) vs* WT-pEV*(HIS3)*. A.** Principal Component Analysis (PCA) of the top 500 transcripts with the highest variance across all samples, the two Principal Components. Each dot corresponds to one sample and is colored by the respective condition. **B.** Sample2sample plot. The euclidean distance between the transcript expression patterns of all samples to each other is plotted using heatmap. Normalized gene counts were used for distance computation. **C.** Volcano plots. The differential transcript expression analysis results from DESeq2 are shown by plotting the log2FoldChange between the conditions of interest against the –log10 adjusted p-value per gene observed from the Wald-Test integrated in DESeq2. The top 10 significant genes are labeled by their symbol. **D.** Heatmaps. The expression of the top 20 differentially expressed transcripts is plotted using Heatmap by coloring the number of reads per transcript. The legend on the right side describes which colors correspond to highly and lowly expressed transcripts.

General overview

WT-pNew1*(HIS3)* vs WT-pEV*(HIS3)*

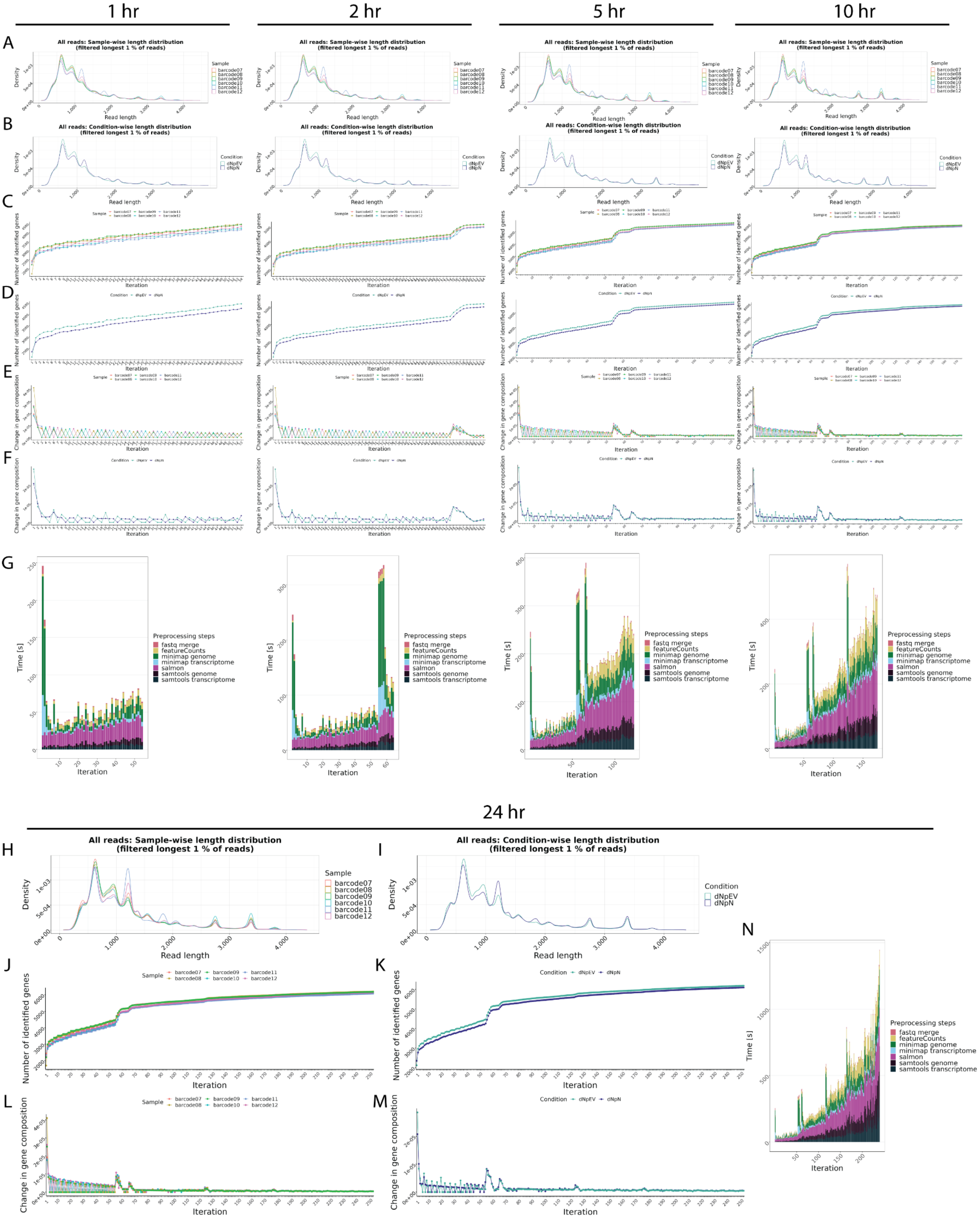

**Supplementary Figure 33. General overview in *new1Δ*-pNew1*(HIS3) vs new1Δ*-pEV*(HIS3)*. A-B**. Read length overview. The distribution of read lengths derived from generated fastq files is plotted per sample (A) and per condition (B). All reads of length over the 99 % quantile of all lengths are removed from these visualizations. **C-D**. Gene expression variability. The number of identified genes (> 0 reads counted) is plotted after each iteration per sample (C) and per condition (D). When the detection of additional genes finishes, the lines reach a plateau. **E-F**. Change in gene composition. The change in gene composition (CGC) plot shows the sum of absolute differences of relative gene (%gi) counts per individual gene (g) in iteration (i) to the relative gene counts in iteration (i-1) for each respective sample (E) and condition (F). CGC = sum(|(%g(i-1))-(%g(i))|). **G.** Process time. The bar plots show the time in seconds all preprocessing steps needed per iteration. One iteration process at maximum 30 files from all samples. This plot updates automatically as a new process finish. **H-N**. Final general overview. Read length (H-I), Gene expression variability (J-K), Change in gene composition (L-M), and processing time (N) for the 24hr time point. All plots are organized according to their respective time points of collection.

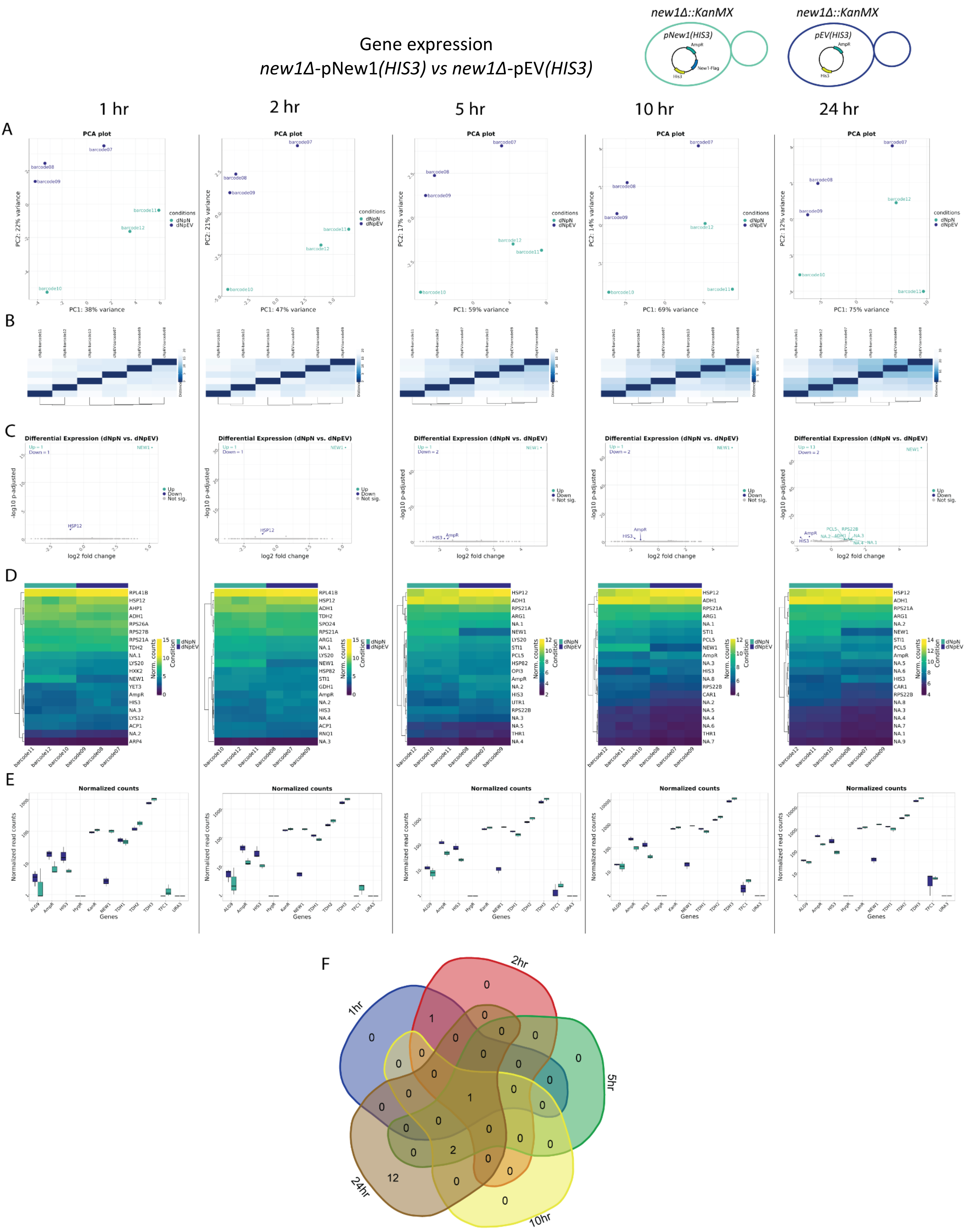
Gene expression
new1Δ-pNew1(HIS3) vs new1Δ-pEV(HIS3)
new1Δ::KanMX
pNew1(HIS3)
AmpR
New1-Flag
His3
new1Δ::KanMX
pEV(HIS3)
AmpR
His3
1 hr
2 hr
5 hr
10 hr
24 hr
A
PCA plot
PC1: 38% variance
PC2: 22% variance
PC1: 47% variance
PC2: 21% variance
PC1: 59% variance
PC2: 17% variance
PC1: 69% variance
PC2: 14% variance
PC1: 75% variance
PC2: 12% variance
conditions
dNpN
dNpEV
B
C
Differential Expression (dNpN vs. dNpEV)
log2 fold change
-log10 p-adjusted
Up
Down
Not sig.
D
Norm. counts
Condition
E
Normalized counts
Normalized read counts
Genes
F
1hr
2hr
5hr
10hr
24hr

**Supplementary Figure 34. Gene expression in *new1Δ*-pNew1*(HIS3)* *vs* *new1Δ*-pEV*(HIS3)*. A.** Principal Component Analysis (PCA) of the top 500 genes with the highest variance across all samples, the two Principal Components. Each dot corresponds to one sample and is colored by the respective condition. **B.** Sample2sample distance plot. The euclidean distance between the gene expression patterns of all samples to each other is plotted using heatmap. Normalized gene counts were used for distance computation. **C.** Volcano plots. The differential gene expression analysis results from DESeq2 are shown by plotting the log2FoldChange between the conditions of interest against the –log10 adjusted p-value per gene observed from the Wald-Test integrated in DESeq2 [Love, Huber, and Anders et al. 2014]. The top 10 significant genes are labeled by their symbol. **D.** Heatmaps. The expression of the top 20 differentially expressed genes is plotted using Heatmap by coloring the number of reads per gene. The legend on the right side describes which colors correspond to highly and lowly expressed genes. **E**. Normalized gene count for selected genes. Normalized gene counts from FeatureCounts [Liao et al. 2014] are visualized for selected genes per condition using boxplots. The median-of-ratios normalization method from DESeq2 [Love et al. 2014] was used for normalization. **F**. 5-way Venn diagram showing the differentially expressed gene overlaps between the distinct collected time points.

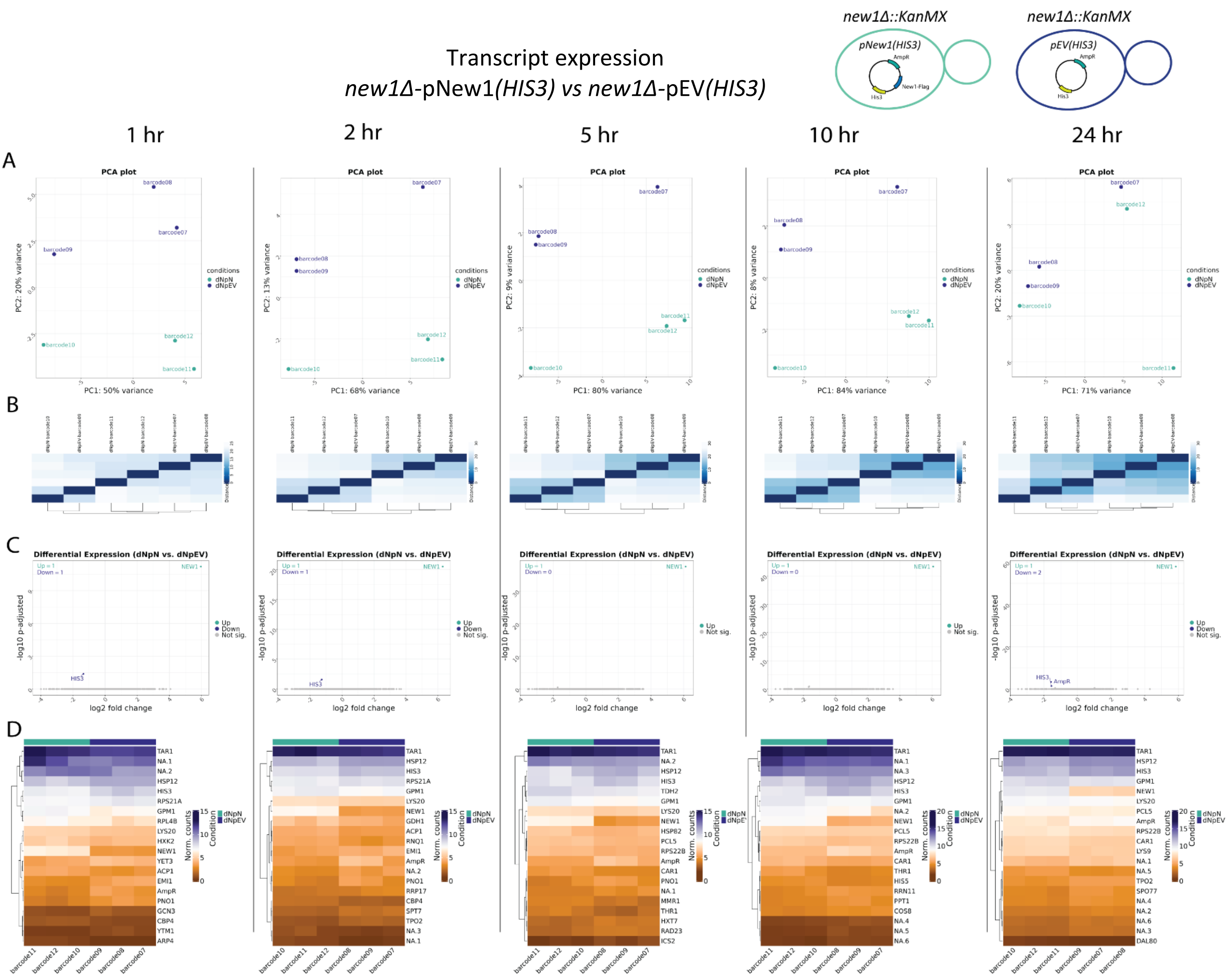


**Supplementary Figure 35. Transcript expression in *new1Δ*-pNew1*(HIS3)* vs *new1Δ*-pEV*(HIS3)*. A.** Principal Component Analysis (PCA) of the top 500 transcripts with the highest variance across all samples, the two Principal Components. Each dot corresponds to one sample and is colored by the respective condition. **B.** Sample2sample plot. The euclidean distance between the transcript expression patterns of all samples to each other is plotted using heatmap. Normalized gene counts were used for distance computation. **C.** Volcano plots. The differential transcript expression analysis results from DESeq2 are shown by plotting the log2FoldChange between the conditions of interest against the –log10 adjusted p-value per gene observed from the Wald-Test integrated in DESeq2. The top 10 significant genes are labeled by their symbol. **D.** Heatmaps. The expression of the top 20 differentially expressed transcripts is plotted using Heatmap by coloring the number of reads per transcript. The legend on the right side describes which colors correspond to highly and lowly expressed transcripts.

General overview
*new1Δ*-pNew1(*HIS3*) *vs new1Δ*-pEV(*HIS3*)

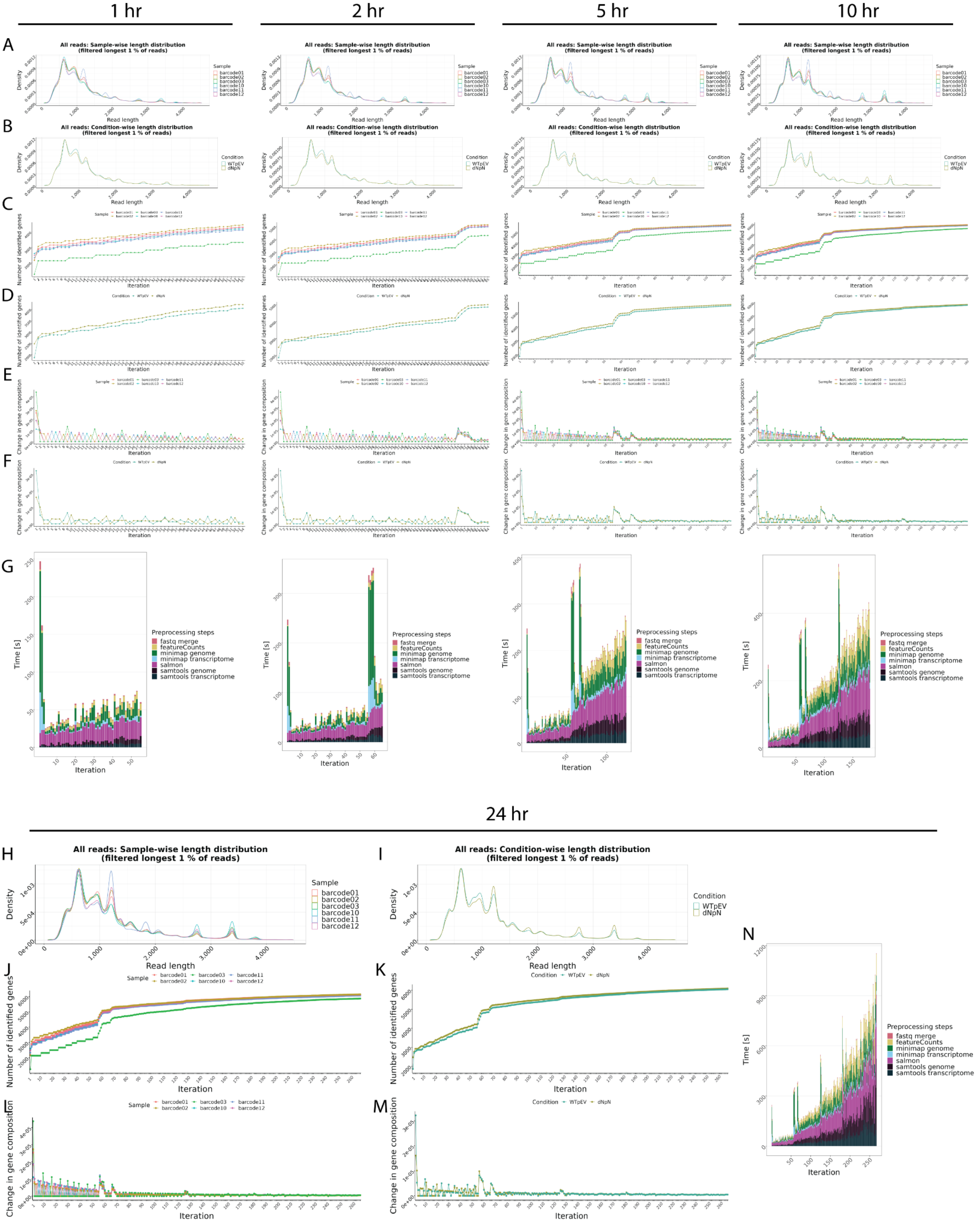

**Supplementary Figure 36. General overview in *new1Δ*-pNew1*(HIS3) vs* WT-pEV*(HIS3)*. A-B**. Read length overview. The distribution of read lengths derived from generated fastq files is plotted per sample (A) and per condition (B). All reads of length over the 99 % quantile of all lengths are removed from these visualizations. **C-D**. Gene expression variability. The number of identified genes (> 0 reads counted) is plotted after each iteration per sample (C) and per condition (D). When the detection of additional genes finishes, the lines reach a plateau. **E-F**. Change in gene composition. The change in gene composition (CGC) plot shows the sum of absolute differences of relative gene (%gi) counts per individual gene (g) in iteration (i) to the relative gene counts in iteration (i-1) for each respective sample (E) and condition (F). CGC = sum(|(%g(i-1))-(%g(i))|). **G.** Process time. The bar plots show the time in seconds all preprocessing steps needed per iteration. One iteration process at maximum 30 files from all samples. This plot updates automatically as a new process finish. **H-N**. Final general overview. Read length (H-I), Gene expression variability (J-K), Change in gene composition (L-M), and processing time (N) for the 24hr time point. All plots are organized according to their respective time points of collection.

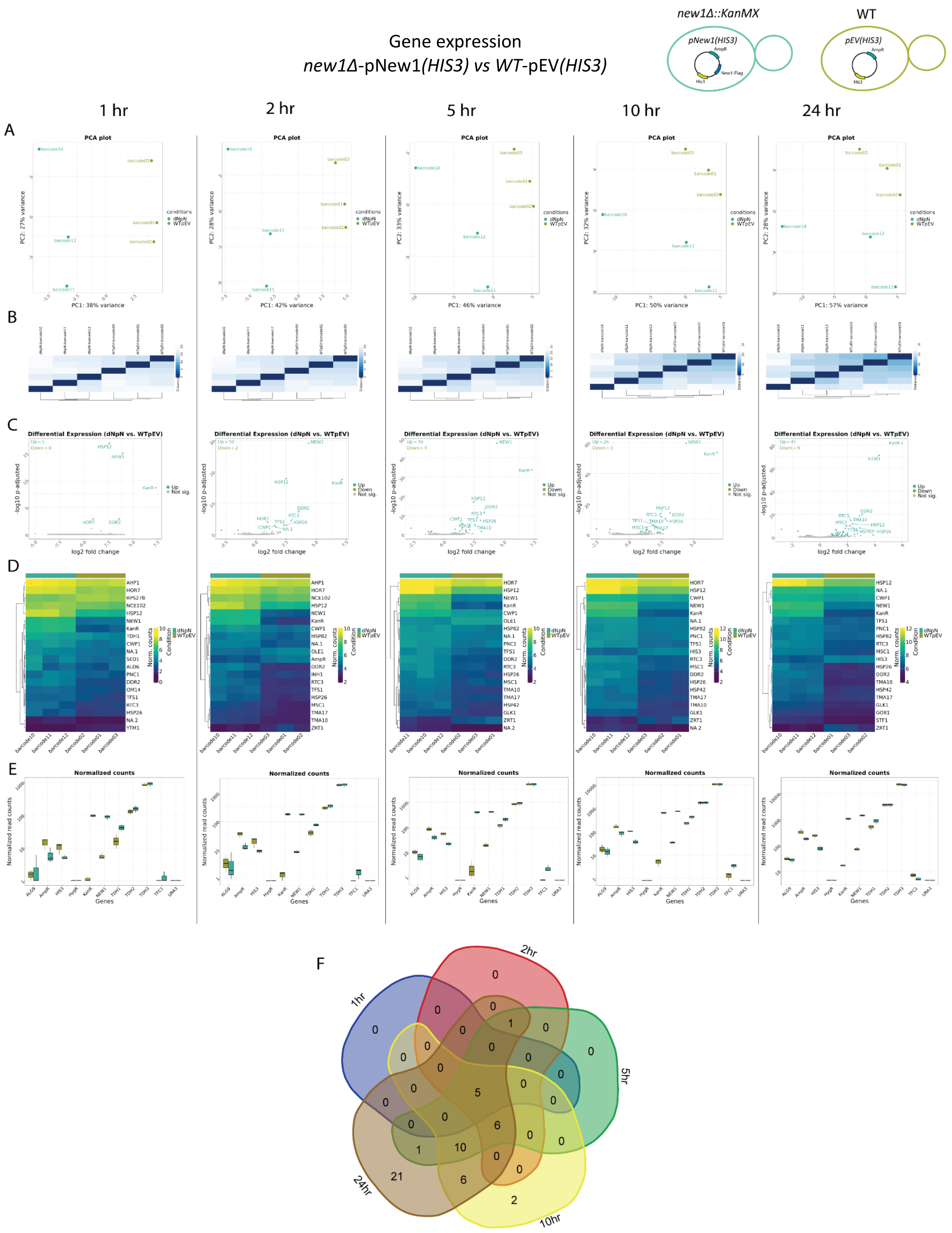
Gene expression
new1Δ-pNew1(HIS3) vs WT-pEV(HIS3)
new1Δ::KanMX
WT
pNew1(HIS3)
pEV(HIS3)
AmpR
New1-Flag
His3
1 hr
2 hr
5 hr
10 hr
24 hr
A
PCA plot
PC1: 38% variance
PC1: 42% variance
PC1: 46% variance
PC1: 50% variance
PC1: 57% variance
PC2: 27% variance
PC2: 28% variance
PC2: 33% variance
PC2: 32% variance
conditions
dNpN
WTpEV
B
C
Differential Expression (dNpN vs. WTpEV)
log2 fold change
-log10 p-adjusted
Up
Down
Not sig.
D
Norm. counts
Condition
E
Normalized counts
Normalized read counts
Genes
F
1hr
2hr
5hr
10hr
24hr

**Supplementary Figure 37. Gene expression in *new1Δ*-pNew1*(HIS3) vs* WT-pEV*(HIS3)*. A.** Principal Component Analysis (PCA) of the top 500 genes with the highest variance across all samples, the two Principal Components. Each dot corresponds to one sample and is colored by the respective condition. **B.** Sample2sample distance plot. The euclidean distance between the gene expression patterns of all samples to each other is plotted using heatmap. Normalized gene counts were used for distance computation. **C.** Volcano plots. The differential gene expression analysis results from DESeq2 are shown by plotting the log2FoldChange between the conditions of interest against the –log10 adjusted p-value per gene observed from the Wald-Test integrated in DESeq2 [Love, Huber, and Anders et al. 2014]. The top 10 significant genes are labeled by their symbol. **D.** Heatmaps. The expression of the top 20 differentially expressed genes is plotted using Heatmap by coloring the number of reads per gene. The legend on the right side describes which colors correspond to highly and lowly expressed genes. **E**. Normalized gene count for selected genes. Normalized gene counts from FeatureCounts [Liao et al. 2014] are visualized for selected genes per condition using boxplots. The median-of-ratios normalization method from DESeq2 [Love et al. 2014] was used for normalization. **F**. 5-way Venn diagram showing the differentially expressed gene overlaps between the distinct collected time points.

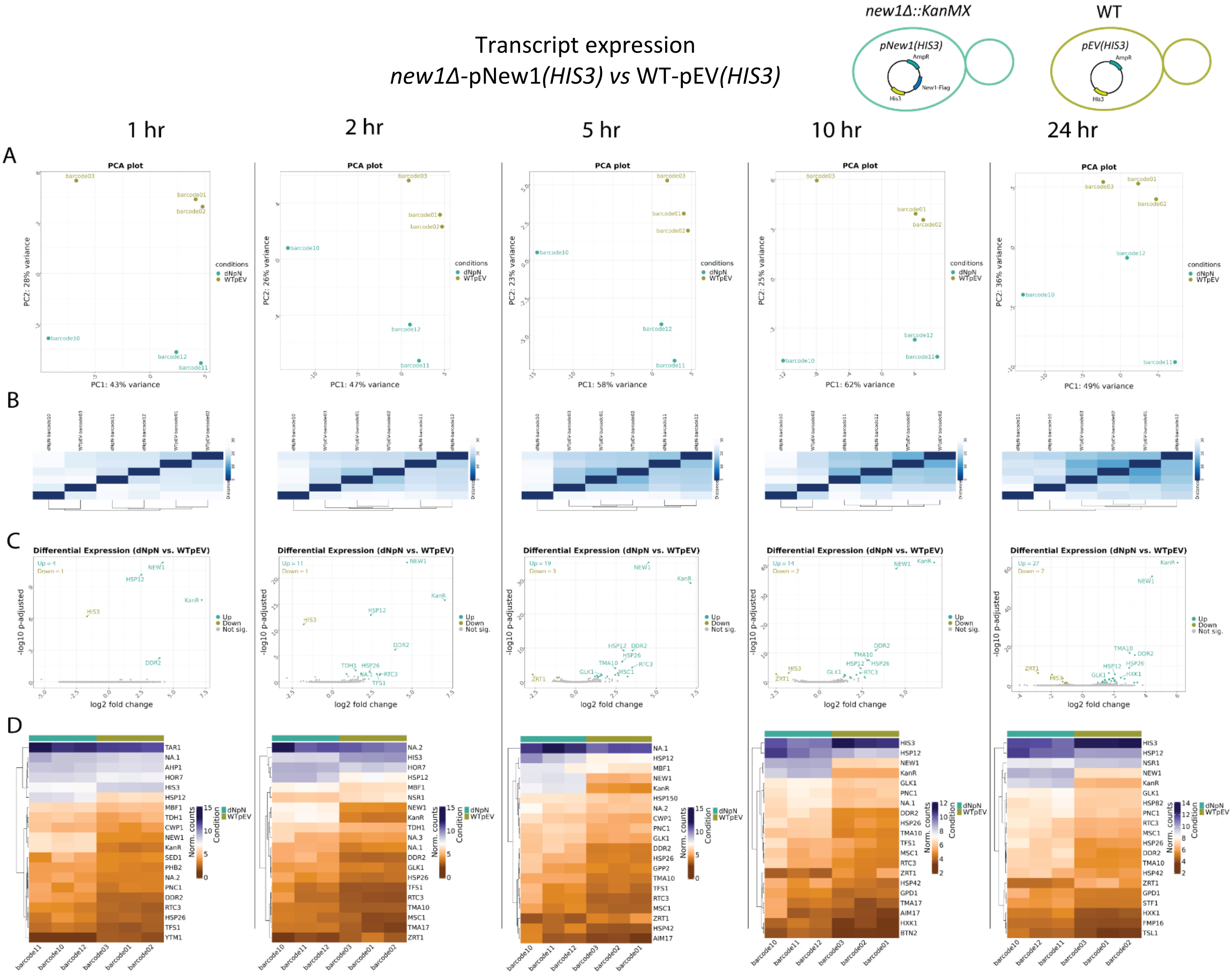


**Supplementary Figure 38. Transcript expression in *new1Δ*-pNew1*(HIS3)* *vs* WT-pEV*(HIS3)*. A.** Principal Component Analysis (PCA) of the top 500 transcripts with the highest variance across all samples, the two Principal Components. Each dot corresponds to one sample and is colored by the respective condition. **B.** Sample2sample plot. The euclidean distance between the transcript expression patterns of all samples to each other is plotted using heatmap. Normalized gene counts were used for distance computation. **C.** Volcano plots. The differential transcript expression analysis results from DESeq2 are shown by plotting the log2FoldChange between the conditions of interest against the –log10 adjusted p-value per gene observed from the Wald-Test integrated in DESeq2. The top 10 significant genes are labeled by their symbol. **D.** Heatmaps. The expression of the top 20 differentially expressed transcripts is plotted using Heatmap by coloring the number of reads per transcript. The legend on the right side describes which colors correspond to highly and lowly expressed transcripts.

#### General overview
#### *new1Δ*-pNew1*(HIS3)* *vs* WT-pEV*(HIS3)*

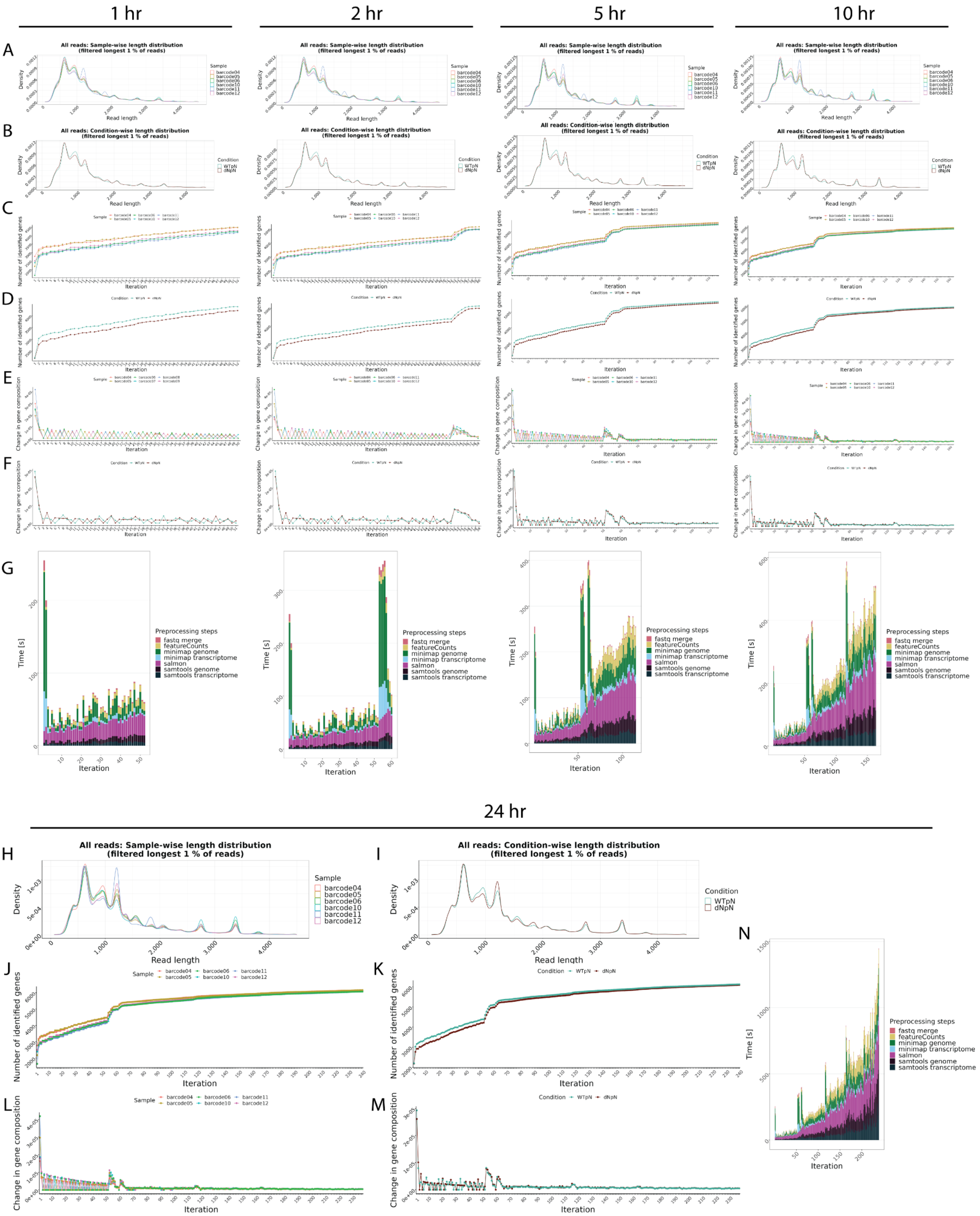

**Supplementary Figure 39. General overview in *new1Δ*-pNew1*(HIS3) vs* WT-pNew1*(HIS3)*. A-B**. Read length overview. The distribution of read lengths derived from generated fastq files is plotted per sample (A) and per condition (B). All reads of length over the 99 % quantile of all lengths are removed from these visualizations. **C-D**. Gene expression variability. The number of identified genes (> 0 reads counted) is plotted after each iteration per sample (C) and per condition (D). When the detection of additional genes finishes, the lines reach a plateau. **E-F**. Change in gene composition. The change in gene composition (CGC) plot shows the sum of absolute differences of relative gene (%gi) counts per individual gene (g) in iteration (i) to the relative gene counts in iteration (i-1) for each respective sample (E) and condition (F). CGC = sum(|(%g(i-1))-(%g(i))|). **G.** Process time. The bar plots show the time in seconds all preprocessing steps needed per iteration. One iteration process at maximum 30 files from all samples. This plot updates automatically as a new process finish. **H-N**. Final general overview. Read length (H-I), Gene expression variability (J-K), Change in gene composition (L-M), and processing time (N) for the 24hr time point. All plots are organized according to their respective time points of collection.

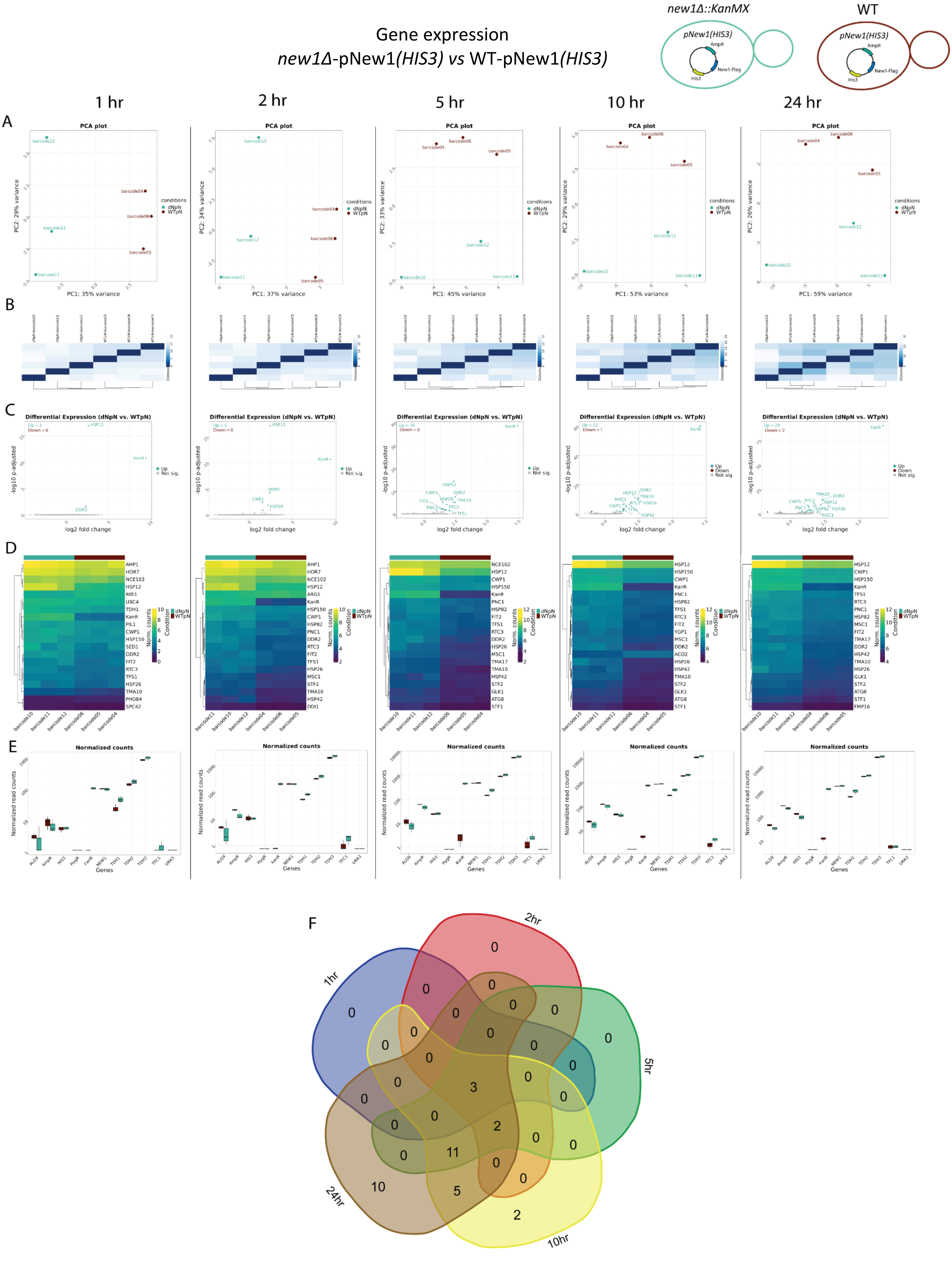
Gene expression
new1Δ-pNew1(HIS3) vs WT-pNew1(HIS3)
new1Δ::KanMX
WT
pNew1(HIS3)
AmpR
New1-Flag
His3
1 hr
2 hr
5 hr
10 hr
24 hr
A
B
C
D
E
F
PCA plot
Differential Expression (dNpN vs. WTpN)
log2 fold change
-log10 p-adjusted
Normalized counts
Normalized read counts
Genes
conditions
dNpN
WTpN
1hr
2hr
5hr
10hr
24hr

**Supplementary Figure 40. Gene expression in *new1Δ*-pNew1*(HIS3) vs* WT-pNew1*(HIS3)*. A.** Principal Component Analysis (PCA) of the top 500 genes with the highest variance across all samples, the two Principal Components. Each dot corresponds to one sample and is colored by the respective condition. **B.** Sample2sample distance plot. The euclidean distance between the gene expression patterns of all samples to each other is plotted using heatmap. Normalized gene counts were used for distance computation. **C.** Volcano plots. The differential gene expression analysis results from DESeq2 are shown by plotting the log2FoldChange between the conditions of interest against the –log10 adjusted p-value per gene observed from the Wald-Test integrated in DESeq2 [Love, Huber, and Anders et al. 2014]. The top 10 significant genes are labeled by their symbol. **D.** Heatmaps. The expression of the top 20 differentially expressed genes is plotted using Heatmap by coloring the number of reads per gene. The legend on the right side describes which colors correspond to highly and lowly expressed genes. **E**. Normalized gene count for selected genes. Normalized gene counts from FeatureCounts [Liao et al. 2014] are visualized for selected genes per condition using boxplots. The median-of-ratios normalization method from DESeq2 [Love et al. 2014] was used for normalization. **F**. 5-way Venn diagram showing the differentially expressed gene overlaps between the distinct collected time points.

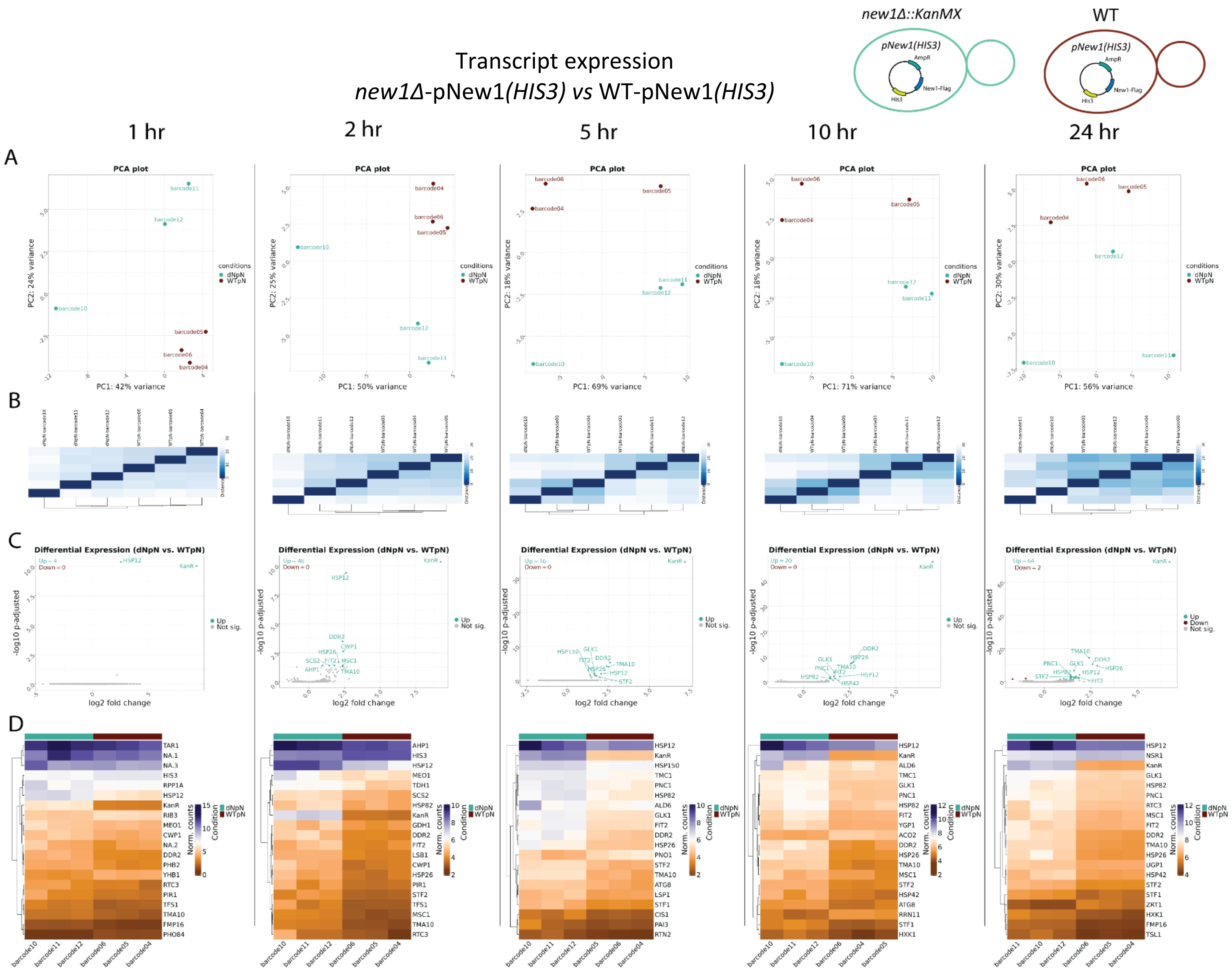


**Supplementary Figure 41. Transcript expression in *new1Δ*-pNew1*(HIS3) vs* WT-pNew1*(HIS3)*. A.** Principal Component Analysis (PCA) of the top 500 transcripts with the highest variance across all samples, the two Principal Components. Each dot corresponds to one sample and is colored by the respective condition. **B.** Sample2sample plot. The euclidean distance between the transcript expression patterns of all samples to each other is plotted using heatmap. Normalized gene counts were used for distance computation. **C.** Volcano plots. The differential transcript expression analysis results from DESeq2 are shown by plotting the log2FoldChange between the conditions of interest against the –log10 adjusted p-value per gene observed from the Wald-Test integrated in DESeq2. The top 10 significant genes are labeled by their symbol. **D.** Heatmaps. The expression of the top 20 differentially expressed transcripts is plotted using Heatmap by coloring the number of reads per transcript. The legend on the right side describes which colors correspond to highly and lowly expressed transcripts.
