## Supplementary Document S5 for "Real-time transcriptomic profiling in distinct experimental conditions"

### Supplementary Figures 42-56, related to Figure 6

#### Yeat Setup 2


**Supplementary Figure 42|** Sequencing overview -Yeast setup 2 ..... 2
**Supplementary Figure 43|** Real-time transcriptomic analysis in yeast setup 2 samples using NanopoReaTA. ..... 4

**rkr1Δ-pEV(URA3) vs WT-pEV(HIS3)**

**Supplementary Figure 44 |** General overview *rkr1Δ*-pEV*(URA3) vs* WT-pEV*(HIS3)* ..... 6
**Supplementary Figure 45 |** Gene expression *rkr1Δ*-pEV*(URA3) vs* WT-pEV*(HIS3)* ..... 8
**Supplementary Figure 46 |** Transcript expression *rkr1Δ*-pEV*(URA3) vs WT*-pEV*(HIS3)* ..... 10

**rkr1Δ-pJlp2(URA3) vs rkr1Δ-pEV(URA3)**

**Supplementary Figure 47 |** General overview *rkr1Δ*-pJlp2*(URA3) vs rkr1Δ*-pEV*(URA3)* ..... 11
**Supplementary Figure 48 |** Gene expression *rkr1Δ*-pJlp2*(URA3) vs rkr1Δ*-pEV*(URA3)* ..... 13
**Supplementary Figure 49 |** Transcript expression *rkr1Δ*-pJlp2*(URA3) vs rkr1Δ*-pEV*(URA3)* ..... 15

**rkr1Δ jlp2Δ-pJlp2(URA3) vs rkr1Δjlp2Δ-pEV(URA3)**

**Supplementary Figure 50 |** General overview *rkr1Δ jlp2Δ*-pJlp2*(URA3) vs rkr1Δjlp2Δ*-pEV*(Ura3)* ..... 16
**Supplementary Figure 51 |** Gene expression *rkr1Δ jlp2Δ*-pJlp2*(URA3) vs rkr1Δjlp2Δ*-pEV*(URA3)* ..... 18
**Supplementary Figure 52 |** Transcript expression *rkr1Δ jlp2Δ*-pJlp2*(URA3) vs rkr1Δjlp2Δ*-pEV*(URA3)* ..... 20

**rkr1Δjlp2Δ-pEV(URA3) vs WT-pEV(HIS3)**

**Supplementary Figure 53 |** General overview *rkr1Δjlp2Δ*-pEV*(URA3) vs WT*-pEV*(HIS3)* ..... 21
**Supplementary Figure 54 |** Gene expression *rkr1Δjlp2Δ*-pEV*(URA3) vs WT*-pEV*(HIS3)* ..... 23
**Supplementary Figure 55 |** Transcript expression *rkr1Δjlp2Δ*-pEV*(URA3) vs* WT-pEV*(HIS3)* ..... 25

**Supplementary Figure 56 |** IGV coverage snapshots of selected gene loci in yeast setup 1 and 2 ..... 26


#### *General supplementary figures description*

**Sequencing overview** – Refers to sequencing metrics including total reads generated per barcode/sample, mapped reads, gene counts and transcript counts.
**General overview –** Refers to data collected from NanopoReaTA regaring the general sample/condition overview including: Read length overview (per sample and condition), gene expression variability (per sample and condition), change in gene composition (per sample and condition) and Process time.
**Gene expression** - Refers to data collected from NanopoReaTA regaring differential gene expression analysis including: Principal Component Analysis (PCA), Volcano plots, Sample-to-sample variaiblity and Heatmaps.
**Transcript expression** - Refers to data collected from NanopoReaTA regaring differential transcript expression analysis including: Principal Component Analysis (PCA), Volcano plot, Sample-to-sample variaiblity and Heatmaps.

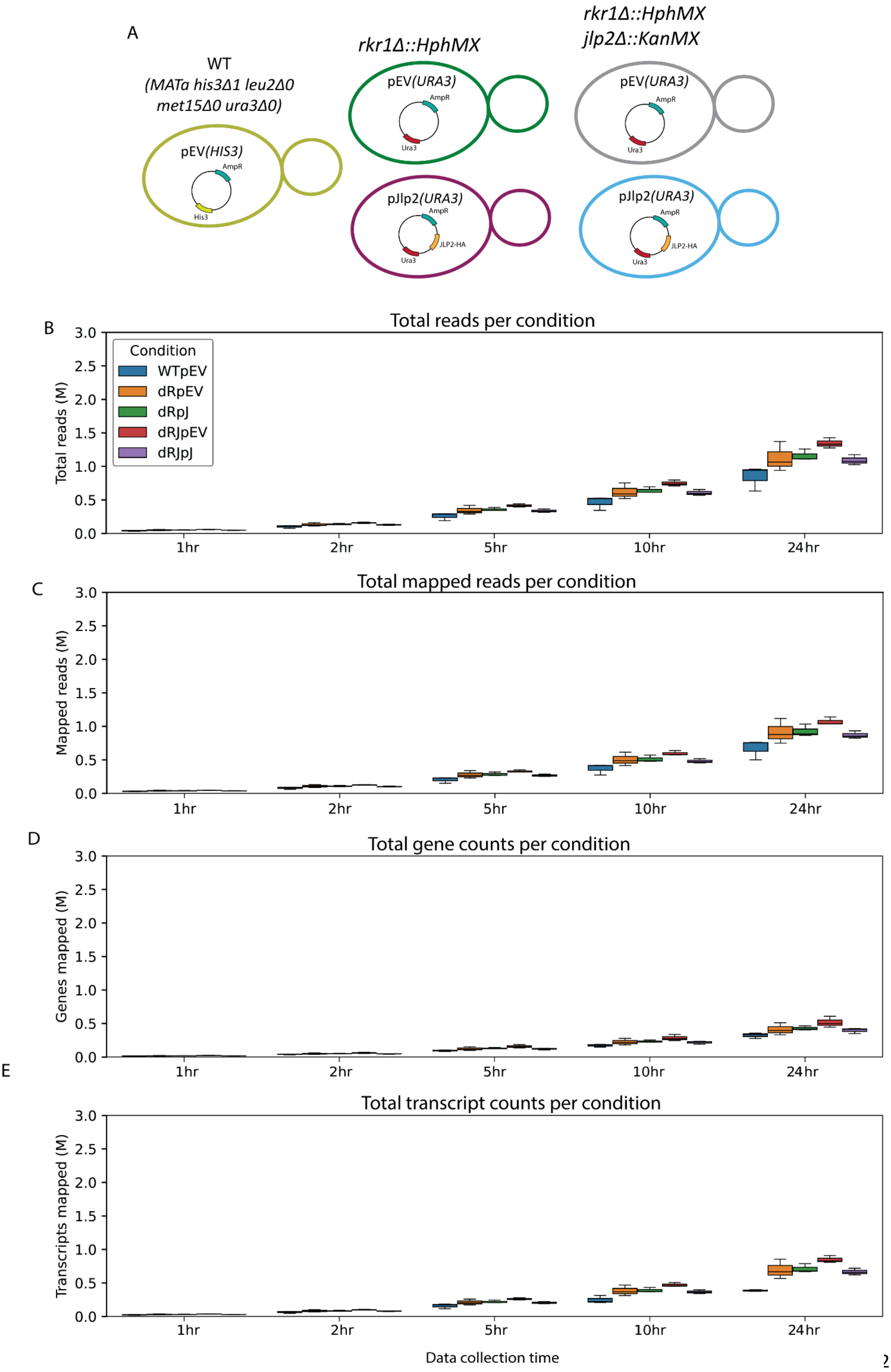
A
WT
(MATa his3Δ1 leu2Δ0 met15Δ0 ura3Δ0)
pEV(HIS3)
AmpR
His3
rkr1Δ::HphMX
pEV(URA3)
AmpR
Ura3
pJlp2(URA3)
AmpR
JLP2-HA
Ura3
rkr1Δ::HphMX
jlp2Δ::KanMX
pEV(URA3)
AmpR
Ura3
pJlp2(URA3)
AmpR
JLP2-HA
Ura3
B
Total reads per condition
Total reads (M)
Condition
WTpEV
dRpEV
dRpJ
dRJpEV
dRJpJ
0.0
0.5
1.0
1.5
2.0
2.5
3.0
1hr
2hr
5hr
10hr
24hr
C
Total mapped reads per condition
Mapped reads (M)
D
Total gene counts per condition
Genes mapped (M)
E
Total transcript counts per condition
Transcripts mapped (M)
Data collection time

**Supplementary Figure 42. Sequencing Overview Yeast Setup 2. A.** For Yeast setup 2, WT strain (BY4741, *MATa, his3Δ1, leu2Δ0, met15Δ0, ura3Δ0), rkr1Δ::HphMX* strain, where the *rkr1* gene was replaced with the Hygromycin B resistance gene (*HygR*) and double KO *jlp2Δ::KanMX rkr1Δ::HphMX*, where the *jlp2* gene was replaced with *KanR*. These strains were transformed with either an empty vector with the *URA3* selection marker (pEV*(URA3)*) or an overexpression vector for C-terminally HA-tagged *jlp2* with the *URA3* selection marker (pJlp2*(URA3)*). WT was transformed with an empty vector with the His3 selection marker (pEV(HIS3)). **B.** Total read generated per condition per time point. **B-E.** The box plot represent total read generated (**B**), total mapped reads (**C**), total gene counts (**D**) and total transcript count (**E**) per condition, per time point (3 biological replicates per condition). Abbreviations: WTpEV - WTpEV(*HIS3*), dRpEV - *rkr1Δ*pEV(*URA3*)*,* dRpj - *rkr1Δ* pJlp2*(URA3),* dRjpEV - *jlp2Δrkr1Δ*pEV(*URA3*) ,dRjpj - *jlp2Δrkr1Δ*pJlp2(*URA3*).

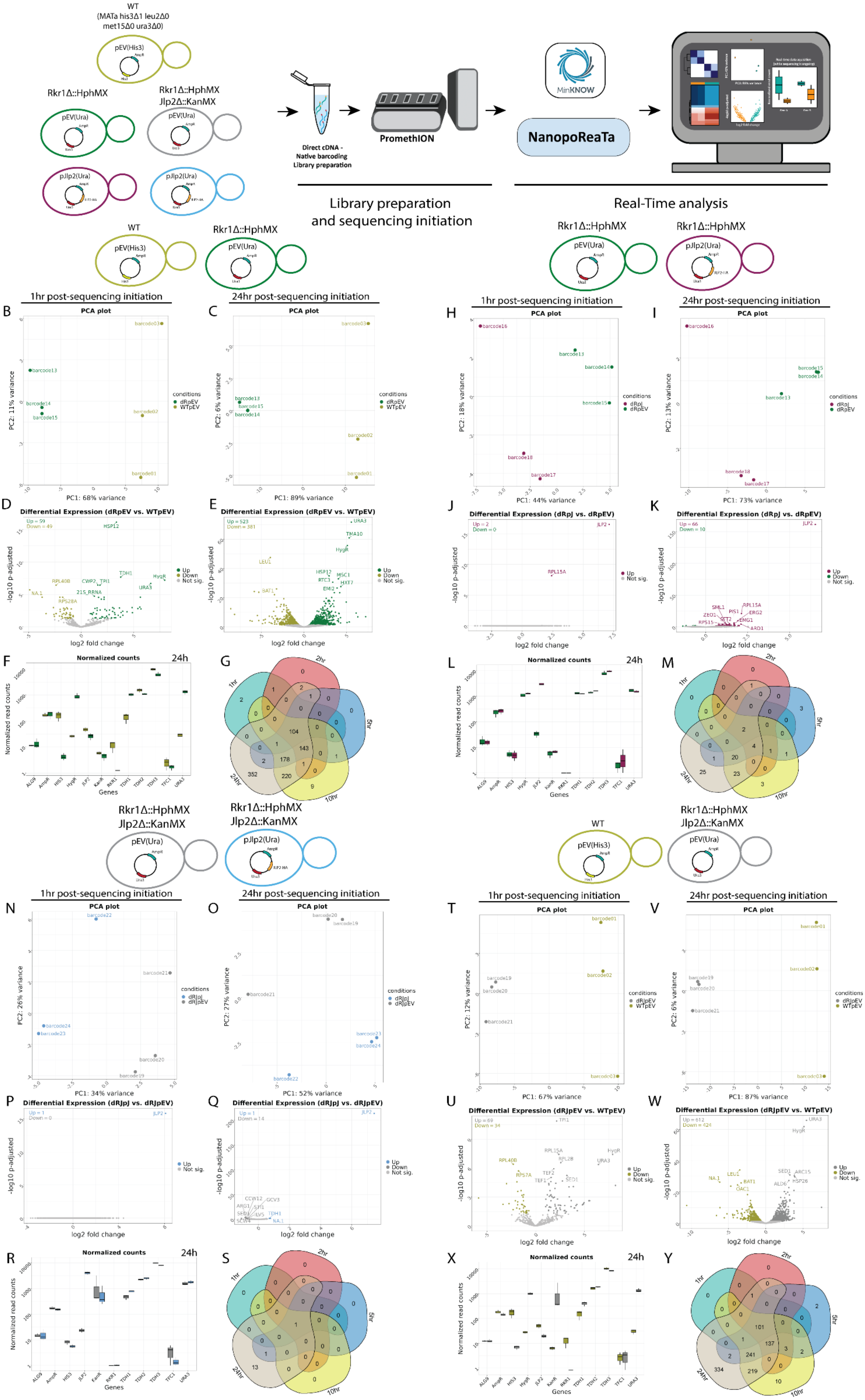
WT
(MATa his3Δ1 leu2Δ0
met15Δ0 ura3Δ0)
pEV(His3)
Rkr1Δ::HphMX
Rkr1Δ::HphMX
Jlp2Δ::KanMX
pEV(Ura)
pJlp2(Ura)
Direct cDNA -
Native barcoding
Library preparation
PromethION
MinKNOW
NanopoReaTa
Library preparation
and sequencing initiation
Real-Time analysis
1hr post-sequencing initiation
24hr post-sequencing initiation
PCA plot
Differential Expression (dRpEV vs. WTpEV)
Differential Expression (dRpJ vs. dRpEV)
Differential Expression (dRJpJ vs. dRJpEV)
Differential Expression (dRJpEV vs. WTpEV)
Normalized counts
24h
log2 fold change
-log10 p-adjusted
Normalized read counts
Genes
conditions
1hr
2hr
5hr
10hr
24hr
B
C
D
E
F
G
H
I
J
K
L
M
N
O
P
Q
R
S
T
U
V
W
X
Y

**Supplementary Figure 43. Real-time transcriptomic analysis in yeast setup 2 samples using NanopoReaTA. A.** Experimental strategy of yeast setup 2. RNA was isolated from selected yeast strains and the dscDNA library was prepared, which included sample barcoding and adapter ligation. Samples were loaded and sequenced using a PromethION R10 flow cell. NanopoReaTA was activated shortly after sequencing initiation and data was collected 1hr, 2hr, 5hr, 10hr, and 24hr post-sequencing initiation. **B-G.** Differential gene expression in *rkr1Δ*-pEV(*URA3*) versus WT-pEV(*HIS3*). Selected data plots showing PCA (**B-C**) and volcano plots (**D-E**) 1hr and 24hr post sequencing initiation. **F.** Normalized gene counts for selected genes 24hr post-sequencing initiation. Normalized gene counts are visualized for selected genes per condition using boxplots. The median-of-ratios normalization method from DESeq2 was used for normalization. **G.** Five-way Venn diagram showing the differentially expressed gene overlaps between the distinct collected time points. **H-M.** Differential gene expression in *rkr1Δ*-pJlp2(*URA3*) versus *rkr1Δ*-pEV(*URA3*). Similar analyses to B-G were conducted for PCA (**H-I**) and volcano plots (**J-K**) as well as normalized gene counts (**L**) and Venn diagram (**M**). **N-S.** Differential gene expression in *rkr1Δ, jlp2Δ*-pJlp2(*URA3*) versus *rkr1Δ, jlp2Δ*-pEV(*URA3*).Similar analyses to B-G were conducted for PCA (**N-O**) and volcano plots (**P-Q**) as well as normalized gene counts (**R**) and Venn diagram (**S**). **T-Y.** Differential gene expression in in *rkr1Δ, jlp2Δ*-pEV(*URA3*) versus WT-pEV(*HIS3*). Similar analyses to B-G were conducted for PCA (**T-V**) and volcano plots (**U-W**) as well as normalized gene counts (**X**) and Venn diagram (**Y**).

General overview
*rkr1Δ*-pEV(*URA3*) *vs* WT-pEV(*HIS3*)

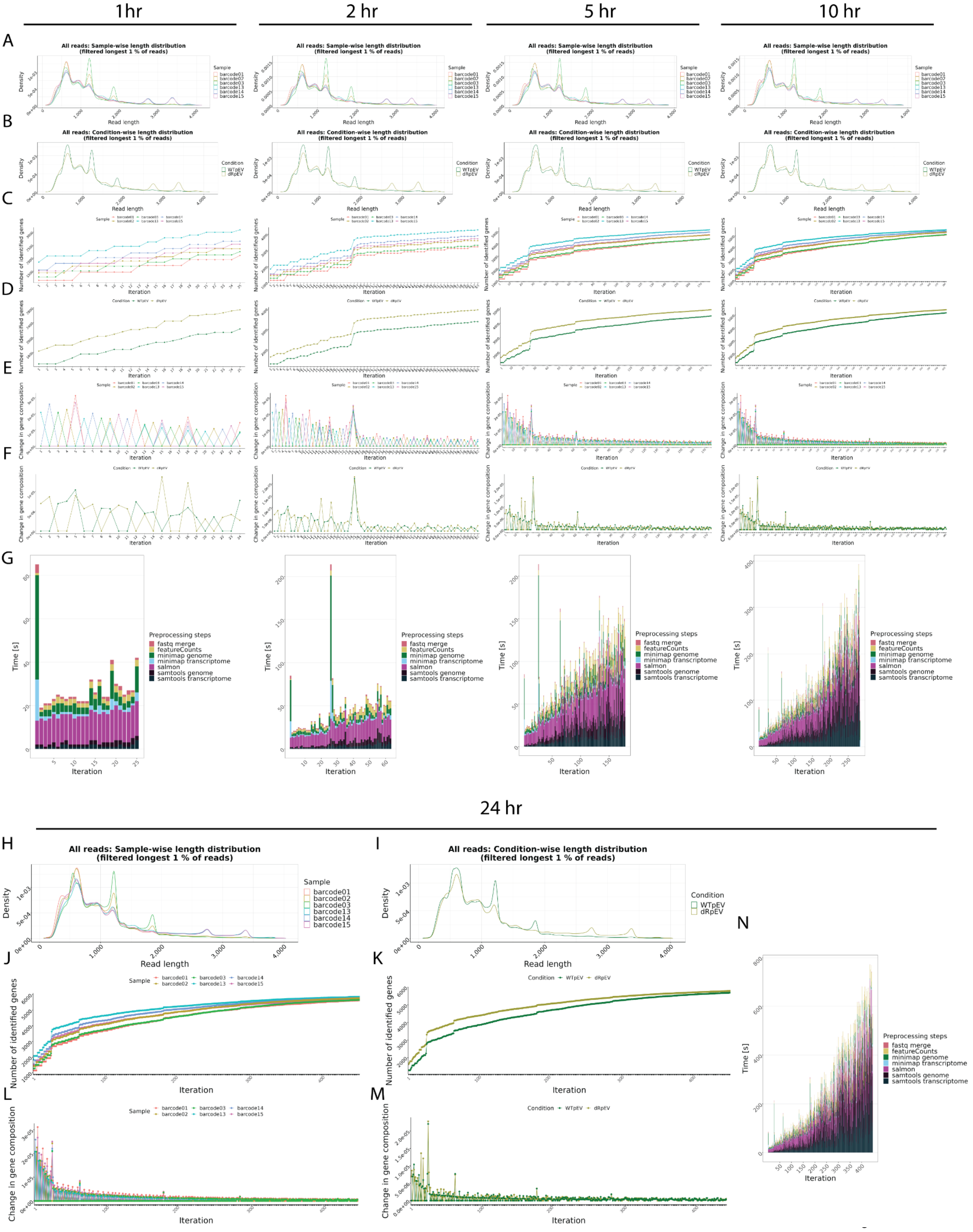

**Supplementary Figure 44. General overview in *rkr1Δ*-pEV*(URA3) vs* WT-pEV*(HIS3)*. A-B**. Read length overview. The distribution of read lengths derived from generated fastq files is plotted per sample (A) and per condition (B). All reads of length over the 99 % quantile of all lengths are removed from these visualizations. **C-D**. Gene expression variability. The number of identified genes (> 0 reads counted) is plotted after each iteration per sample (C) and per condition (D). When the detection of additional genes finishes, the lines reach a plateau. **E-F**. Change in gene composition. The change in gene composition (CGC) plot shows the sum of absolute differences of relative gene (%gi) counts per individual gene (g) in iteration (i) to the relative gene counts in iteration (i-1) for each respective sample (E) and condition (F). CGC = sum(|(%g(i-1))-(%g(i))|). **G.** Process time. The bar plots show the time in seconds all preprocessing steps needed per iteration. One iteration process at maximum 30 files from all samples. This plot updates automatically as a new process finish. **H-N**. Final general overview. Read length (H-I), Gene expression variability (J-K), Change in gene composition (L-M), and processing time (N) for the 24hr time point. All plots are organized according to their respective time points of collection.

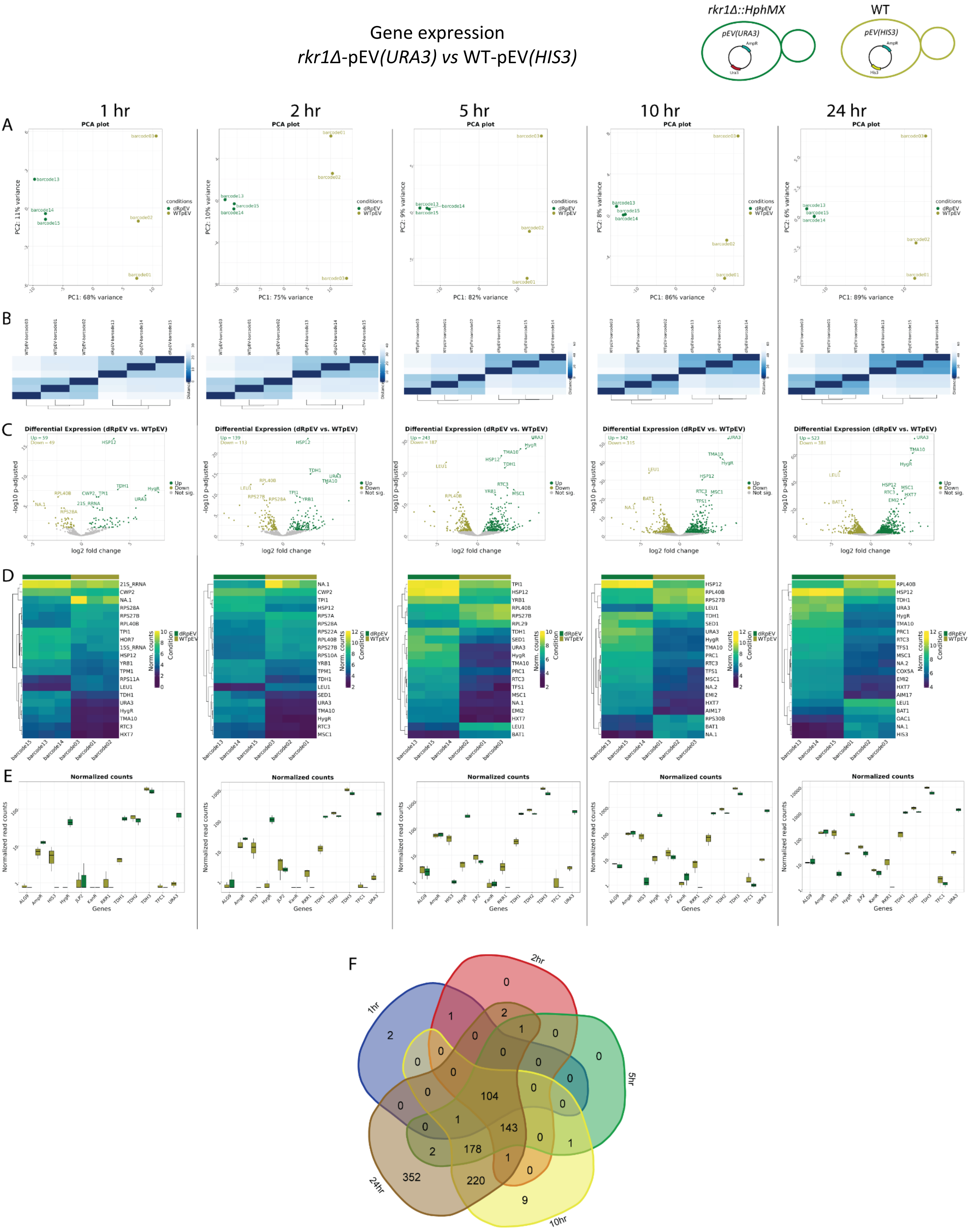
Gene expression
rkr1Δ-pEV(URA3) vs WT-pEV(HIS3)
rkr1Δ::HphMX
WT
1 hr
2 hr
5 hr
10 hr
24 hr
A
PCA plot
B
C
Differential Expression (dRpEV vs. WTpEV)
D
E
Normalized counts
F
1hr
2hr
5hr
10hr
24hr

**Supplementary Figure 45. Gene expression in *rkr1Δ*-pEV*(URA3) vs* WT-pEV*(HIS3)*. A.** Principal Component Analysis (PCA) of the top 500 genes with the highest variance across all samples, the two Principal Components. Each dot corresponds to one sample and is colored by the respective condition. **B.** Sample2sample distance plot.The euclidean distance between the gene expression patterns of all samples to each other is plotted using heatmap. Normalized gene counts were used for distance computation. **C.** Volcano plots. The differential gene expression analysis results from DESeq2 are shown by plotting the log2FoldChange between the conditions of interest against the –log10 adjusted p-value per gene observed from the Wald-Test integrated in DESeq2 (Love, Huber, and Anders et al. 2014). The top 10 significant genes are labeled by their symbol. **D.** Heatmaps. The expression of the top 20 differentially expressed genes is plotted using Heatmap by coloring the number of reads per gene. The legend on the right side describes which colors correspond to highly and lowly expressed genes. **E**. Normalized gene count for selected genes. Normalized gene counts from FeatureCounts (Liao et al. 2014) are visualized for selected genes per condition using boxplots. The median-of-ratios normalization method from DESeq2 (Love et al. 2014) was used for normalization. **F**. 5-way Venn diagram showing the differentially expressed gene overlaps between the distinct collected time points.

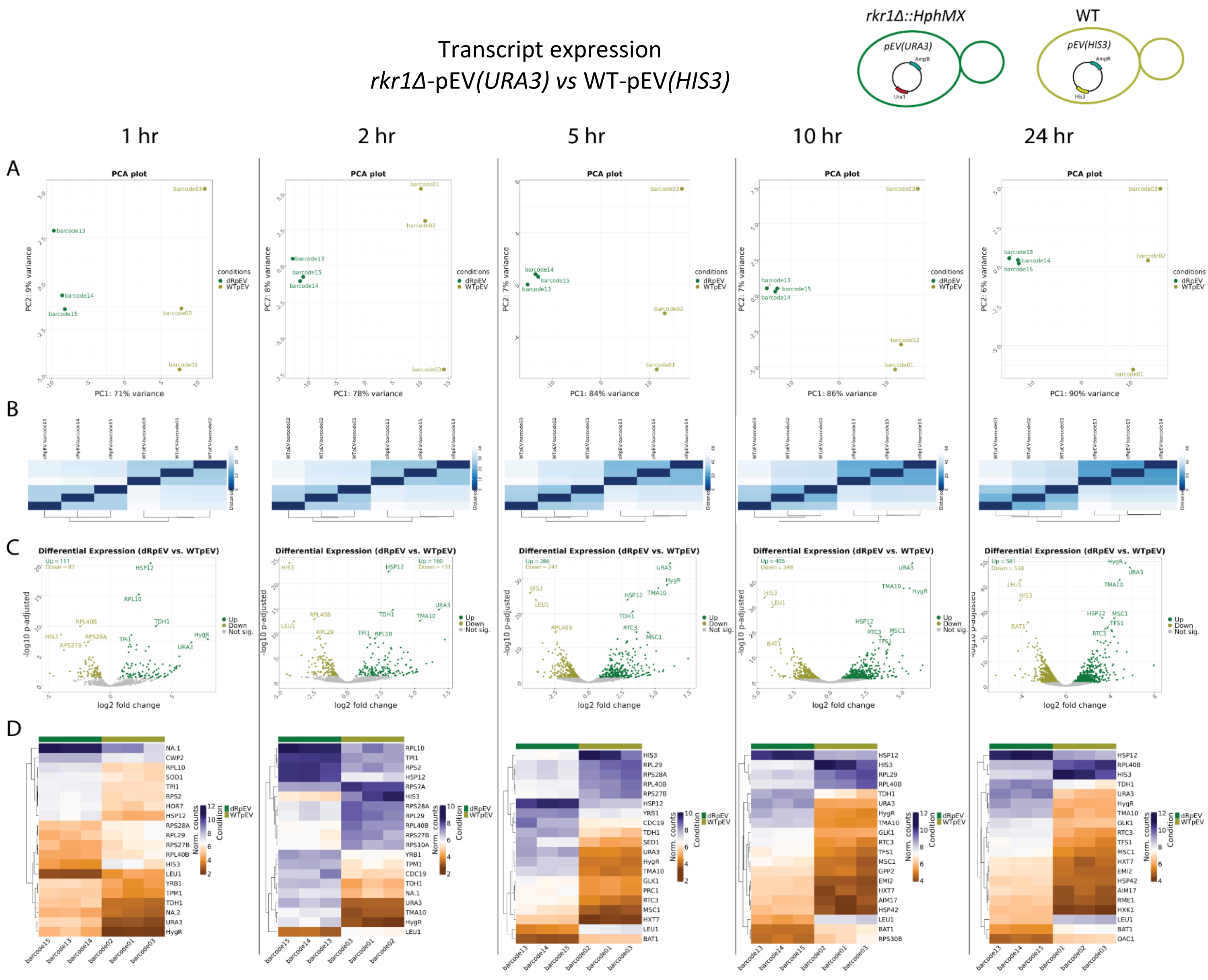


**Supplementary Figure 46. Transcript expression in *rkr1Δ*-pEV*(URA3) vs WT*-pEV*(HIS3)*. A.** Principal Component Analysis (PCA) of the top 500 transcripts with the highest variance across all samples, the two Principal Components. Each dot corresponds to one sample and is colored by the respective condition. **B.** Sample2sample plot. The euclidean distance between the transcript expression patterns of all samples to each other is plotted using heatmap. Normalized gene counts were used for distance computation. **C.** Volcano plots. The differential transcript expression analysis results from DESeq2 are shown by plotting the log2FoldChange between the conditions of interest against the –log10 adjusted p-value per gene observed from the Wald-Test integrated in DESeq2. The top 10 significant genes are labeled by their symbol. **D.** Heatmaps. The expression of the top 20 differentially expressed transcripts is plotted using Heatmap by coloring the number of reads per transcript. The legend on the right side describes which colors correspond to highly and lowly expressed transcripts.

### General overview

*rkr1Δ*-pJlp2*(URA3) vs rkr1Δ*-pEV*(URA3)*

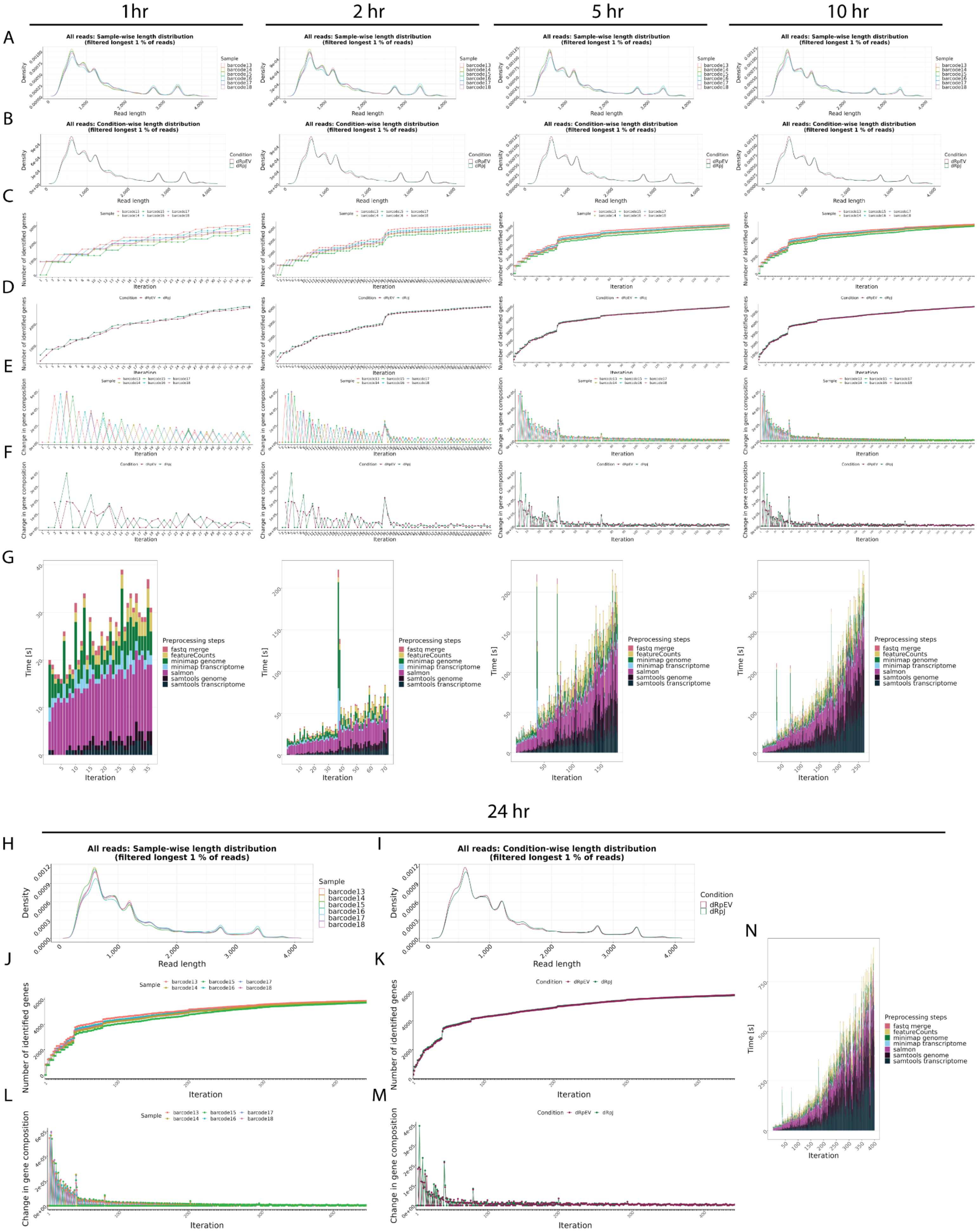

**Supplementary Figure 47. General overview in *rkr1Δ*-pJlp2*(URA3) vs rkr1Δ*-pEV*(URA3)*. A-B**. Read length overview. The distribution of read lengths derived from generated fastq files is plotted per sample (A) and per condition (B). All reads of length over the 99 % quantile of all lengths are removed from these visualizations. **C-D**. Gene expression variability. The number of identified genes (> 0 reads counted) is plotted after each iteration per sample (C) and per condition (D). When the detection of additional genes finishes, the lines reach a plateau. **E-F**. Change in gene composition. The change in gene composition (CGC) plot shows the sum of absolute differences of relative gene (%gi) counts per individual gene (g) in iteration (i) to the relative gene counts in iteration (i-1) for each respective sample (E) and condition (F). CGC = sum(|(%g(i-1))-(%g(i))|). **G.** Process time. The bar plots show the time in seconds all preprocessing steps needed per iteration. One iteration process at maximum 30 files from all samples. This plot updates automatically as a new process finish. **H-N**. Final general overview. Read length (H-I), Gene expression variability (J-K), Change in gene composition (L-M), and processing time (N) for the 24hr time point. All plots are organized according to their respective time points of collection.

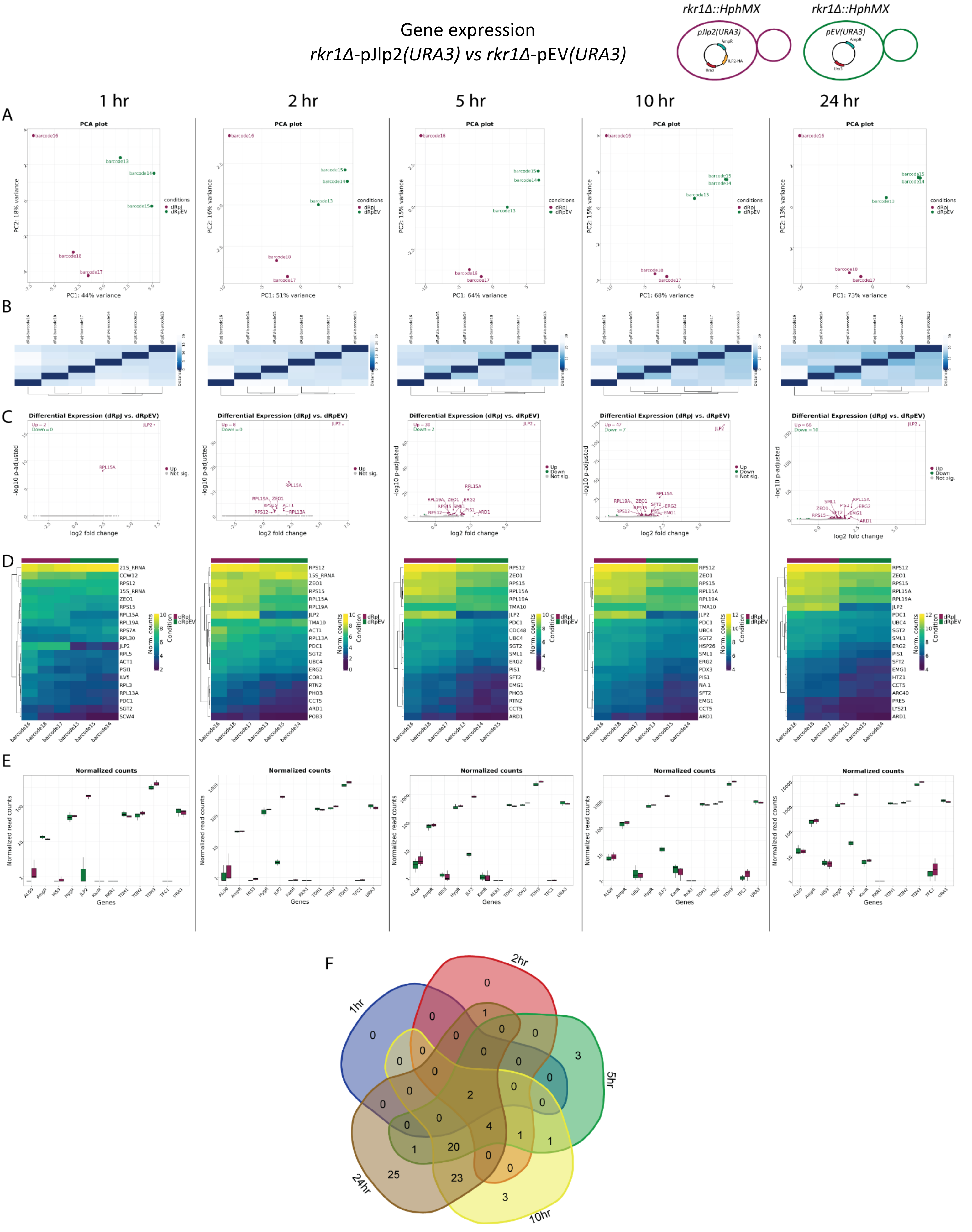
Gene expression
rkr1Δ-pJlp2(URA3) vs rkr1Δ-pEV(URA3)
rkr1Δ::HphMX
pJlp2(URA3)
rkr1Δ::HphMX
pEV(URA3)
1 hr
2 hr
5 hr
10 hr
24 hr
A
B
C
D
E
F
PCA plot
PC1: 44% variance
PC1: 51% variance
PC1: 64% variance
PC1: 68% variance
PC1: 73% variance
Differential Expression (dRpJ vs. dRpEV)
log2 fold change
-log10 p-adjusted
Normalized counts
Normalized read counts
Genes
1hr
2hr
5hr
10hr
24hr

**Supplementary Figure 48. Gene expression in *rkr1Δ*-pJlp2*(URA3) vs rkr1Δ*-pEV*(URA3)*. A.** Principal Component Analysis (PCA) of the top 500 genes with the highest variance across all samples, the two Principal Components. Each dot corresponds to one sample and is colored by the respective condition. **B.** Sample2sample distance plot.The euclidean distance between the gene expression patterns of all samples to each other is plotted using heatmap. Normalized gene counts were used for distance computation. **C.** Volcano plots. The differential gene expression analysis results from DESeq2 are shown by plotting the log2FoldChange between the conditions of interest against the –log10 adjusted p-value per gene observed from the Wald-Test integrated in DESeq2 (Love, Huber, and Anders et al. 2014). The top 10 significant genes are labeled by their symbol. **D.** Heatmaps. The expression of the top 20 differentially expressed genes is plotted using Heatmap by coloring the number of reads per gene. The legend on the right side describes which colors correspond to highly and lowly expressed genes. **E**. Normalized gene count for selected genes. Normalized gene counts from FeatureCounts (Liao et al. 2014) are visualized for selected genes per condition using boxplots. The median-of-ratios normalization method from DESeq2 (Love et al. 2014) was used for normalization. **F**. 5-way Venn diagram showing the differentially expressed gene overlaps between the distinct collected time points.

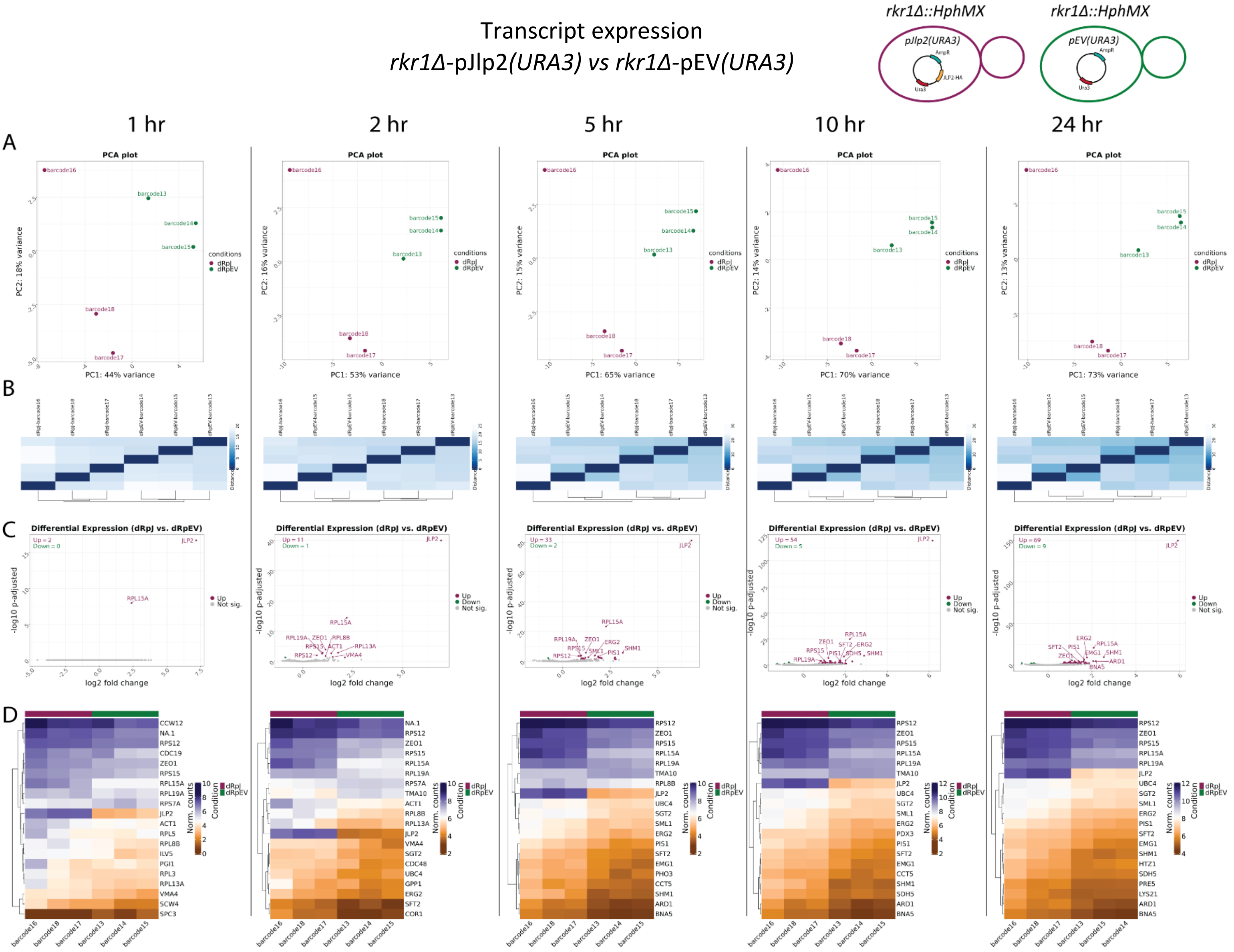


**Supplementary Figure 49. Transcript expression in *rkr1Δ*-pJlp2*(URA3)* vs *rkr1Δ*-pEV*(URA3)*. A.** Principal Component Analysis (PCA) of the top 500 transcripts with the highest variance across all samples, the two Principal Components. Each dot corresponds to one sample and is colored by the respective condition. **B.** Sample2sample plot. The euclidean distance between the transcript expression patterns of all samples to each other is plotted using heatmap. Normalized gene counts were used for distance computation. **C.** Volcano plots. The differential transcript expression analysis results from DESeq2 are shown by plotting the log2FoldChange between the conditions of interest against the –log10 adjusted p-value per gene observed from the Wald-Test integrated in DESeq2. The top 10 significant genes are labeled by their symbol. **D.** Heatmaps. The expression of the top 20 differentially expressed transcripts is plotted using Heatmap by coloring the number of reads per transcript. The legend on the right side describes which colors correspond to highly and lowly expressed transcripts.

General overview

*rkr1Δ jlp2Δ*-pJlp2*(URA3)* vs *rkr1Δ jlp2Δ*-pEV*(URA3)*

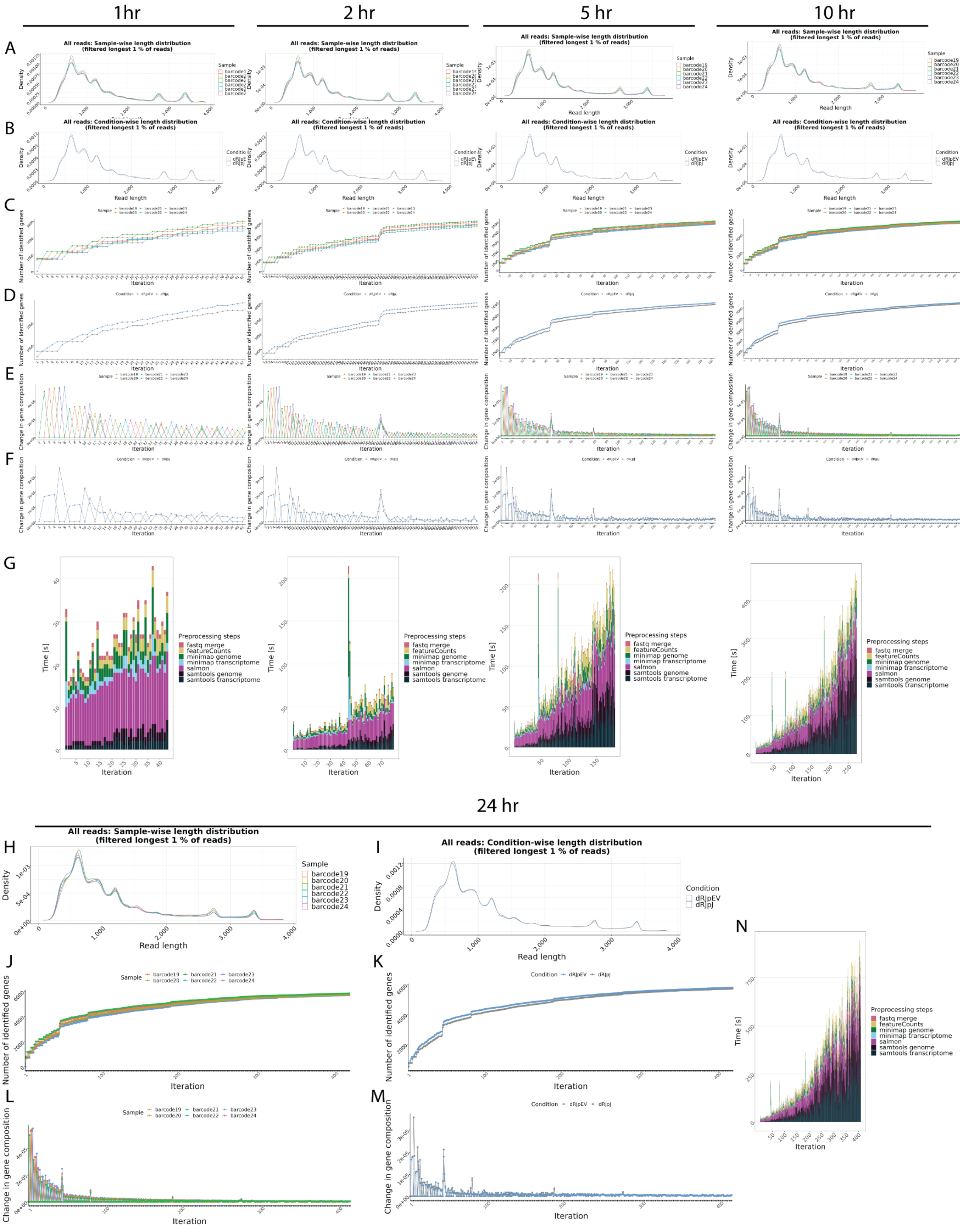

**Supplementary Figure 50. General overview in *rkr1Δjlp2Δ*-pJlp2*(URA3) vs rkr1Δjlp2Δ*-pEV*(URA3)*. A-B**. Read length overview. The distribution of read lengths derived from generated fastq files is plotted per sample (A) and per condition (B). All reads of length over the 99 % quantile of all lengths are removed from these visualizations. **C-D**. Gene expression variability. The number of identified genes (> 0 reads counted) is plotted after each iteration per sample (C) and per condition (D). When the detection of additional genes finishes, the lines reach a plateau. **E-F**. Change in gene composition. The change in gene composition (CGC) plot shows the sum of absolute differences of relative gene (%gi) counts per individual gene (g) in iteration (i) to the relative gene counts in iteration (i-1) for each respective sample (E) and condition (F). CGC = sum(|(%g(i-1))-(%g(i))|). **G.** Process time. The bar plots show the time in seconds all preprocessing steps needed per iteration. One iteration process at maximum 30 files from all samples. This plot updates automatically as a new process finish. **H-N**. Final general overview. Read length (H-I), Gene expression variability (J-K), Change in gene composition (L-M), and processing time (N) for the 24hr time point. All plots are organized according to their respective time points of collection.

#### Gene expression
#### *rkr1Δ jlp2Δ*-pJlp2*(URA3)* vs *rkr1Δ jlp2Δ*-pEV*(URA3)*

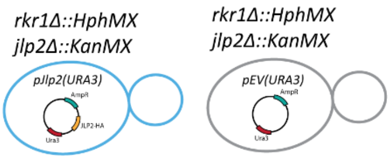


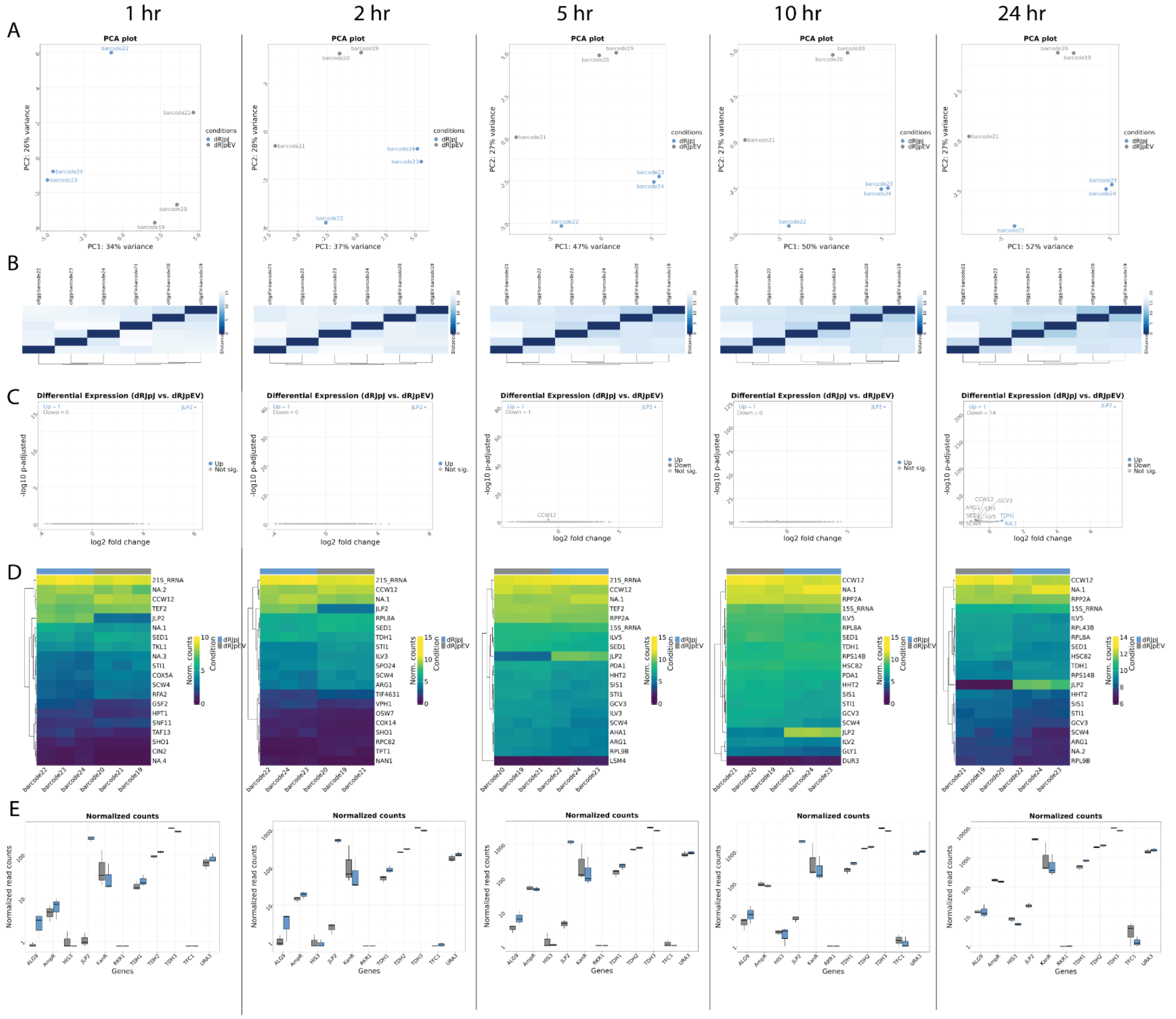


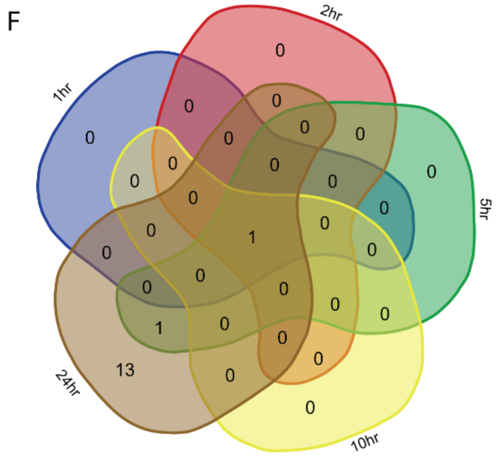

**Supplementary Figure 51. Gene expression in *rkr1Δjlp2Δ*-pJlp2*(URA3) vs rkr1Δjlp2Δ*-pEV*(URA3)*. A.** Principal Component Analysis (PCA) of the top 500 genes with the highest variance across all samples, the two Principal Components. Each dot corresponds to one sample and is colored by the respective condition. **B.** Sample2sample distance plot. The euclidean distance between the gene expression patterns of all samples to each other is plotted using heatmap. Normalized gene counts were used for distance computation. **C.** Volcano plots. The differential gene expression analysis results from DESeq2 are shown by plotting the log2FoldChange between the conditions of interest against the –log10 adjusted p-value per gene observed from the Wald-Test integrated in DESeq2 (Love, Huber, and Anders et al. 2014). The top 10 significant genes are labeled by their symbol. **D.** Heatmaps. The expression of the top 20 differentially expressed genes is plotted using Heatmap by coloring the number of reads per gene. The legend on the right side describes which colors correspond to highly and lowly expressed genes. **E**. Normalized gene count for selected genes. Normalized gene counts from FeatureCounts (Liao et al. 2014) are visualized for selected genes per condition using boxplots. The median-of-ratios normalization method from DESeq2 (Love et al. 2014) was used for normalization. **F**. 5-way Venn diagram showing the differentially expressed gene overlaps between the distinct collected time points.

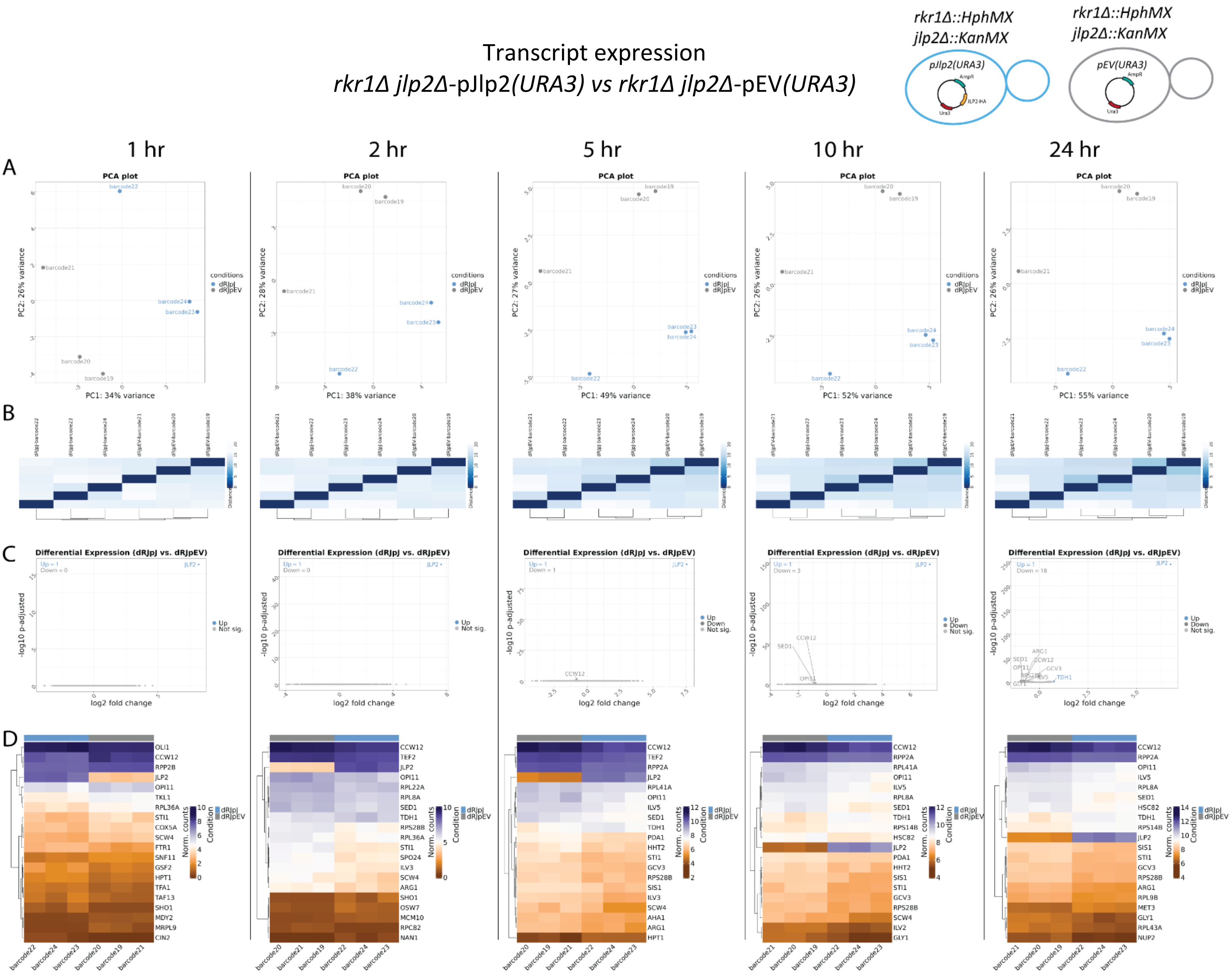


**Supplementary Figure 52. Transcript expression in *rkr1Δjlp2Δ-pJlp2(URA3) vs rkr1Δjlp2Δ-pEV(URA3)*. A.** Principal Component Analysis (PCA) of the top 500 transcripts with the highest variance across all samples, the two Principal Components. Each dot corresponds to one sample and is colored by the respective condition. **B.** Sample2sample plot. The euclidean distance between the transcript expression patterns of all samples to each other is plotted using heatmap. Normalized gene counts were used for distance computation. **C.** Volcano plots. The differential transcript expression analysis results from DESeq2 are shown by plotting the log2FoldChange between the conditions of interest against the –log10 adjusted p-value per gene observed from the Wald-Test integrated in DESeq2. The top 10 significant genes are labeled by their symbol. **D.** Heatmaps. The expression of the top 20 differentially expressed transcripts is plotted using Heatmap by coloring the number of reads per transcript. The legend on the right side describes which colors correspond to highly and lowly expressed transcripts.

General overview

*rkr1Δ jlp2Δ*-pEV(*URA3*) *vs* WT-pEV(*HIS3*)

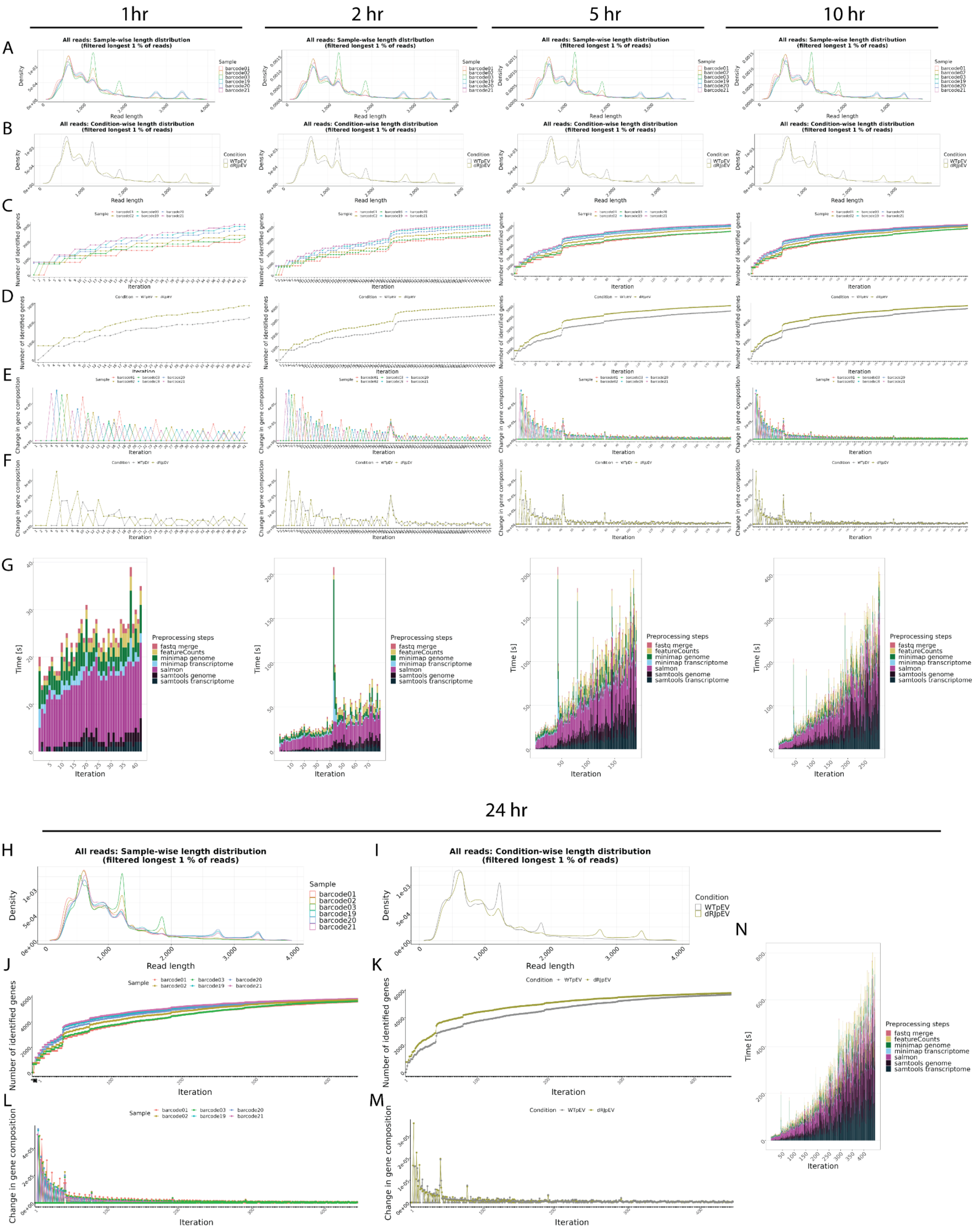

**Supplementary Figure 53. General overview in *rkr1Δjlp2Δ*-pEV*(URA3) vs WT*-pEV*(HIS3)*. A-B**. Read length overview. The distribution of read lengths derived from generated fastq files is plotted per sample (A) and per condition (B). All reads of length over the 99 % quantile of all lengths are removed from these visualizations. **C-D**. Gene expression variability. The number of identified genes (> 0 reads counted) is plotted after each iteration per sample (C) and per condition (D). When the detection of additional genes finishes, the lines reach a plateau. **E-F**. Change in gene composition. The change in gene composition (CGC) plot shows the sum of absolute differences of relative gene (%gi) counts per individual gene (g) in iteration (i) to the relative gene counts in iteration (i-1) for each respective sample (E) and condition (F). CGC = sum(|(%g(i-1))-(%g(i))|). **G.** Process time. The bar plots show the time in seconds all preprocessing steps needed per iteration. One iteration process at maximum 30 files from all samples. This plot updates automatically as a new process finish. **H-N**. Final general overview. Read length (H-I), Gene expression variability (J-K), Change in gene composition (L-M), and processing time (N) for the 24hr time point. All plots are organized according to their respective time points of collection.

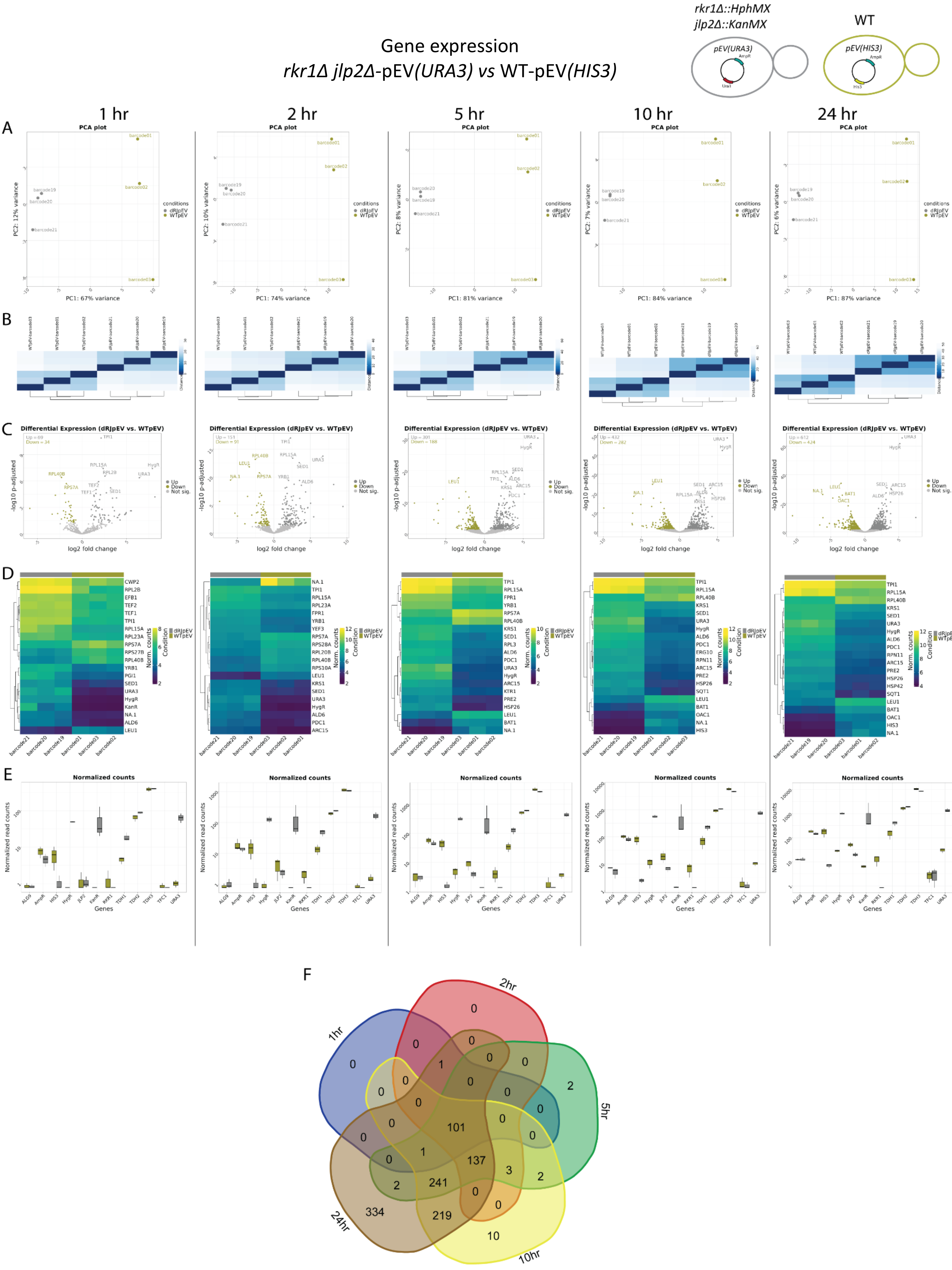
Gene expression
rkr1Δ jlp2Δ-pEV(URA3) vs WT-pEV(HIS3)
rkr1Δ::HphMX
jlp2Δ::KanMX
WT
pEV(URA3)
pEV(HIS3)
1 hr
2 hr
5 hr
10 hr
24 hr
A
B
C
D
E
F
PCA plot
PC1: 67% variance
PC2: 12% variance
PC1: 74% variance
PC2: 10% variance
PC1: 81% variance
PC2: 8% variance
PC1: 84% variance
PC2: 7% variance
PC1: 87% variance
PC2: 6% variance
conditions
dRJpEV
WTpEV
Differential Expression (dRJpEV vs. WTpEV)
log2 fold change
-log10 p-adjusted
Up
Down
Not sig.
Norm. counts
Condition
Normalized counts
Normalized read counts
Genes
1hr
2hr
5hr
10hr
24hr
101
137
241
219
334
10

**Supplementary Figure 54. Gene expression in *rkr1Δjlp2Δ*-pEV*(URA3) vs WT*-pEV*(HIS3)*. A.** Principal Component Analysis (PCA) of the top 500 genes with the highest variance across all samples, the two Principal Components. Each dot corresponds to one sample and is colored by the respective condition. **B.** Sample2sample distance plot.The euclidean distance between the gene expression patterns of all samples to each other is plotted using heatmap. Normalized gene counts were used for distance computation. **C.** Volcano plots. The differential gene expression analysis results from DESeq2 are shown by plotting the log2FoldChange between the conditions of interest against the –log10 adjusted p-value per gene observed from the Wald-Test integrated in DESeq2 (Love, Huber, and Anders et al. 2014). The top 10 significant genes are labeled by their symbol. **D.** Heatmaps. The expression of the top 20 differentially expressed genes is plotted using Heatmap by coloring the number of reads per gene. The legend on the right side describes which colors correspond to highly and lowly expressed genes. **E**. Normalized gene count for selected genes. Normalized gene counts from FeatureCounts (Liao et al. 2014) are visualized for selected genes per condition using boxplots. The median-of-ratios normalization method from DESeq2 (Love et al. 2014) was used for normalization. **F**. 5-way Venn diagram showing the differentially expressed gene overlaps between the distinct collected time points.

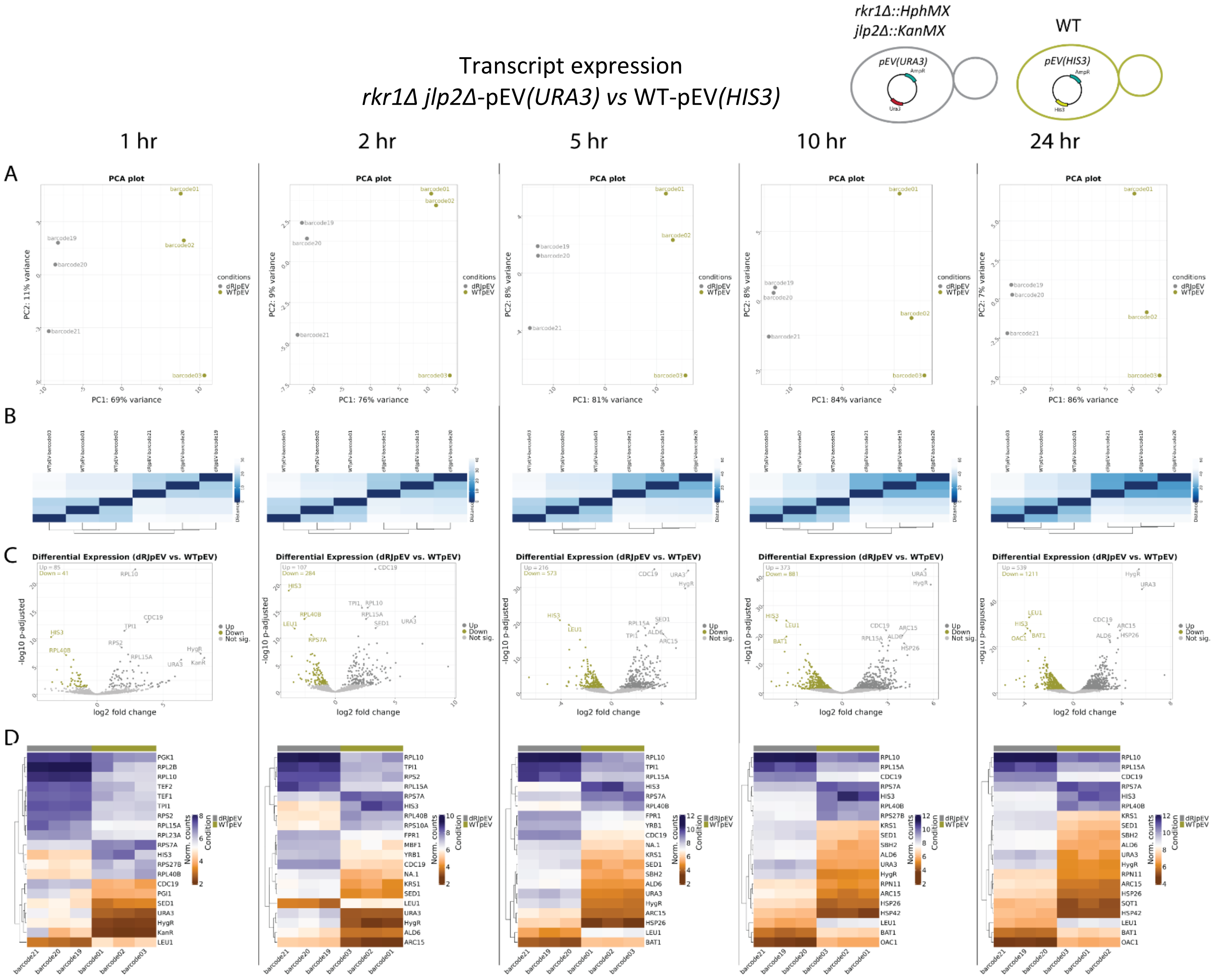


**Supplementary Figure 55. Transcript expression in *rkr1Δjlp2Δ*-pEV(*URA3*) *vs* WT-pEV(*HIS3*). A.** Principal Component Analysis (PCA) of the top 500 transcripts with the highest variance across all samples, the two Principal Components. Each dot corresponds to one sample and is colored by the respective condition. **B.** Sample2sample plot. The euclidean distance between the transcript expression patterns of all samples to each other is plotted using heatmap. Normalized gene counts were used for distance computation. **C.** Volcano plots. The differential transcript expression analysis results from DESeq2 are shown by plotting the log2FoldChange between the conditions of interest against the –log10 adjusted p-value per gene observed from the Wald-Test integrated in DESeq2. The top 10 significant genes are labeled by their symbol. **D.** Heatmaps. The expression of the top 20 differentially expressed transcripts is plotted using Heatmap by coloring the number of reads per transcript. The legend on the right side describes which colors correspond to highly and lowly expressed transcripts.

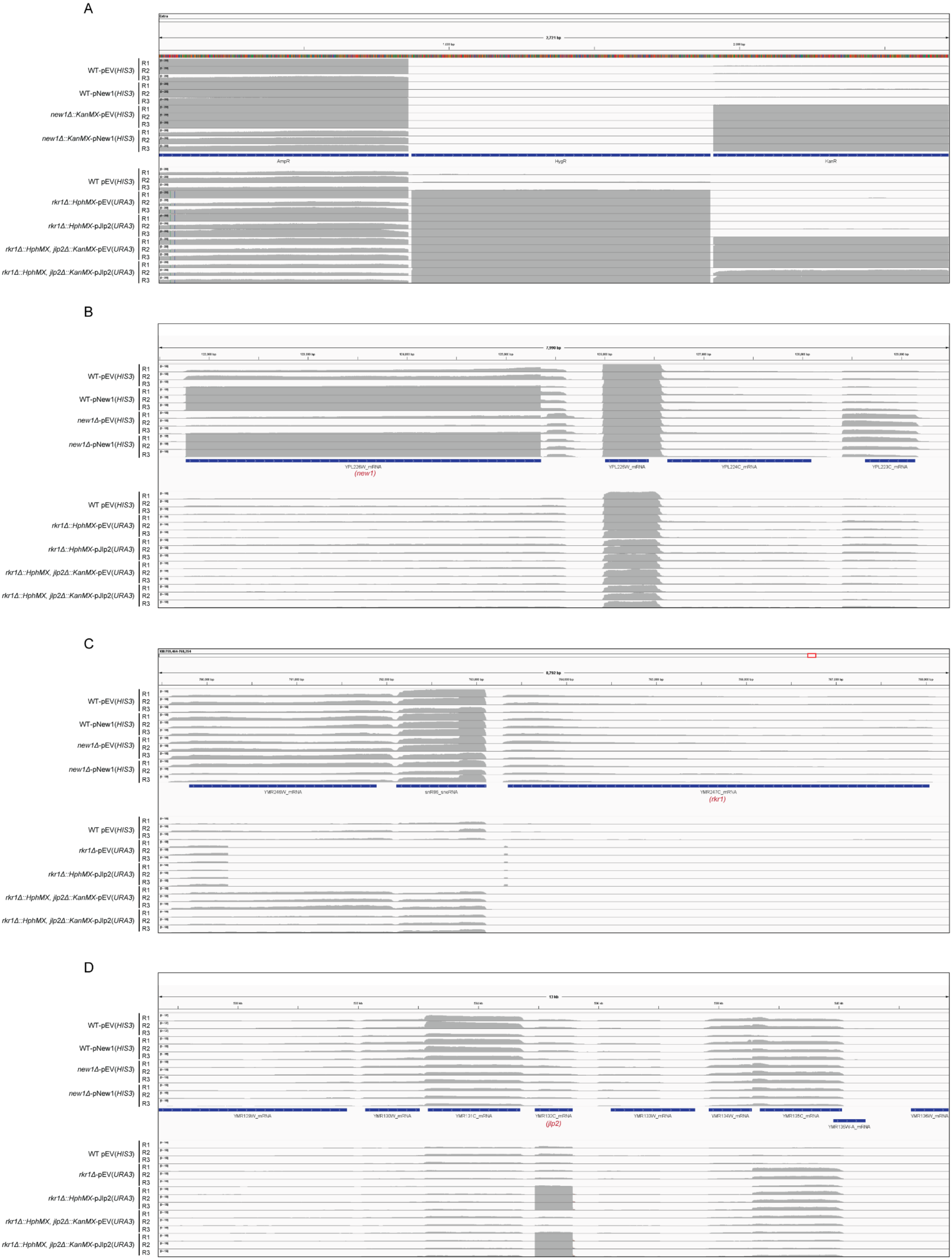
A
WT-pEV(HIS3)
WT-pNew1(HIS3)
new1Δ::KanMX-pEV(HIS3)
new1Δ::KanMX-pNew1(HIS3)
WT pEV(HIS3)
rkr1Δ::HphMX-pEV(URA3)
rkr1Δ::HphMX-pJlp2(URA3)
rkr1Δ::HphMX, jlp2Δ::KanMX-pEV(URA3)
rkr1Δ::HphMX, jlp2Δ::KanMX-pJlp2(URA3)
R1
R2
R3
2,721 bp
AmpR
HygR
KanR
B
WT-pEV(HIS3)
WT-pNew1(HIS3)
new1Δ-pEV(HIS3)
new1Δ-pNew1(HIS3)
WT pEV(HIS3)
rkr1Δ::HphMX-pEV(URA3)
rkr1Δ::HphMX-pJlp2(URA3)
rkr1Δ::HphMX, jlp2Δ::KanMX-pEV(URA3)
rkr1Δ::HphMX, jlp2Δ::KanMX-pJlp2(URA3)
7,990 bp
YPL226W_mRNA
(new1)
YPL225W_mRNA
YPL224C_mRNA
YPL223C_mRNA
C
WT-pEV(HIS3)
WT-pNew1(HIS3)
new1Δ-pEV(HIS3)
new1Δ-pNew1(HIS3)
WT pEV(HIS3)
rkr1Δ-pEV(URA3)
rkr1Δ::HphMX-pJlp2(URA3)
rkr1Δ::HphMX, jlp2Δ::KanMX-pEV(URA3)
rkr1Δ::HphMX, jlp2Δ::KanMX-pJlp2(URA3)
8,792 bp
YMR246W_mRNA
snR86_snoRNA
YMR247C_mRNA
(rkr1)
D
WT-pEV(HIS3)
WT-pNew1(HIS3)
new1Δ-pEV(HIS3)
new1Δ-pNew1(HIS3)
WT pEV(HIS3)
rkr1Δ-pEV(URA3)
rkr1Δ::HphMX-pJlp2(URA3)
rkr1Δ::HphMX, jlp2Δ::KanMX-pEV(URA3)
rkr1Δ::HphMX, jlp2Δ::KanMX-pJlp2(URA3)
13 kb
YMR129W_mRNA
YMR130W_mRNA
YMR131C_mRNA
YMR132C_mRNA
(jlp2)
YMR133W_mRNA
YMR134W_mRNA
YMR135C_mRNA
YMR135W-A_mRNA
YMR136W_mRNA

E

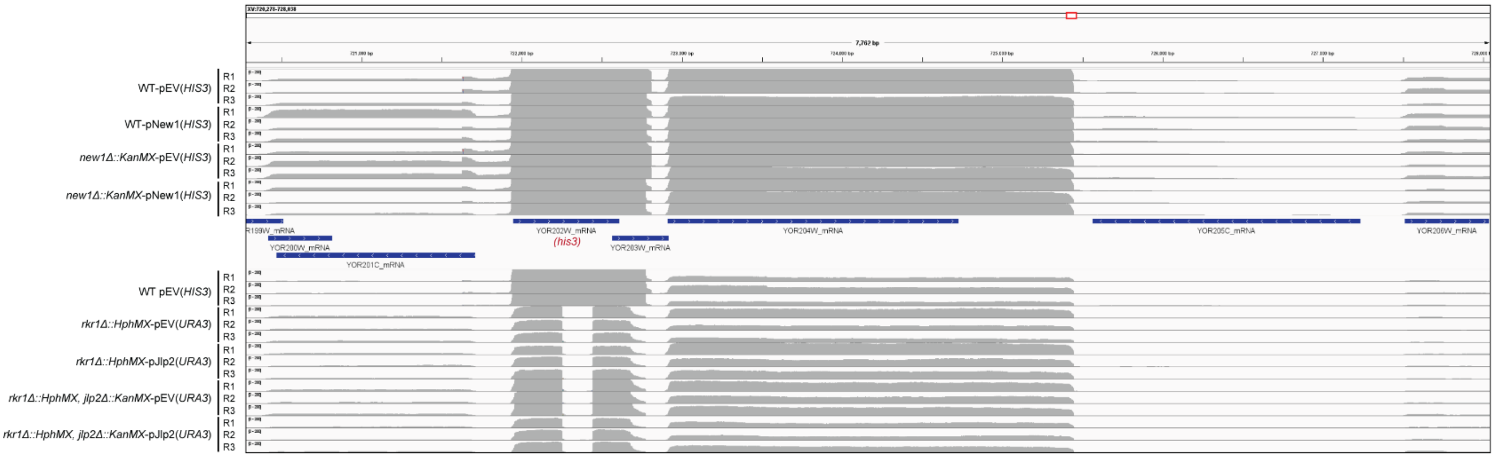


F

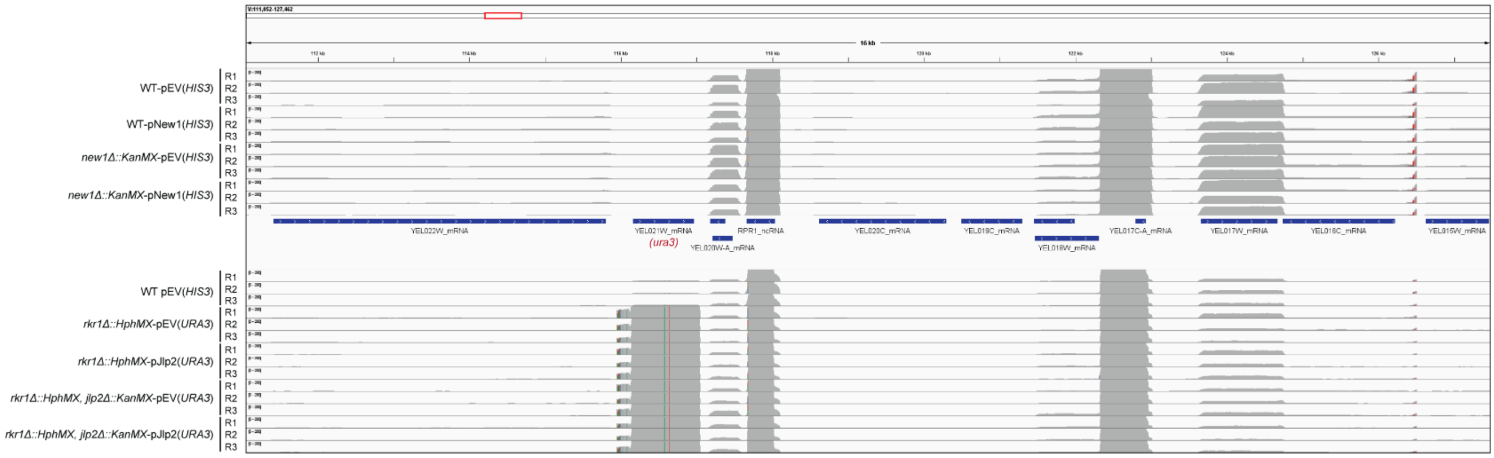


**Supplementary Figure 56. IGV coverage snapshots of selected gene loci in yeast setup 1 and 2. A-F.** Integrative Genome Viewer (IGV) snapshots of reads mapping to *AmpR*, *HygR* and *KanR* **(A)**, *NEW1* (YPL226W) (**B**), *RKR1* (YMR247C) (**C**), *JLP2* (YMR132C) (**D**), *HIS3* (YOR202W)(**E**) and *URA3* (YEL021W)(**F**).
